# ExPert infers causal perturbations from cell states across biological and clinical contexts

**DOI:** 10.64898/2026.11.02.762549

**Authors:** Leon Schwartz, Jacob L. Schlamowitz, Benjamin Kuznets-Speck, Conor W. Johnson, Diptatanu Das, Emanuelle I. Grody, Tao Bai, Hanxiao Sun, Irish Senthilkumar, Gillian Primavera, Jaekwon Jung, Britta Kuusik, Rogan A. Grant, Enrique Rojas, Abdalla M. Abdrabou, Brandon Peiffer, Zhaoli Sun, Anna Mae Diehl, Carsten Marr, Navdeep S. Chandel, Divakar S. Mithal, Samuel E. Weinberg, Mazhar Adli, Auinash Kalsotra, Ekta Prashnani, Yogesh Goyal

**Affiliations:** Department of Cell and Developmental Biology, Feinberg School of Medicine, Northwestern University, Chicago, IL, USA; Center for Synthetic Biology, McCormick School of Engineering, Northwestern University, Evanston, IL, USA; Robert H. Lurie Comprehensive Cancer Center, Feinberg School of Medicine, Northwestern University, Chicago, IL, USA; NSF-Simons National Institute for Theory and Mathematics in Biology, Chicago, IL, USA; Data Science of Systems Biology, Technical University of Munich, Freising, Germany; Department of Biochemistry, University of Illinois Urbana-Champaign, Urbana, IL, USA; Department of Obstetrics and Gynecology, Feinberg School of Medicine, Northwestern University, Chicago, IL, USA; Section of Neurology, Ann and Robert H. Lurie Children’s Hospital of Chicago, Chicago, IL, USA; Department of Pediatrics, Northwestern University Feinberg School of Medicine, Chicago, IL, USA; Division of Pulmonary and Critical Care Medicine, Department of Medicine, Feinberg School of Medicine, Northwestern University, Chicago, IL, USA; Department of Surgery, Johns Hopkins School of Medicine, Baltimore, MD, USA; Department of Medicine, Duke University School of Medicine, Durham, NC, USA; Computational Health Center & Helmholtz AI, Helmholtz Munich, Neuherberg, Germany; Munich Center for Machine Learning (MCML), Munich, Germany; Department of Biochemistry and Molecular Genetics, Feinberg School of Medicine, Northwestern University, Chicago, IL, USA; Biohub, Chicago, IL, USA; Department of Pathology, Feinberg School of Medicine, Northwestern University, Chicago, IL, USA; Carl R. Woese Institute for Genomic Biology, University of Illinois Urbana-Champaign, Urbana, IL, USA; Division of Nutritional Sciences, University of Illinois Urbana-Champaign, Urbana, IL, USA; Cancer Center at Illinois, University of Illinois Urbana-Champaign, Urbana, IL, USA; NVIDIA, Santa Clara, CA, USA; Department of Molecular Biology and Genetics, Koç University, Istanbul, Turkey; Department of Medicine III, LMU University Hospital, LMU Medizin, LMU Munich, Germany; Department of Physics, Ludwig-Maximilians-Universität München, Munich, Germany; German Cancer Consortium (DKTK), partner site Munich, Germany

**Keywords:** perturbation inference, single-cell CRISPR screens, zero-shot prediction, alcohol-associated liver disease, primary mitochondrial disease

## Abstract

Pooled single-cell CRISPR screens have driven substantial effort toward models predicting the response to a perturbation. We posed the inverse question: which perturbation produced an observed cell state? Here we developed ExPert (Extracting Perturbations), which learns one prototype per perturbation in a latent space shared by 12 cellular contexts, trained on 8,964 perturbations and 19.5 million cells, and ranks 19,895 candidate genes per query, over half absent from training. ExPert more than doubled held-out-context recall over the best existing method with 1,000-fold less compute, and transferred without retraining to drugs, protein degradation and combinations. In alcohol-associated liver disease it implicated mRNA export, unchanged in expression, and we experimentally confirmed nuclear poly(A) retention and isoform switching. In primary mitochondrial disease it recovered the affected complex from blood, localized the signal to monocytes, and flagged a misdiagnosed control. Perturbation atlases can thus be queried in reverse, from state to likely driver.

## INTRODUCTION

A cell that has drifted out of its normal state, whether in a patient biopsy or a culture dish, arrives as an endpoint whose cause is unlabeled and often inaccessible. The default for characterizing such an endpoint against an appropriate control is computing differentially expressed genes, which reflects the downstream consequences of a perturbation and not necessarily the upstream driver(s) that produced it.

Such endpoints are most often interpreted against static single-cell atlases, which now span hundreds of millions of cells across tissues, organisms, and conditions^1–3^. Models trained on them learn low-dimensional representations of cell state that support cell-type classification, reconstruct developmental hierarchies, identify disease states, and transfer across species^4–8^. In specific cases, such approaches have recovered aspects of causal structure from observational data alone, through *in silico* gene deletion and network-based simulations^6,9^. However, these models do not always outperform linear baselines in predicting measured perturbation responses^10^, a limitation attributed in part to the training data rather than model capacity. Pooled single-cell CRISPR screens break that limitation by measuring the consequences of an intervention directly, coupling a genetic perturbation to a single-cell transcriptomic readout^11–18^. They now operate at genome scale across millions of cells and several thousand targeted genes^19^, with comparable screens extending to primary human cells^20^, chemical perturbations^21,22^, cytokine stimulation^23^, and *in vivo* screens^24^. Unlike a static atlas, every state in a perturbation atlas has a known cause, so an unexplained state can be matched against it to ask which measured perturbation most likely produced it, the question we tackle in this study.

Almost all modeling built on these data has addressed the forward problem: given a perturbation, predict the resulting gene expression state. Early approaches regressed expression onto perturbation identity^13^ or factorized responses into gene programs^25,26^. More recent approaches have pursued broader predictive capabilities, including compositional autoencoders^27–29^, graph-based propagation through gene networks^30,31^, optimal-transport methods^32^, transformers pretrained on observational data^33,34^, language-model embeddings of gene function^35^, and response models trained on perturbation data at scale^36^. Although prediction accuracy has improved, it is often at the expense of linear interpretability. Others derive perturbation responses from quantities that are directly interpretable, such as gene–gene covariances in unperturbed cells^37^. Across these approaches, generalization to unseen contexts and combinations remains limited^10^, and none is designed to infer which perturbation produced a given state.

Disease presents the inverse problem, yet far less attention has been paid to it. Two recent methods attempt this problem with Perturb-seq reference data, assigning query cells to reference perturbations under explicit uncertainty^38^ or reconstructing a cell-state signature as a combination of perturbation effects^20^. Both build on the Connectivity Map and its L1000 expansion, which match a query transcriptional signature against a library of bulk perturbational profiles^39,40^. Each fits its reference within a single screen rather than across screens, and thus both report that perturbation effects transfer poorly between cellular contexts^20,38^.

Here we present ExPert (Extracting Perturbations), a single model trained across 22 single-cell CRISPR interference (CRISPRi) screens spanning 12 cellular contexts (different cell lines, stimulation conditions, and human donors), comprising 61,445 perturbation–context pairs, 8,964 unique perturbations, and 19.5 million cells. ExPert learns a shared latent representation, or “prototype,” for each perturbation across different contexts. By incorporating prior knowledge of gene function, it can also score and predict the effects of perturbing genes not tested in existing screens. ExPert more than doubled the top-10 recall of the best existing method in contexts held out from training, used three orders of magnitude less compute, and ranked 19,895 candidate gene perturbations for every query, 10,931 of them absent from its training data. ExPert generalized beyond the perturbation type it was trained on: without retraining, it outperformed other methods at recovering annotated drug targets across two small-molecule screens, recovered degraded proteins in a degron experiment we generated for this study, and recognized perturbation combinations it never trained on, including activating perturbations. Applied to two complex disease settings, ExPert recovered biology that differential expression did not. In alcohol-associated liver disease (ALD), it implicated aberrant mRNA export as a previously unrecognized disease-associated process, leading us to uncover nuclear polyadenylated transcript retention and isoform switching across the mRNA export machinery. In primary mitochondrial disease where differential expression does not identify clear signatures of mitochondrial impairment, ExPert localized respiratory chain dysfunction to monocytes and isolated patient-specific genetic impairments including in donors with uncertain diagnoses. ExPert inverts the traditional query, going from an observed state back to the perturbation that could have produced it, and extends the use of perturbation atlases from predicting responses to identifying the genetic and pharmacological knobs to tune them.

## RESULTS

### A single corpus spans cellular contexts and perturbation screens

A perturbation’s effect in one context may be different in another^20,41^. We reasoned that a model trained across diverse contexts could learn the shared perturbation component capable of generalizing to new contexts. We therefore set out to maximize the diversity of cellular contexts (cell line, stimulation condition, and donor) a single model sees rather than the depth of any one screen. We pooled 22 large-scale single-cell CRISPRi screens into one corpus (**Figure 1A**; **Figure S1**; **Table S1A**): ten screens in nine cell lines (K562, HEK293T, HCT116, KOLF2.1J, hESC, TeloHAEC, Jurkat, RPE1, and HepG2) and twelve in primary CD4⁺ T cells from four donors in three stimulation states, giving 12 contexts and 31.8 million perturbed cells^19,20,41–45^. Input features to ExPert were the 3,096 genes that best separated perturbations within each dataset (**Figure S1F, Supplementary note 2**; **Methods**). Of 19,191 perturbations measured in at least one screen, we retained the 8,964 present in at least 60% of contexts (**Figure 1A**, right), so that every perturbation the model learns has been observed in the majority of contexts. The retained perturbations span transcription factors, signaling components, chromatin regulators, metabolic enzymes, and protein-complex subunits (**Table S1C**). The 19.5 million cells carrying these perturbations formed the training set (**Methods**), comprising 61,445 perturbation–context pairs.

**Figure 1.**
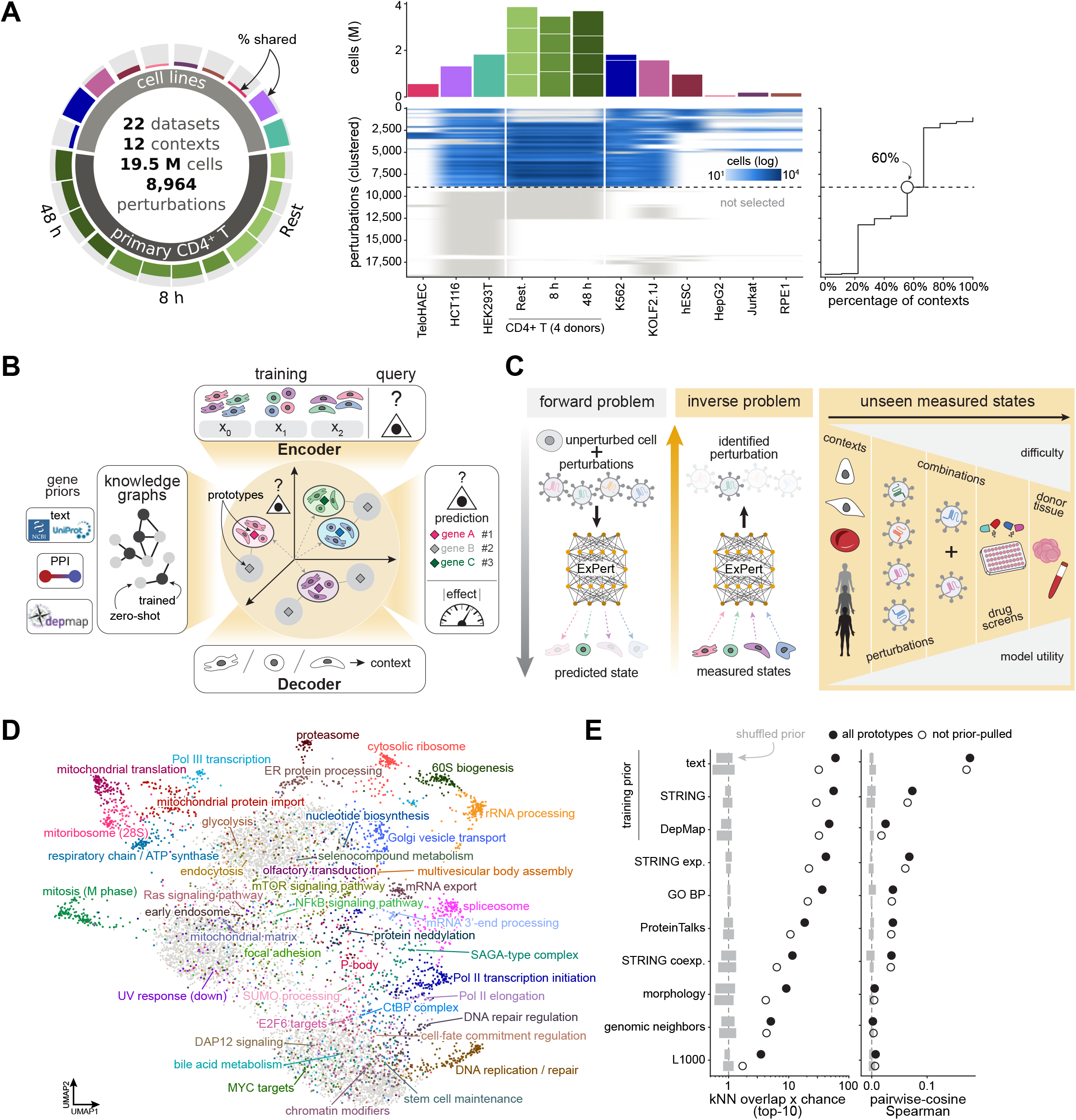
ExPert learns cross-context perturbation prototypes for inverse inference. (A) Training corpus. Left, corpus summary (22 datasets, 12 contexts, 19.5 million cells, 8,964 perturbations). Outer ring, datasets colored by context, with the filled radial extent of each segment giving the percentage of its perturbations shared with other datasets; inner ring, primary CD4⁺ T cells (resting, 8 h and 48 h stimulation) and cell lines. Middle, perturbations (rows; n = 19,191 with ≥10 cells in at least one context, clustered) by context (columns); color, cells per perturbation and context (log scale); gray, measured but not selected for training. Dashed line separates the 8,964 perturbations selected for training. Top, cells per context (millions); CD4⁺ T cell columns are stacked by donor (n = 4). Right, number of perturbations (rows as in the matrix) present in at least the indicated percentage of contexts; open circle, selection threshold (≥60% of contexts; **Methods**). (B) Model overview. Top, cells from multiple training contexts (x_0_, x_1_, x_2_), and query cells at inference (triangle) are embedded by a shared encoder into a common latent space in which each perturbation is represented by a prototype (diamonds). Three terms position the prototypes: proximity to the cells carrying that perturbation (Proxy Anchor loss, with each cell weighted by its measured knockdown effect; center); for perturbations with little training support, pull toward the mean of their neighbors’ prototypes in gene-prior graphs built from gene text descriptions, STRING^51,52^ protein-protein interactions (PPI)^108^, and DepMap^46,47^ co-dependency (left); and reconstruction of the input through a context-conditioned decoder (bottom). Perturbations outside the training set are placed at the mean of their trained prior neighbors (gray diamonds) and serve as negatives during training. At inference (right), each query cell is encoded into the latent space and scored by cosine similarity against all 19,895 prototypes in a single pass, giving a ranked list of perturbations; a separate head predicts the per-cell perturbation strength (|effect|). (C) Forward and inverse problems. Forward models predict the expression state of an unperturbed cell given a perturbation; ExPert addresses the inverse problem, identifying the perturbation that produced a measured state. The same trained model is applied without retraining to five classes of unseen measured states, ordered by increasing difficulty of generalization from the training data and increasing model utility: held-out contexts (cell lines, stimulation conditions, or donors), unseen perturbations, perturbation combinations, drug screens, and donor tissue (Figures 2–6). (D) Uniform manifold approximation and projection (UMAP; cosine distance) of the 8,964 learned prototypes of the final model (one training run on all 22 datasets; also used in (E)). Prototypes are colored by Leiden cluster (79 clusters) where the cluster is enriched for a specific annotation term (44 clusters; hypergeometric test against GO^109^, KEGG^110^, Reactome^111^, CORUM^112^ and MSigDB^113^ Hallmark gene sets; Benjamini–Hochberg *q* < 0.05; term covering ≥20% of the cluster’s annotated members); labels give the top term. Gray, 5,370 prototypes in clusters without a specific enrichment. (E) Agreement between prototype geometry and external gene priors. Left, overlap between each prototype’s 10 nearest neighbors in the latent space and in the prior, relative to chance (log scale). Right, Spearman correlation (ρ) between pairwise prototype cosine similarity and pairwise prior similarity, over 300,000 random pairs of prototypes covered by the prior. Filled circles, all prototypes; open circles, prototypes not pulled toward priors during training; gray boxes, label-shuffled prior (mean ± 2 SD over three shuffles). Bracket, priors used during training. GO BP, Gene Ontology biological process; STRING exp. and STRING coexp., STRING experimental and co-expression channels; morphology, JUMP^114^ Cell Painting CRISPR profiles; genomic neighbors, chromosomal position; L1000^40^, LINCS L1000 consensus signatures. See also **Figures S1, S10** and **S11** and **Tables S1–S3**.

Perturbations were unevenly represented, ranging from tens of cell–dataset observations to more than ten thousand (**Figure S1A**). Common essential genes (DepMap co-essentiality^46,47^) accounted for 47% of the 1,004 least-represented perturbations, compared with 6% of the remaining 7,960 (**Figure S1B**). For these, sparsity likely reflected perturbation-induced lethality rather than a technical problem with the screen. Rather than removing these perturbations, we treated low representation as a signal and used external knowledge to better position them and recover their biological context (**Methods**).

### ExPert learns one prototype per perturbation

We developed ExPert to learn a single representation, per perturbation, that holds across contexts (**Figure 1B,C**). We define that representation as the perturbation’s “prototype”: one direction in a latent space shared by every context, toward which all cells carrying that perturbation are drawn. Four features of the data make this goal challenging. First, baseline gene expression state of each cell type and the technical signature of each screen dominate the transcriptome signal, such that cells group largely by dataset rather than by perturbation. Second, a perturbation’s effect is only partly shared across contexts, since the same knockdown can act differently in distinct cell types (e.g., a T cell versus a hepatocyte) (**Figure S1C**). Third, responses to a single-gene perturbation are weak relative to a cell’s global expression profile (**Figure S1D**). Fourth, the effect size of a perturbation also varies from cell-to-cell, so cells cannot be treated equally in training (**Figure S1E**).

We reasoned that a model constrained to a single prototype per perturbation would be forced to find the common component across contexts. Relaxing that constraint in favor of a conventional classifier cost held-out accuracy under an otherwise identical model (**Table S2A**; **Table S3A,D**). We assigned each of the 8,964 perturbations in the training data a prototype in a 2,048-dimensional latent space (**Figure 1D**), with dimension and encoder depth set by a parameter sweep of both (**Table S2B**).

ExPert compresses each cell’s transcriptomic profile into a vector in that latent space (encoding) and reconstructs the observed gene expression counts from this vector (decoding). Throughout this work, ablations are measured by held-out top-10 recall (top-10 hereafter), always on contexts unseen during training. We centered each cell on the control cells of its own dataset before encoding, removing the baseline expression of its cell type and screen^19,48^. An uncentered input cost fit (-28%) and held-out transfer (-57%), even with an invariance term to compensate (**Table S2D**). Z-scoring or projecting the dataset-specific offset out of the latent space were each worse than centering (**Table S2D**). We conditioned the decoder on a context embedding to absorb the remaining technical variation, rather than removing context adversarially. Adversarial removal cost up to 7.3% of held-out top-10 and fit its own training cells worse by up to 31.7%, consistent with capacity diverted into invariance rather than a failure to converge (**Table S2C**).

To draw each cell toward the prototype of its perturbation and away from all others, we deployed a Proxy Anchor objective^49^. We chose this approach over a softmax head because softmax stopped learning once the correct prototype won by a clear margin, whereas Proxy Anchor kept pushing the other prototypes away (**Figure S10**). A reconstruction term kept the latent space tethered to the observed gene expression counts the model had to decode and remained the largest term in the loss even at half weight; cutting its share of the gradient made the model overfit, gaining fit (34%) and losing transfer (-40%) (**Table S2A**). We weighted cells by knockdown effect size measured with Mixscale⁴⁰ (weighted between 0.1-1; **Methods**) rather than discarding cells with weak effects. Without the weighting, the model gained fit (20%) and lost transfer (-12%), suggesting that it was overfitting to low-effect-size cells when they were not downweighted (**Tables S3B, S2B**). We also predicted per-cell effect size as an auxiliary output, without the perturbation label and differential-expression step that Mixscale requires. The prediction recovered the measured ground-truth value with an average correlation of 0.74 on essential screens (**Figure S11A**), so every query cell received an estimate of perturbation strength alongside its ranking.

So far each prototype was positioned only by the cells carrying its perturbation, which could leave perturbations with few or weakly perturbed cells poorly placed. We therefore drew on external knowledge of gene function as priors: language-model embeddings of gene annotation (e.g. GenePT^50^), STRING protein-protein interactions^51,52^ and DepMap co-essentiality^46,47^; how these were built and selected is described with the zero-shot results (**Methods**). We supplied them only as “neighborhoods”, that is, the set of genes most similar to each perturbation in the prior, rather than as coordinates for where its prototype should sit.

For each gene perturbation present in training, we took its 25 nearest neighbors in each prior and pulled the prototype toward the mean of those neighbors’ prototypes weighted by similarity. We scored each perturbation by the number of training cells it contributed multiplied by the number of datasets those cells came from, and swept how far down that ranking this pull should reach. Held-out top-10 hits rose steeply over the first few percent of that ranking, flattened between 6 and 11%, and fell away once better-supported perturbations began to be moved, meaning the priors helped where training support was low and hurt where it was not. We therefore applied the pull to the 1,004 least-supported perturbations (11.2% of 8,964; **Figure S2A**). How the pull was parameterized made little difference: a weight decaying with evidence, a gate keyed on prior agreement, three evidence statistics, and five pull strengths all fell within run-to-run variation (**Table S2D**). Only extreme pull weights reversed the gain (**Table S2D**). Pulling every trained prototype toward its neighbors raised top-10 hits on held-out queries for the K562 cell type by 2.2% (0.1889 versus 0.1849, *p* = 0.042), and restricting that pull to the least-supported prototypes, as ExPert does, raised them by up to 3.1% (0.1907, *p* = 0.0034; **Figure S2B**). The gain appeared only when the pull was applied during training rather than to a finished model (**Figure S2C-E**; **Supplementary note 1**).

During training, the priors improved top-10 performance only when the ground-truth data set each prototype’s position and the prior adjusted it, not when the prior set the position and the data adjusted it afterward. Both alternatives of the latter cost accuracy: fixing prototypes to the prior embeddings and learning a mapping onto them fell 4.8% below a conventional classifier head, which the prototype constraint outperformed (**Tables S2A, S3C**), and initializing the prototype table from the priors cost 7.0% against random initialization (**Table S3B**).

Since the priors helped only in specific cases, we asked how the learned prototype structure relates to the structure the priors encode. Pairwise similarity in the strongest prior tracked similarity among learned prototypes at only Spearman ρ = 0.17, yet the two agreed on nearest neighbors 32 times more often than chance (**Figure 1E**). The priors therefore carried local neighbor identity but not global geometry. The priors could not leak into the rest of the model: they carried no gradient and moved only the 1,004 least-supported prototypes, leaving the other 7,960 positioned by expression alone (**Methods**). Because that local agreement is what the neighborhood pull already relies on, the same graph can place a perturbation the model never saw, provided the prior covers it: we set its prototype to the mean of its nearest trained neighbors and call these filled prototypes, or fills (**Figure 1B**; gray diamonds). Filling added 10,931 perturbations to the 8,964 learned during training, without retraining; 9,874 of the fills were measured in at least one screen and are used for evaluation below (**Methods**).

The result is a single table of 19,895 prototypes, one per gene perturbation, in a 2,048 dimensional latent space shared by every context. For inferring perturbation from cell state, ExPert requires only this table and the encoder. A query cell is centered on its own controls, encoded to a latent vector, and scored against all prototypes in one pass by cosine similarity (**Methods**), without access to the training cells and without retraining. The model, the prototype table, and the code are available as a Python package (expert, PyPI) and at https://github.com/GoyalLab/ExPert. Model hyperparameters are documented in **Table S4**.

### ExPert recovers perturbations in held-out cellular contexts at scale

We first asked whether ExPert recovers single-gene perturbations from cells it had not seen, both in contexts (cell line, stimulation condition, or donor) it was trained on, and in contexts it was not. We benchmarked different methods on the largest subset of the corpus that every method could complete within a realistic compute budget, 1,157 perturbations and 2,741,678 cells, and evaluated across twelve “folds” (**Methods**). Each “fold” is one train/held-out split of the same corpus, withholding entire datasets or contexts rather than a random sample of cells to avoid leakage. Held-out cells were scored either in distribution (ID), from a dataset seen in training, or out of distribution (OOD), where the whole context had been withheld. We report macro-F1, mean reciprocal rank (MRR), and top-k (5, 10, 100) recall per dataset and fold, scoring pseudobulks of held-out cells unless stated otherwise (**Figure 2**).

**Figure 2.**
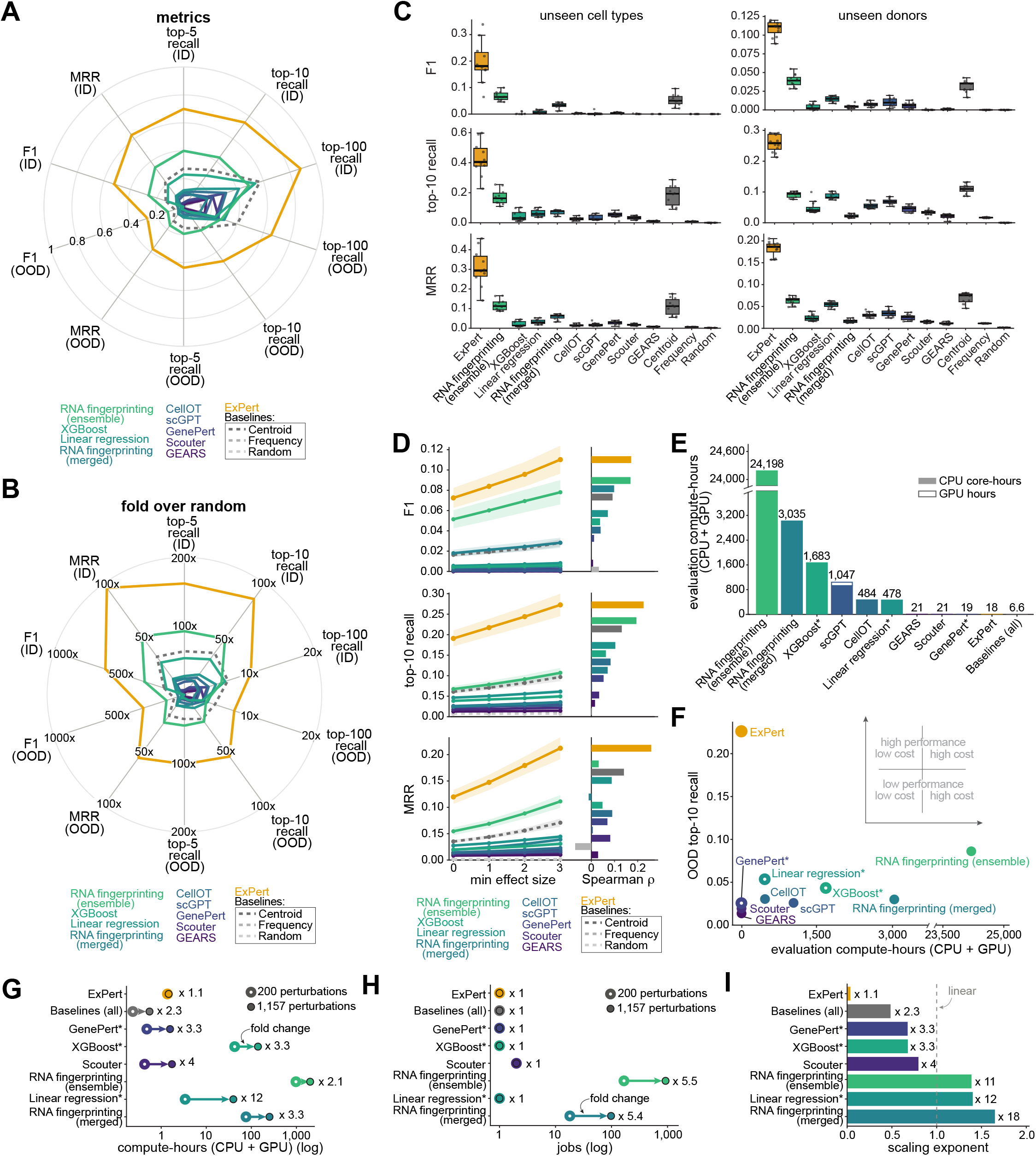
ExPert identifies perturbations in held-out cells and contexts accurately and at low computational cost. (A) Retrieval performance of ExPert, nine comparison methods, and three baselines (Centroid, Frequency, and Random), in-distribution (ID; upper half) and out-of-distribution (OOD; lower half). Spokes show F1, mean reciprocal rank (MRR), and top-5, top-10, and top-100 recall over 1,157 candidate perturbations. Scores are computed on pseudobulks (cell scores averaged per perturbation and dataset), with each perturbation weighted equally. Values are means over n = 230 (ID) and n = 34 (OOD) dataset by fold combinations from 12 folds. Forward-prediction methods were inverted by ranking candidate perturbations by the similarity of their predicted response to the query (**Methods**). (B) The data in (A) expressed as fold change over the expectation under random ranking of 1,157 candidates (e.g., k/1,157 for top-k recall). Each spoke has its own scale. (C) OOD performance on pseudobulks for each held-out dataset by fold combination, split by held-out context: cell line or cell type (left; n = 10) and donor (right; n = 12). Rows show F1, top-10 recall, and MRR. Boxes show the median and interquartile range (IQR); whiskers extend to 1.5x IQR. (D) Left, OOD single-cell performance when only query cells at or above a minimum knockdown effect size are scored, on the same n = 30 dataset by fold combinations at each threshold. Data are represented as mean ± SEM. Right, median Spearman correlation (ρ) between each cell’s knockdown effect size and its score for the metric in that row (top-1 hit, top-10 hit, or reciprocal rank), across n = 34 OOD dataset by fold combinations. (E) Total evaluation compute for scoring 1,157 candidate perturbations, summed over 12 folds, in CPU core-hours (filled) and GPU-hours (open); y-axis broken. (F) OOD top-10 recall on single cells (weighted by knockdown effect size; mean over n = 34 dataset by fold combinations) against total evaluation compute (CPU core-hours plus GPU-hours, summed over 12 folds; x-axis broken). Inset, guide to reading the plot. (G and H) Evaluation cost per fold when the candidate set grows from 200 to 1,157 perturbations: compute-hours (G) and number of SLURM jobs (H), each the mean over 12 folds, on log scales. Arrows run from 200 (open circles) to 1,157 (filled circles) perturbations; numbers give the fold change (xN). (I) Scaling exponent b of the combined evaluation cost (compute-hours by jobs per fold), assuming that cost is proportional to (number of perturbations)^b^ and estimated from the two measurements as b = log(fold change)/log(1,157/200). The dotted line marks linear scaling (b = 1); numbers give the fold change in combined cost (xN). In (E)–(I), asterisks mark methods whose training and evaluation run as a single job, so their cost includes training; in (F), these methods are also drawn as open circles. Baselines (all) is the cost of all baselines, which are computed together in one job. See also **Figures S3** and **S4**.

Scoring existing methods on this task required two accommodations. Of the methods we compared, only RNA fingerprinting^38^ is designed explicitly for the inverse direction, mapping query cells onto a reference of measured perturbation responses. The rest predict forward, so we adapted each method to run in the inverse direction, predicting the state of every candidate perturbation (forward problem) and ranking candidates by similarity to the query (**Methods**). Most are also designed to be fit to a single screen (RNA fingerprinting^38^, CellOT^32^, GEARS^31^, Scouter^35^ and GenePert^53^). For all methods, we trained one model for each dataset-fold combination, and scored test cells without providing cell origin information to the method (**Methods**). Models that can train across datasets (ExPert, XGBoost^54^, logistic regression, scGPT^34^ and the pooled centroid) were fit once per fold on all datasets and scored in a single pass over the shared class space (**Methods**). We also included three baselines that bound what is achievable without a model: Random, which ranks candidates uniformly; Frequency, which ranks perturbations by how often each occurs in the training data; and Centroid, which matches the query to each perturbation’s mean training data response (**Methods**).

ExPert ranked first on every ID and OOD metric (**Figure 2A**). In distribution (ID), several methods fit the training screens well: the RNA fingerprinting (ensemble), the nearest-centroid baseline, and XGBoost reached top-10 recall of up to 0.44 on held-out cells of seen datasets, compared to ExPert’s 0.75 (**Figure 2A**). None of the methods retained their performance once a context was withheld. OOD, ExPert reached an F1 of 0.27, MRR of 0.38, and top-10 recall of 0.48, against 0.12, 0.17 and 0.21 for RNA fingerprinting (ensemble), with every other method at or below the nearest-centroid baseline (0.07, 0.13 and 0.19). Measured against random ranking of the 1,157 candidates, ExPert’s OOD performance was 56-fold above chance for top-10 recall, 57-fold for MRR, and 315-fold for F1, more than twice the enrichment of the next-best method (RNA fingerprinting ensemble 24-, 25-, and 138-fold) (**Figure 2B**).

The twelve folds tested generalization along three axes: a new stimulation condition within an otherwise seen context (same donor, different timepoint), a new cell line or cell type, and a new donor. Held-out stimulation conditions were the easiest case for all methods and preserved the same ordering (ExPert top-10 recall 0.75, MRR 0.63, F1 0.50; RNA fingerprinting (ensemble) 0.36, 0.31, 0.24; **Figure S3A**). Generalization to unseen cell lines and cell types was more difficult, but ExPert maintained a wide margin on evaluation metrics: top-10 recall 0.43, MRR 0.31 and F1 0.20, ∼2.3-fold higher than the best other method on recall (nearest-centroid baseline, 0.18) and roughly 3-fold RNA fingerprinting (ensemble) on F1 (0.07), whose recall (0.17) did not exceed the baseline (**Figure 2C**, left). Held-out donors were the hardest split for every method, ExPert included, consistent with the donor-specific responses reported in the original study^20^. However, ExPert’s margin persisted: top-10 recall 0.26 against 0.11 for the next-best method, MRR 0.18 against 0.07, and F1 0.11 against 0.04 (**Figure 2C**, right). Across folds, ExPert’s median increase in top-10 recall over the next-best method was 2.14 times ID and 2.38 times OOD (**Figure S3B**). Masking the CRISPRi target gene in each query cell lowered recovery for all methods, including ExPert (6–11%), but ExPert still maintained a 2.1-fold lead over the next-best method (**Figure S3C**). The learned prototype space therefore transferred along every axis of the corpus.

ExPert’s margin over other methods held at the single-cell level, though absolute recovery fell for every method: ExPert’s median OOD top-10 recall on unseen cell lines fell from 0.43 on pseudobulks to 0.22, and RNA fingerprinting (ensemble) from 0.16 to 0.07 (**Figure S3D**). Further analysis revealed that the classification failures concentrated in weakly perturbed cells. Restricting the OOD evaluation to cells above increasing thresholds of measured knockdown efficiency raised ExPert’s top-10 recall from 0.19 to 0.27, MRR from 0.12 to 0.21, and F1 from 0.07 to 0.11, roughly twice the improvement over any other method (**Figure 2D**, left). Within cells, knockdown effect size predicted retrieval for every learned method far above a shuffled null, and translated into the largest absolute gain for ExPert: its top-10 hit rate rose from 6% of cells in the weakest knockdown quartile to 30% in the strongest, compared with 2% to 12% for RNA fingerprinting (ensemble) (**Figure 2D**, right). A random subsample of the same sample size at each threshold showed no such rise (**Figure S3E**), so the effect can be attributed to perturbation strength and not to sample size. RNA fingerprinting left many cells unassigned (20% ID, 54% OOD), but ExPert’s lead held when both methods were restricted to the cells it assigned, and at every confidence threshold (**Figures S4A–S4C**). ExPert predicts perturbation strength for every query cell, including cells from unseen contexts (**Figure 1B, Figure S11**), so each call carries its own estimate of confidence.

We next evaluated compute cost. ExPert scored all 12 folds in 18 CPU core-hours, less than every other method we tested apart from the untrained baselines (**Figure 2E**). The RNA fingerprinting ensemble reached its OOD accuracy by fitting a separate model for every dataset by fold and scoring every query cell against all of them, which took 24,198 CPU-hours (**Figure 2E**). Merging the RNA fingerprinting models into one reduced this to 3,035 hours but removed most of the accuracy (**Figure 2F**). Every other method was either computationally cheap and no better than baseline, or expensive and still below 0.05 top-10 recall OOD (**Figure 2F**). ExPert was the computationally cheapest and most accurate, and those closest in accuracy required at least two orders of magnitude more compute cost (**Figure 2F**).

The compute gap widened as the number of candidate perturbations grew. Increasing the candidate set 5.8-fold, from 200 to 1,157 perturbations, made each job more expensive for every method (**Figure 2G**). For the RNA fingerprinting approaches, it also multiplied the number of jobs about 5-fold, because each perturbation needed its own model (**Figure 2H**). Computational cost rose 11-fold for the RNA fingerprinting ensemble, 12-fold for logistic regression and 18-fold for the merged RNA fingerprinting, with peak memory climbing to 85–100 GB per job (**Figure 2I**). In contrast, ExPert’s cost rose 1.1-fold and its peak memory stayed at about 21 GB (20.3 GB for 200 and 20.6 GB for 1,157 candidates), because it scores every perturbation prototype in one pass regardless of the number of perturbations. We attempted the same evaluation against the full set of 8,964 perturbations and found the per-dataset ensembles computationally infeasible; ExPert scored the complete set in about 11 minutes per fold (1.5 CPU core-hours; 18 core-hours across all 12 folds).

### ExPert zero-shots perturbations absent from training

We placed a perturbation prototype absent from training at the mean of its nearest trained neighbors in the prior (**Figure 3A**). We reasoned that if the priors carry information about response, that placement should land near where the gene’s own prototype would have been; if they do not, it is arbitrary, and retrieval would be no better than random ranking. This is zero-shot identification in the sense used for image classes described only by attributes^55,56^, with gene priors taking their role.

**Figure 3.**
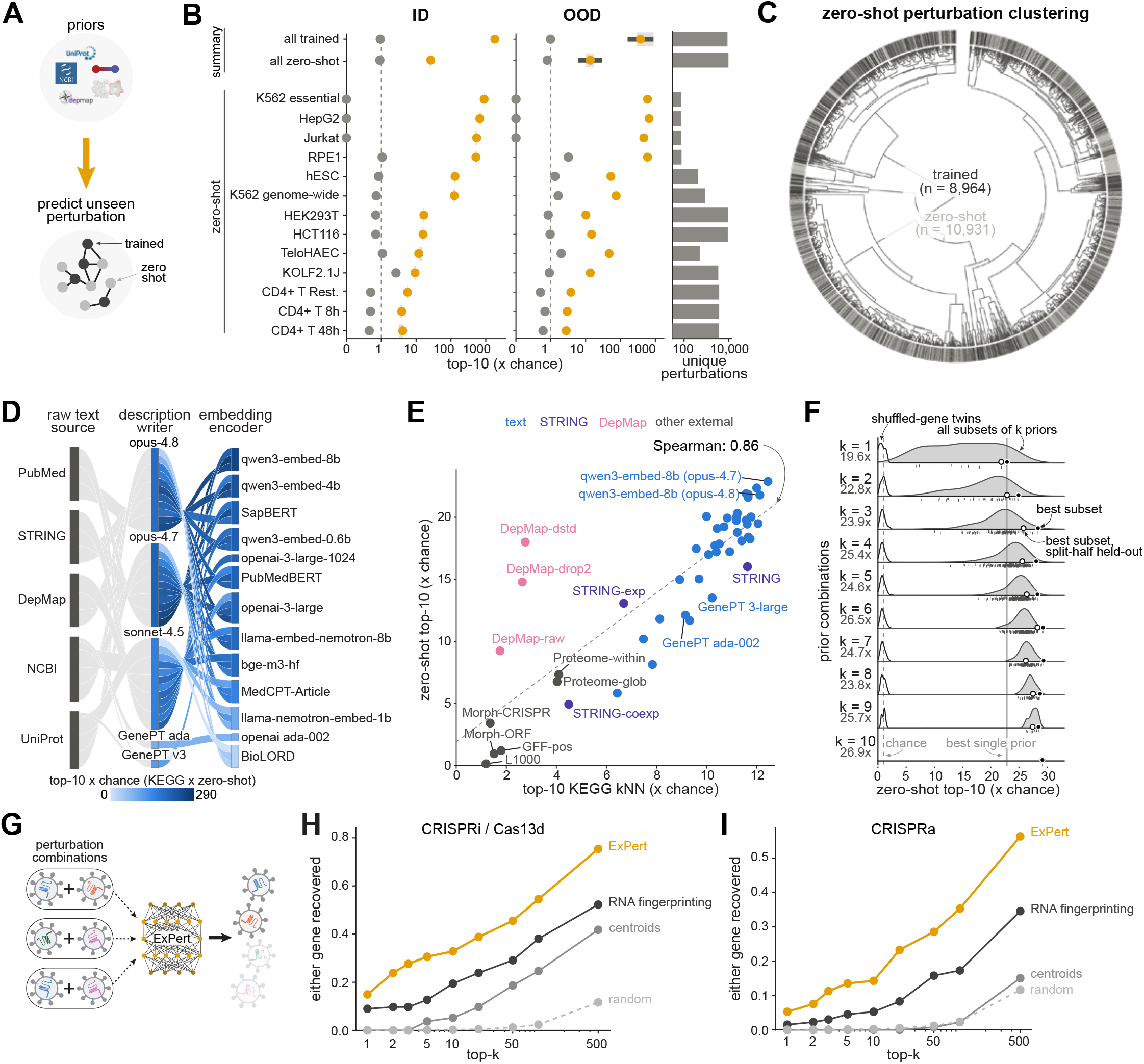
Prior gene knowledge extends ExPert to perturbations absent from its training screens. (A) Schematic. Prior gene knowledge (text descriptions, protein interactions, genetic dependencies; top) places a prototype for every gene in the model’s vocabulary (bottom): trained perturbations (dark gray) are learned from screens; unseen (zero-shot) perturbations (light gray) are positioned from their prior neighbors. (B) Zero-shot retrieval. Each query is the pseudobulk of up to 25 cells of one perturbation in one screen, ranked against all 19,895 prototypes. Score, top-10 recall weighted by the within-screen percentile of knockdown effect size, expressed as fold over chance (chance = 10/19,895). Left, in-distribution (ID): screens from contexts seen in training. Middle, out-of-distribution (OOD): screens from the context held out by that model; each context is held out by one model, so OOD screen rows show a single value. Summary rows pool all screens: all trained, held-out cells of the 8,964 trained perturbations, scored against the same 19,895 candidates; all zero-shot, the 9,874 of 10,931 zero-shot perturbations measured in at least one screen. Remaining rows, zero-shot on individual screens, ordered by ID median, with the CD4^+^ T cell conditions in time order. Orange dot, median over models; light gray box, interquartile range; dark gray bar, ±1 SD of log-transformed values. Gray dot (random), zero-shot prototypes reassigned to random genes (mean of 20 shuffles per model). Dashed line, chance. Right, unique perturbations scored per row. n = 12 models (one training run each), each trained with one cell line, donor, or condition held out. (C) Zero-shot prototypes fall throughout the trained prototype space. Circular dendrogram (Ward linkage, cosine) of the 8,964 trained prototypes of the production model; each of the 10,931 zero-shot prototypes is placed beside its most similar trained prototype. Outer ring, trained (dark gray) and zero-shot (light gray) leaves. (D) Generation of gene text embeddings. Source text (left) is summarized by a description-writing language model (middle) and encoded by an embedding model (right); each ribbon is one writer-encoder combination (n = 36), including the published GenePT embeddings^50^. Color, zero-shot top-10 recall (fold over chance) multiplied by KEGG^57^ neighbor lift. (E) Prior quality predicts zero-shot retrieval. Zero-shot top-10 recall (effect-size-weighted, fold over chance) when a single prior positions the zero-shot prototypes, against a training-free measure of the prior’s biological content: lift over chance in KEGG^57^ pathway co-membership among each gene’s ten nearest neighbors, computed on a size-matched set of 2,000 genes (as in (D)). All priors are scored with one model trained without prior information (K562 held out). Points, individual priors (n = 48): text (n = 36, **Methods**), STRING^52^ (n = 3), DepMap^46,47^ (n = 3), and other external sources (n = 6, **Methods**). Dashed line, least-squares fit. Spearman ρ = 0.86 (all priors), 0.82 (text), 0.64 (non-text). (F) Combining priors. Zero-shot top-10 recall (as in (E)) for every subset of the ten best single priors, by subset size k (number of subsets = 10 choose k). Gray, subsets; white, the same subsets with gene labels shuffled. Filled circle, best subset; open circle, best subset selected on one half of the perturbations and scored on the other half. Values below each k, held-out best subset. Vertical lines, best single prior (dashed) and chance (dotted). (G) Schematic of combinatorial perturbation recovery. Pseudobulk queries from double perturbations (left) are ranked by ExPert (middle) against the single perturbations in its trained vocabulary (right). (H) Recovery of combinatorial loss-of-function perturbations by the production model. Fraction of double perturbations for which either targeted gene ranks within the top-k of the 8,347 candidate perturbations scorable by all methods. Orange, ExPert; dark gray, RNA fingerprinting^38^ custom-trained on genome-wide K562 Perturb-seq^19^; mid gray, nearest centroid of each perturbation’s training cells; light gray dashed, random ranking. *n* = 134 double perturbations from a CRISPRi screen^60^ (*n* = 10) and a Cas13d screen (CaRPool-seq^61^; *n* = 124). (I) Recovery of combinatorial gain-of-function perturbations, as in (H). *n* = 133 double perturbations from two CRISPRa screens (Norman et al.^62^, *n* = 77; ProPer-seq^63^, *n* = 56). None of the double-perturbation screens in (H) and (I) were used in training. Statistics: (E) Spearman correlation, two-sided (all priors, *p* < 0.0001; text, *p* < 0.0001; non-text, *p* = 0.026). See also **Figures S2, S5**, and **S6**, **Tables S6** and **S7**, and **Methods**.

ExPert placed each perturbation absent from training by the same pull toward prior neighbors used to augment weakly supported prototypes during training (**Figure 1B**). Genes not measured in any screen carried no evidence, so the rotation ran to completion and their prototypes took the mean of their neighbors. Those neighbors came only from the remaining 7,960 best-supported trained perturbations (cross-fitted; **Methods**). These filled perturbations were excluded from training by the context-coverage filter, but 9,874 of them were measured in at least one screen, so ground-truth labels existed for them (**Figure 1C**; **Methods**). We asked whether ExPert ranked the correct gene in the top-10, against the full 19,895-gene vocabulary (chance 10/19,895), with a null in which each fill was assigned to a shuffled gene. Perturbations absent from training were recovered at 26.2-fold over chance overall, with the shuffled null at chance (**Figure 3B**). The gain depended on the screen the query came from: 4–17-fold in the primary T cell, iPSC, and genome-wide HEK293T and HCT116 screens, where most such genes have weak effects, and 520–900-fold in the essential-gene panels of HepG2, RPE1, Jurkat, and K562, whose perturbations are strong (**Figure 3B**). Top-10 recovery held when the query’s cell line, donor, or condition had also been withheld from training, at 13.5-fold overall (**Figure 3B**, OOD).

The 10,931 prior-defined prototypes were not confined to a separate region of the space. Each sat at the similarity-weighted average of its 25 nearest neighbors’ prototypes, and together they spread across almost all clusters of the trained-prototype tree (**Figure 3C**). Although overall similar, missing prototypes lay closer to the mean prototype direction than trained prototypes did (average angle 85.8° vs 89°), which bounds how finely two genes absent from training can be discriminated (**Figure S6A**). Concentrating weight on the nearest neighbors mattered more than how many neighbors were used, raising zero-shot recovery 9% ID and 16% OOD at essentially no cost to trained perturbations (**Figures S6B and S6C**; **Supplementary note 3**). Fills had to be present during training as negatives; adding them only afterwards displaced trained perturbations from the top-10 (**Figure S6E**).

Zero-shot placement depends entirely on the priors, so we asked which sources carried the information and how the text priors should be built. We varied how the text embeddings were produced, testing five raw text sources, three description generators, and thirteen embedding encoders, alongside the published GenePT^50^ embeddings (**Figure 3D**; **Methods**). To measure each prior’s biological content without reference to ExPert, we asked how often a gene’s ten nearest neighbors in that prior shared a KEGG^57^ pathway with it, relative to random gene pairs. That enrichment predicted zero-shot recovery across all 48 priors tested, including STRING^51,52^, DepMap^46,47^, and morphology (Spearman ρ = 0.86; **Figure 3E**). Combining text sources helped; selecting on one half of the perturbations and scoring on the other, the best single prior recovered perturbations at 21.9-fold over chance and combinations reached 29.2-fold. Three sources reached 25.8-fold, and shuffled controls stayed at chance throughout (**Figure 3F**; **Methods**). The final model fused three independent sources with equal weight: gene descriptions written by Claude Opus 4.8 and encoded with qwen3-embed-8b^58,59^, DepMap, and STRING. Combinations that scored higher almost always contained two versions of the same source (407 of 425; **Table S7**), such as a second text encoder or a second DepMap derivative, which under equal-weight fusion amounted to up-weighting that source. ExPert therefore recovered perturbations it had never been trained on, with the quality of the priors setting performance.

### ExPert recovers combinatorial perturbations without training on them

We questioned whether ExPert, trained only on single perturbations, recognizes combinations of perturbations, using four double-perturbation datasets. On two double-knockdown screens using CRISPRi (Replogle et al.^60^, n = 10) or Cas13d (CaRPool-Seq^61^, n = 124), queried without retraining (**Figure 3G**), we scored whether either constituent ranked in the top-k of the 8,347 candidates common to all methods. ExPert ranked at least one constituent first for 14.9% of queries, 1.7-fold more often than RNA fingerprinting^38^, and within the top-500 for 75%, while centroids and random ranking were near zero at the top ranks (**Figure 3H**).

We then asked whether the prototypes extend to gain of function. CRISPRa, which activates rather than represses its targets, tested a direction of perturbation absent from ExPert’s training. On 142 single CRISPRa perturbations from two screens, ExPert ranked the activated gene in the top-10 of 8,347 candidates 15.5% of the time, against 8.5% for RNA fingerprinting and 0.12% by chance (**Figure S5A,B**). Given that ExPert could read CRISPRa singles, we next predicted double-gene activations. Over two CRISPRa double-perturbation screens (Norman et al.^62^, n = 77; ProPer-seq^63^, n = 56), ExPert ranked at least one constituent first for 5.3% of queries, 3.5-fold more often than RNA fingerprinting, and within the top-500 for 56%, while centroids were indistinguishable from random (**Figure 3I**). Recovery was lower than for knockdown, consistent with activation not being the mirror image of inhibition: raising a gene’s expression can engage different programs than lowering it. There were too few untrained genes (23 total) across these screens to test zero-shot recovery of combinations. We found the same performance ordering for CRISPRi and CRISPRa doubles when requiring both constituent perturbations in the top-k, with every method lower in absolute terms (**Figure S5C-F**).

### ExPert recovers the targets of small molecules without retraining

We asked whether ExPert recognizes perturbations delivered by a different approach. Small molecules act on proteins rather than transcripts, so a drug screen separates the consequence of losing a gene’s encoded protein function from transcriptional repression.If the prototypes capture the consequence of losing a protein’s function, a compound inhibiting that protein should fall near the prototype of the gene encoding it, although no small molecule compounds appear in training. We tested this on two single-cell drug screens: sci-Plex3^21^, covering 188 compounds at four concentrations in three cell lines, and Tahoe-100M^64^, covering 379 small molecules at three doses across 50 cell lines, 49 of which are absent from our corpus. We queried each drug and dose against trained prototypes (**Figure 4A**) without retraining, fine-tuning, or recalibrating the model.

**Figure 4.**
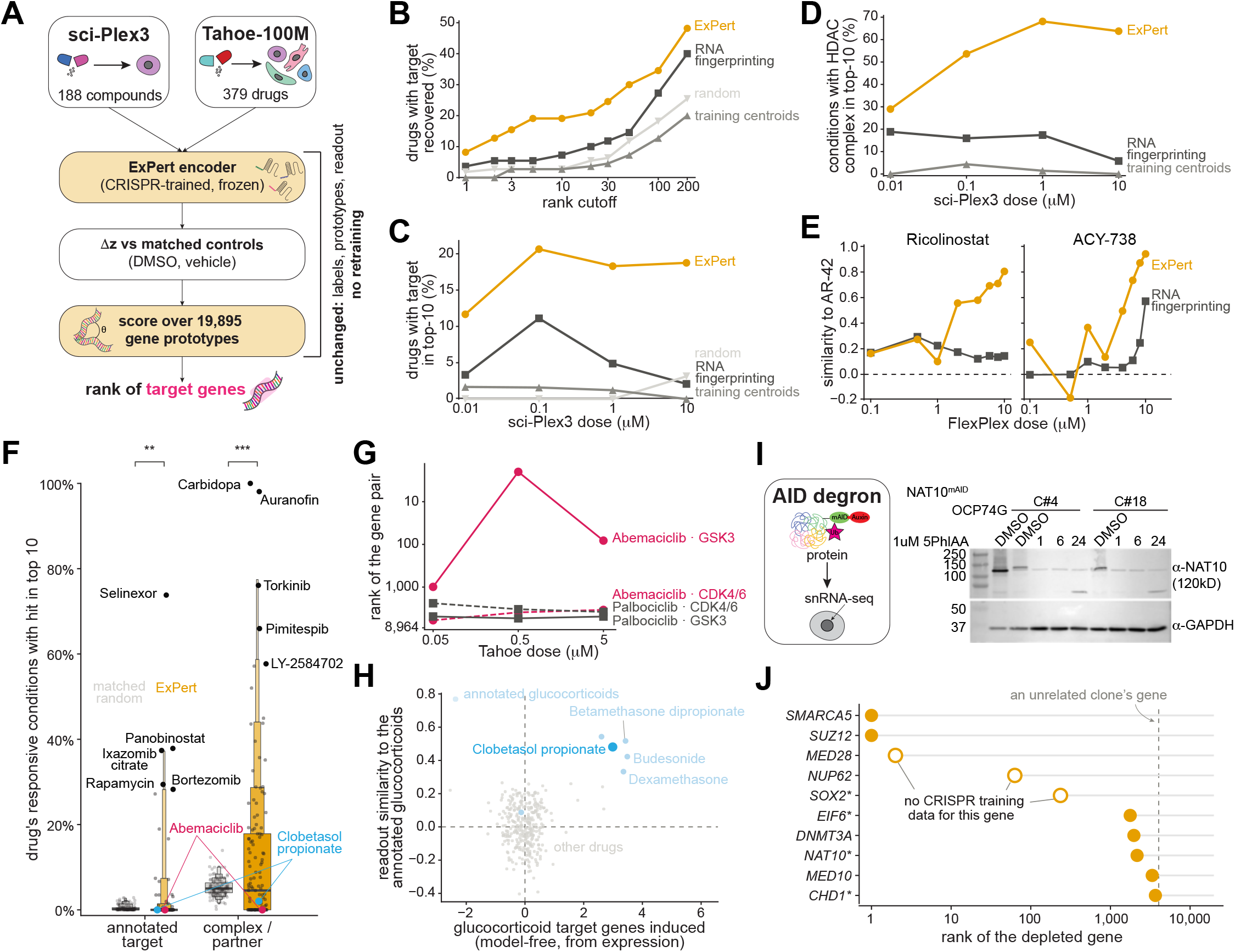
Drug and protein-depletion responses are read out through CRISPR-trained prototypes without retraining. (A) Schematic of the readout. Responses to small molecules (sci-Plex3^21^, Tahoe-100M^64^) are encoded by the frozen, CRISPR-trained ExPert model. Each response is taken as the change in latent state (Δz) relative to matched controls (DMSO or vehicle), scored by cosine against the gene prototypes, and summarised as the rank of the annotated target gene. Labels, prototypes and readouts are unchanged from CRISPR training. (B) sci-Plex3^21^ annotated drug-target recovery as a function of rank cutoff k, over the 110 of 188 compounds that all four methods can score (a compound enters only if it carries an annotated target inside the shared candidate panel and has at least one responsive condition; 179 of 188 carry an annotation). Every method reads the same cells (all vehicle cells and up to 200 cells per cell line by drug by dose, replicate plates pooled), with one input per cell line by drug by dose (**Methods**). A drug counts as recovered at k when any of its responsive conditions places an annotated target at or above k. Orange, ExPert; dark gray, RNA fingerprinting^38^; mid gray, the centroid of each perturbation’s training cells; light gray, a uniform random ranking. All four methods rank the same 3,802 candidates. (C) The same recovery as in (B) at k = 10 resolved by labelled dose, over the drugs responding at that dose (n = 60, 63, 82, and 96 drugs at 0.01, 0.1, 1, and 10 µM; the same drug set for every method at each dose). (D) HDAC inhibitors only, scored against the 48-gene HDAC machinery (SIN3, CoREST, NuRD and NCoR complexes, the HDAC1/2/3 catalytic subunits, and CORUM-core partners of the HDAC family). Plotted is the fraction of conditions placing a machinery gene in the top-10, by dose. Every method is scored on the same conditions: 92 drug by dose pairs (23 HDAC inhibitors by 4 doses), one condition per cell line (n = 69 conditions per dose). (E) Similarity of the HDAC6-selective inhibitors ricolinostat and ACY-738 to the pan-HDAC inhibitor AR-42 across 8 concentrations (0.1–10 µM) in the FlexPlex screen of K562 cells^38^, as the Spearman correlation of their candidate rankings with AR-42’s at the same concentration, one value per compound and concentration. Orange, ExPert v0.1.0; dark gray, RNA fingerprinting. (F) Tahoe-100M^64^ recovery of annotated drug targets and of target complexes within the top-10 of the 8,964 trained prototypes, over the 140 drugs with at least three responsive conditions. For each drug, the fraction of its own responsive conditions placing an annotated target (or a partner of one in a physical complex or tight pathway) in the top-10. Orange, observed; gray, the same statistic for size-and popularity-matched random gene sets drawn for that same drug. Boxes are letter-value summaries of the across-drug distribution; each dot is one drug, with observed values to the right of each pair and matched-random values to the left. Black, the five best-recovered drugs in each group, named with leader lines. Abemaciclib (red) and clobetasol propionate (blue) dots are highlighted. Each drug is compared with the matched-random gene sets drawn for that same drug by a one-sided paired sign-flip permutation test on the mean difference (20,000 permutations); *p < 0.05, **p < 0.01, ***p < 0.001, n.s., not significant. (G) Rank of the annotated CDK4/CDK6 targets (dashed) and of the abemaciclib-specific secondary target GSK3α/GSK3β (solid) for abemaciclib (red) and palbociclib (gray) across 3 concentrations (0.05–5 µM) in Tahoe-100M^64^, each plotted as the mean rank of the gene pair among the 8,964 trained prototypes. (H) Similarity of clobetasol propionate to the glucocorticoids labelled in Tahoe-100M^64^. One dot per drug: y, the mean Pearson correlation of the drug’s ExPert score profile over the 8,964 prototypes with those of dexamethasone, triamcinolone, betamethasone dipropionate and budesonide, over drugs with at least two active doses; x, the induction at 5 µM of eight glucocorticoid target genes (*FKBP5*, *TSC22D3*, *PER1*, *DUSP1*, *KLF9*, *CEBPD*, *DDIT4*, *ERRFI1*) from expression, as the mean z across 379 drugs of the log fold change against same-plate, same-line DMSO. Light blue, drugs annotated as glucocorticoids in Tahoe-100M^64^; blue, clobetasol propionate; gray, other drugs. (I) Left, schematic of the custom auxin-inducible degron (AID) system, in which target proteins are depleted upon auxin exposure. Right, representative immunoblots of degron protein depletion after 1, 6, and 24 h of exposure to the auxin analogue 5-Ph-IAA (1 µM) compared with DMSO, in two NAT10-mAID clones (C#4 and C#18) and the OCP74G line. Upper blot, NAT10; lower blot, GAPDH as the loading control. One experiment; C#4 and C#18 are two independently derived NAT10-mAID clones. (J) An in-house auxin-inducible degron village, in which each clone degrades one tagged protein after 24 h of auxin exposure. Shown are the 10 best of the 18 responding clones (of 25 scored), as the rank of the depleted gene among all 19,895 prototypes, which include zero-shot perturbations never seen in CRISPR training. Open rings, genes with no CRISPR training data; asterisked gene names are ranked in the activation rather than the depletion direction. Dotted line, permutation null, i.e., the median rank that the other clones’ genes reach in the same query (uniform chance is rank 9,948). *SMARCA5* and *SUZ12* rank 1^st^, *MED28* 2^nd^, *NUP62* 63^rd^, and *SOX2* 236^th^; three of these five have no trained prototype. Across all 25 clones, the median rank is 3,492 against a null of 4,784. The improvement over each clone’s own permutation null is tested one-sided by sign-flip permutation (20,000 permutations; *p* = 0.0023) and by a paired t-test on log_10_ rank (p = 0.0076). A single pooled degron experiment with one 10x Multiome library per arm (26,956 DMSO and 31,475 auxin-treated nuclei after quality control), in which the nuclei of each of the 25 clones are the replicate unit. Statistics. In (B) and (C), rank comparisons between methods use a one-sided paired sign-flip permutation test on log₁₀ rank (20,000 permutations) and comparisons of the top-10 hit itself use an exact two-sided McNemar test on the discordant drugs. No error bars are plotted in any panel; (B)–(D) and (F) show proportions over the stated denominators, (E) and (G) show one value per compound and concentration, and (H) one value per drug. Replicates: (B)–(D) are the published sci-Plex3 screen^21^, with conditions defined as cell line by drug by dose; (E) is the published FlexPlex screen^38^, with conditions defined as drug by dose; (F)–(H) are Tahoe-100M56, with conditions defined as cell line by dose. See also **Figures S7–S9**,**Tables S8**, **S9,** and **Methods**.

On sci-Plex3, ExPert ranked a true annotated target (**Methods**) first for 8.2% of 110 compounds that every method could score, within the top-10 for 19.1% and within the top-50 for 30.0%, against 3.6%, 7.3% and 14.5% for RNA fingerprinting^38^ (**Figure 4B**). Training centroids stayed close to the random baseline at every cutoff, and neither named a target first (top-1). Over the top-10, ExPert recovered a target for 15 compounds that RNA fingerprinting did not, and RNA fingerprinting for 2 that ExPert did not (McNemar *p* = 0.0023). ExPert recovered more targets at every dose (**Figure 4C**), and held that recovery as dose increased. Both methods peaked at 0.1 µM, but RNA fingerprinting then declined about fivefold, to 2.1% at 10 µM, while ExPert retained most of its recovery (18.3% at 1 µM and 18.8% at 10 µM, against 20.6% at 0.1 µM; **Figure 4C**).

Drugs commonly act on more than one protein^65^, a known limit on matching them to single genetic perturbations^38^, with HDAC inhibitors presenting a clear case. Compounds sold as isoform-selective engage much of the class I and class II family^66^, and HDAC1 through HDAC3 act within multi-subunit corepressor complexes rather than alone^67^, so losing a scaffold subunit phenocopies losing the targeted enzyme. Across the HDAC-targeting compounds in sci-Plex3, ExPert placed the annotated isoform or a member of its complex, including SIN3, CoREST, NuRD and NCoR subunits, in the top ten for 29.0% of conditions at the lowest dose, rising to 68.1% at 1 µM (63.8% at 10 µM), exceeding all other methods at every dose (**Figure 4D**).

Drug target annotations are not always reliable. Chemoproteomic profiling of 53 HDAC drugs found that several widely used compounds engage targets well beyond their nominal target, HDAC6 inhibitors in particular^68^. We examined this on a FlexPlex screen^38^ of K562 cells containing two nominally HDAC6-selective compounds, ricolinostat and ACY-738, alongside the pan-HDAC inhibitor AR-42. AR-42 provides a reference for broad HDAC engagement, so if ricolinostat and ACY-738 lose selectivity at higher doses, their target rankings should shift toward it. With ExPert, the similarity of ricolinostat and ACY-738 to AR-42 (the Spearman correlation of their candidate rankings) rose with dose overall for both compounds, reaching 0.80 for ricolinostat and 0.94 for ACY-738 at the highest concentration, against 0.14 and 0.57 respectively for RNA fingerprinting on the same cell type (**Figure 4E**). RNA fingerprinting reached a similar conclusion previously, but required a reference assembled for that question: CRISPRi fingerprints for ten HDAC genes, Perturb-seq profiles generated for that study, and 311 drug-dose RNA fingerprints from sci-Plex3^21^. ExPert scored the same queries against its existing prototypes, with no new data and no retraining. Moreover, ExPert performance did not depend on having seen the cell line: with both K562 screens held out, it still recovered twice as many top-10 targets as RNA fingerprinting in sci-Plex3 (14.5% against 7.3%) and matched the full model on HDAC machinery recovery and AR-42 similarity (**Figure S7**).

We extended this to Tahoe-100M, which pairs small molecules with cell lines absent from our corpus, thus changing modality and context at once. We queried each drug and cell line against all of ExPert’s trained prototypes (**Methods**). ExPert enriched annotated targets, members of the target’s complex, and direct interaction partners in the top-10 by 2.4-to 5.6-fold over a matched random ranking (**Figure 4F**). We further probed the queries whose annotated target ExPert did not recover (highlighted in red and blue in **Figure 4F**), and discovered that several were not misses. Abemaciclib is annotated as a CDK4/6 inhibitor, but competitive chemical proteomics found GSK3α and GSK3β among its most potently inhibited off-targets, an activity palbociclib and ribociclib do not share and one that activates β-catenin-dependent WNT signaling^69^. ExPert ranked GSK3 highly for abemaciclib and not for palbociclib (**Figure 4G**). In another case, Tahoe-100M names only a secondary activity: clobetasol propionate is a potent glucocorticoid but is labeled by its CYP3A4/5 inhibition. ExPert placed it with the other glucocorticoids in the screen and ranked canonical glucocorticoid response genes highly (**Figure 4H**), matching the drug rather than its label. Adding filled zero-shot prototypes improved recovery relative to chance on sci-Plex3, from 25.6-to 37.8-fold over random at the top-10 despite ranking 2.2 times as many candidates, and held it on Tahoe-100M (5.0-against 5.6-fold for annotated targets; **Figure S8**). We manually reviewed 22 of the 125 queries scored as misses, selecting known prodrugs, compounds closely resembling another drug class, and compounds that activate rather than inhibit their target, which our ranking did not score. We also included queries whose top-ranked perturbations agreed across active doses more than randomly chosen wells from the same plates (*q* < 0.05). 13 of these 22 queries recovered biology consistent with the compound’s known pharmacology, beyond the annotated activity (**Table S9, Supplementary note 5**). Because the remaining misses were not reviewed, this is a lower bound.

We also asked whether the prototype space encodes the pathway structure connecting drug classes (**Methods**). A graph over the prototypes placed 24 genes between the targets of different drug classes, 12 of them also supported by gene co-essentiality, curated pathway annotation, or literature-derived gene descriptions, though no single prior supported all 12 (**Figure S9A**). The connecting genes included *PDPK1* between the mTOR and PI3K inhibitors, and *RALGDS* between the MEK and FGFR inhibitors (**Figure S9B**, **Supplementary note 6**).

### ExPert recognizes loss of function delivered by protein degradation

CRISPRi suppresses gene transcription while small molecules typically inhibit protein activity without necessarily degrading the protein. Neither directly removes the protein while leaving the corresponding gene’s transcription unaffected. To test whether ExPert reads the loss of a protein’s function independently of how that loss is produced, we generated a degron dataset for this study by pooling clones carrying auxin-inducible degron tags on different proteins, inducing degradation with 5-Ph-IAA, and profiling them by single-nucleus RNA sequencing (snRNA-seq) after 24 hours (**Figure 4I**, left; **Methods**). Western blots confirmed that the tagged protein was depleted within hours of induction and was unchanged in the vehicle (representative example in **Figure 4I**, right). CRISPR screens typically profile cells several days after perturbation^19,70^, so this experiment also tests how early the signature becomes readable. Eighteen of the 25 degron-tagged genes showed a reproducible transcriptional response across clones; the remaining seven were indistinguishable from vehicle-treated controls and were excluded (**Methods**). ExPert placed the correct perturbation in the top 10 for 3 of them, against 0.05% expected by chance (332-fold, *p* = 1.0 × 10⁻⁷). *SMARCA5* and *SUZ12* each ranked first out of 19,895 candidates (**Figure 4J**). *MED28* and *NUP62* were never perturbed in training, yet ExPert ranked them 2^nd^ and 63^rd^, extending its zero-shot capability to a new assay and cellular context (**Figure 4J**). Interestingly, ExPert ranked *SOX2* 236 of 19,895 but placed it on the gain side rather than the loss side which could be a property of the auxin-induced degron system or the documented *SOX2* autoregulation^71,72^ (**Supplemental note 7**).

### ExPert uncovers aberrant mRNA export and splicing in alcohol-associated liver disease

We next applied ExPert to human disease to test whether it could identify regulatory processes not apparent from differential expression alone. ALD is the leading cause of chronic liver disease and its incidence is rising^73^. Abstinence remains the only proven cure, and once liver failure develops, transplantation is the only life-saving intervention^74^. We profiled nuclei from healthy liver (n=5) and from patients with alcohol-associated hepatitis (AH, n=5) or cirrhosis (AC, n=5) by snRNAseq^73^, and scored hepatocyte nuclei against the ExPert prototype table (**Figure 5A**). The highest-scoring perturbations for ALD nuclei included nucleoporins and nuclear transport factors (*NUP153*, *THOC7*, *RANGAP1*), splicing factors (*SRSF6*), and chromatin regulators (*SMARCA4*, *RAD21*, *SMC1A*, *CSNK2B*) (**Figure 5B**). Ranking genes by log2 fold change and ExPert score gave no overlap (Spearman ρ = −0.04 in AC and −0.03 in AH; **Figure 5C**): ExPert’s top calls sat near the origin of both volcano plots while the top differentially expressed genes sat at the extremes (**Figure 5D**). ExPert’s ranking was therefore uncorrelated with differential expression.

**Figure 5.**
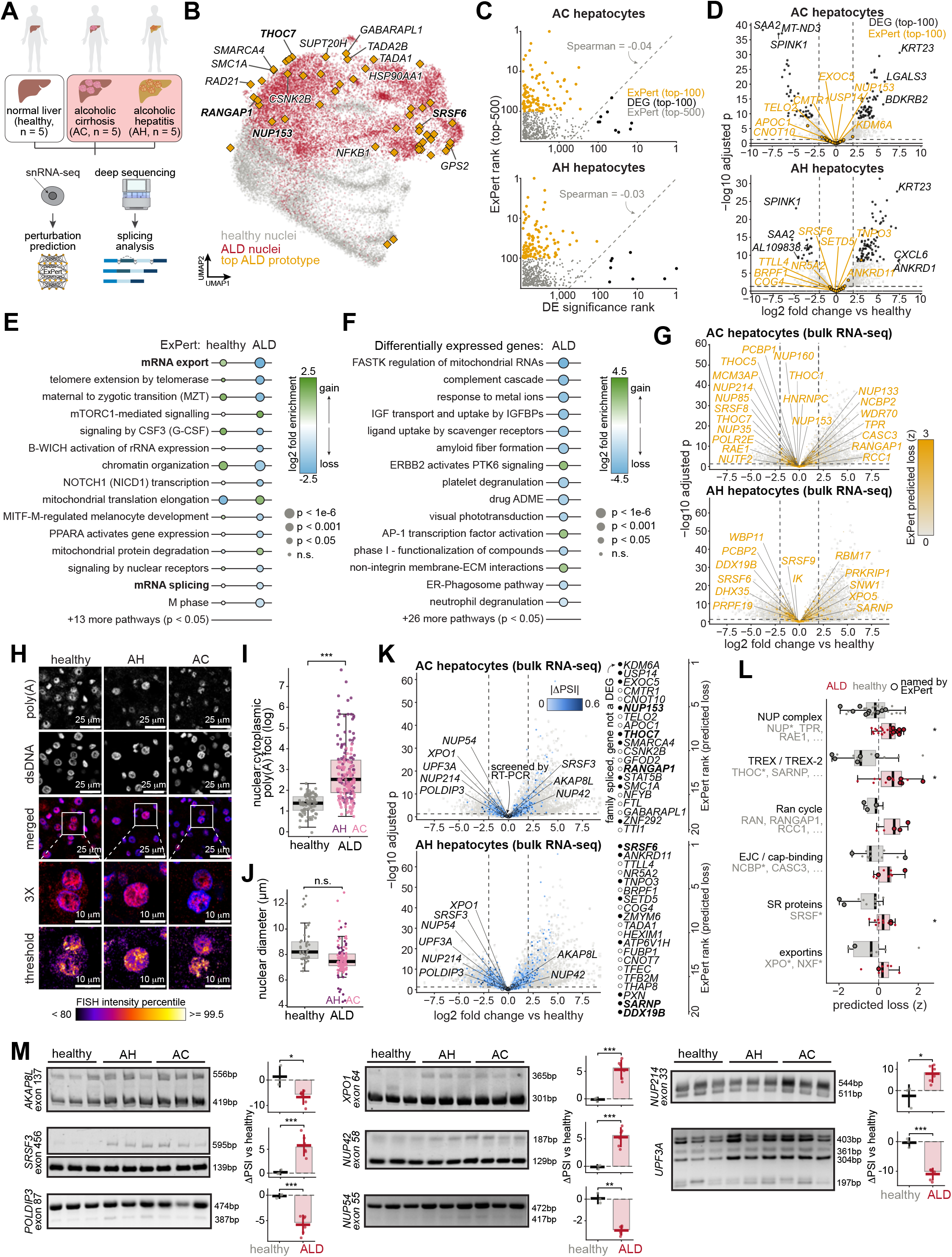
ExPert identifies mRNA export dysregulation in alcohol-associated liver disease. (A) Study design. Liver tissue from 15 donors (healthy, alcohol-associated cirrhosis [AC], and alcohol-associated hepatitis [AH]; 5 donors per condition)^73^, pooled into two libraries per condition, was profiled by single-nucleus RNA sequencing (snRNA-seq) for ExPert inference. The same cohort (healthy, n = 4; AC, n = 6; AH, n = 6 livers) was profiled by bulk RNA-seq for differential expression (DE) and splicing analysis. (B) UMAP of 26,807 nuclei from five liver cell types in the ExPert latent space. Gray, healthy; red, alcohol-associated liver disease (ALD; AC and AH). Orange diamonds, the 25 highest-scoring perturbation prototypes for each of AC and AH (scores averaged over the five cell types). Labeled prototypes belong to mRNA export or splicing (bold), chromatin modification, SUMOylation, or macroautophagy gene sets. (C) ExPert rank of the top-500 predicted perturbations versus DE rank (pyDESeq2 absolute Wald statistic) in AC (top) and AH (bottom) hepatocytes relative to healthy (pseudobulk; n = 2 libraries per condition). Orange, ExPert top-100; black, predictions among the top-100 DE genes; gray, remaining predictions. Axes are log-scaled; dashed line, y = x. Spearman’s ρ is indicated. (D) Volcano plots of hepatocyte DE in AC (top) and AH (bottom) versus healthy (pseudobulk; n = 2 libraries per condition). Shrunken log2 fold change versus Benjamini-Hochberg-adjusted p-value; dashed lines, |log2 fold change| = 2 and adjusted *p* = 0.05. Black, top-100 DE genes; orange, top-100 ExPert predictions (faded if also DE); gray, other genes. Labeled, the three most downregulated and three most upregulated of the top-100 DE genes (black) and eight ExPert predictions (orange). (E) Pathways enriched among the top-200 ExPert predictions in healthy and ALD (AC and AH combined) hepatocytes (one-sided hypergeometric test; background, all perturbations in the ExPert bank; Reactome and curated mRNA export and splicing gene sets, **Methods**). Shown are mRNA export and mRNA splicing (bold) and the 13 most significant non-redundant pathways, ordered by fold enrichment in ALD. Color, signed log2 fold enrichment (blue, predicted loss of function; green, predicted gain of function); dot size, unadjusted p. (F) As in (E), for the top-200 downregulated and top-200 upregulated DE genes in ALD hepatocytes (background, all tested genes), showing the 15 most significant non-redundant pathways. Blue, enrichment among downregulated genes; green, among upregulated genes. (G) Volcano plots of bulk RNA-seq DE in AC (top) and AH (bottom) versus healthy livers (DESeq2; n = 4 healthy, 6 AC and 6 AH livers; thresholds as in (D)). Genes are colored by ExPert predicted loss-of-function score (z-score across the perturbation bank; gray, z ≤ 0, to orange, z ≥ 3) in snRNA-seq hepatocytes of the same condition; light gray, not scored by ExPert. Labeled, top 200 ExPert predictions in the mRNA export or splicing gene sets. (H) Poly(A) RNA-FISH (hybridization chain reaction) on formalin-fixed, paraffin-embedded (FFPE) human liver sections (5 μm) from healthy individuals and patients with AH or AC 63. Representative Airyscan confocal images of healthy (left), AH (center), and AC (right) livers. Rows, top to bottom: poly(A) RNA; nuclear DNA (dsDNA; DAPI); merge (poly(A), red; DNA, blue); 3× magnification of the boxed region; and the magnified poly(A) signal colored by intensity percentile (scale below; < 80th to ≥ 99.5th percentile). Scale bars, 25 μm (rows 1–3) and 10 μm (rows 4 and 5). (I) Nuclear-to-cytoplasmic ratio of poly(A) foci (log2 scale) in healthy and ALD hepatocytes, colored by condition (n = 84 healthy and 193 ALD [AC, 103; AH, 90] imaging frames from three livers per condition). Frames could not be assigned to individual livers and are treated as independent. (J) Diameter of DAPI-segmented hepatocyte nuclei, colored by condition (n = 44 healthy, 93 AC and 33 AH nuclei from 3 livers per condition). Statistics were computed on per-liver medians (healthy, n = 3; ALD, n = 6 livers). (K) Bulk RNA-seq volcano plots as in (G), with significantly spliced genes (rMATS; FDR < 0.01, |ΔPSI| ≥ 0.1, where PSI is percent spliced in) colored by |ΔPSI|. Circled, genes assayed by RT-PCR in (M). Right, the top 20 ExPert hepatocyte predictions per condition, ordered by rank; filled, the gene’s family contains a significantly spliced member and the gene itself is not differentially expressed; bold, mRNA export or splicing genes. The number of filled genes was compared with 5,000 random 20-gene sets drawn from the perturbations in the ExPert bank that pass the bulk count filter (one-sided empirical test). (L) ExPert predicted loss-of-function scores (z) of mRNA export machinery families in healthy (gray) and ALD (red; mean of AC and AH) hepatocytes. Points, family members; large outlined points, members among the top 200 ALD predictions. Tests are paired over all family members, ALD versus healthy. (M) RT-PCR of alternative splicing for eight exons. Left, gels (lanes in triplicate: healthy, AH, AC); right, ΔPSI (percentage points) relative to the healthy median (n = 3 [biological replicates] per condition; ALD, n = 6). Data are represented as individual replicates with mean ± SD (horizontal line and error bars); bars, median. Boxplots (I, J and L): center line, median; box, interquartile range (IQR); whiskers, most extreme points within 1.5× IQR. *p < 0.05, **p < 0.01, ***p < 0.001; n.s., not significant; by two-sided Mann-Whitney U test ((I), BH adjusted; (J)), two-sided paired Wilcoxon signed-rank test ((L), unadjusted) or two-sided Welch’s t test ((M), unadjusted). See also **Figures S12** and **S13** and **Table S10**.

Across its top-200 calls, perturbation enrichment returned 28 regulatory programs (**Figure 5E, Table S10A**). We focused on two: mRNA export had the highest enrichment and no previous link to ALD, whereas dysregulated mRNA splicing was recently found to be associated with regeneration failure and loss of mature hepatocyte function in ALD^73,75^. Enrichment over the differentially expressed genes instead returned inflammatory and metabolic programs such as complement cascade and neutrophil degranulation, more consistent with downstream injury than with drivers (**Figure 5F, Table S10B**); neither mRNA export nor splicing was among them. Single-nucleus counts are sparse, so a null for the genes named by ExPert could reflect a lack of power rather than a lack of change. We therefore compared these results to deeply sequenced bulk RNA from the same cohort of healthy, AH and AC livers^73^. The genes named by ExPert remained near the origin of the volcano plots, among them nucleoporins, TREX subunits, and SR proteins (**Figure 5G**).

This observation was surprising, so we tested the export prediction experimentally. Efficient export of mRNA from the nucleus is required for cytoplasmic translation, so a loss of export capacity should leave poly(A) RNA in the nucleus. We performed poly(A) RNA FISH on liver sections from healthy individuals and from patients with AH or AC. Poly(A) RNA was enriched in the nuclei of both disease groups and most pronounced in AH (**Figure 5H**), raising the nuclear-to-cytoplasmic ratio from 2.6 in healthy liver to 5.8 in ALD (p < 0.0001; **Figure 5I**). Nuclear diameter was comparable across groups, so the enrichment is not explained by nuclear size (**Figure 5J**). To our knowledge, this is the first evidence of nuclear RNA retention in chronic liver disease, though steady-state FISH cannot exclude contributions from altered RNA production, processing, or turnover.

The gene groups ExPert named were not differentially expressed, so we asked whether they changed at the level of isoform composition rather than total abundance. We ran genome-wide differential alternative splicing on the deep-sequenced liver RNA (rMATS, FDR < 0.01, |ΔPSI| ≥ 0.10) and intersected the result with ExPert’s top-200 calls. Both AH and AC livers carried widespread splicing changes in these genes, with overlapping gene sets between the two conditions, and a majority showed no significant change in transcript abundance, indicating altered isoform usage rather than altered expression (**Figure 5K**). The predicted loss was not spread evenly across the machinery. It separated ALD from healthy nuclei for the nucleoporin complex (*NUP* family, *TPR*, *RAE1*), the *TREX* and *TREX-2* adaptors (*THOC* family, *SARNP*) and the SR proteins, but not for the Ran cycle, the exon junction complex, or the exportins (**Figure 5L**). Interestingly, ExPert’s zero-shot predictions identified three further splicing regulators absent from training: *ESRP2, RBM47* and *A1CF*, all reduced in severe alcoholic hepatitis ^73^. All three ranked in the top 6% of the 10,931 genes absent from training as predicted losses in alcoholic hepatitis hepatocytes (**Figure S13**).

We confirmed these splicing events by RT-PCR in human liver tissue, examining selected mRNA-processing, surveillance, and nucleocytoplasmic export factors (**Figure 5M**). All showed disease-associated splicing shifts in both AH and AC relative to healthy liver, spanning multiple steps of mRNA biogenesis. Among the processing and surveillance factors, *SRSF3*, a splicing factor that also serves as an adaptor for *NXF1*-dependent mRNA export ^76,77^, showed increased inclusion of its 456-nt poison exon 4, which introduces a premature termination codon and targets the transcript for nonsense-mediated decay (NMD)^78–80^. This switch mirrors the increase in poison-exon-containing *Srsf3* transcripts and loss of SRSF3 protein previously reported in dietary mouse models of liver disease^81^, suggesting that productive SRSF3 expression is diminished in ALD. *POLDIP3* (*SKAR*), a component of the exon junction complex, shifted toward skipping of its TDP-43-responsive 87-nt exon 3, an in-frame event that removes 29 amino acids and generates an isoform with greater activity in splicing-dependent translation assays^82^. *UPF3A*, whose exon 4 determines whether it supports NMD^83,84^, produced four variably spliced products in diseased liver.

The splicing changes extended to the transport machinery itself (**Figure 5M**). *XPO1* (*CRM1*), the principal export receptor for cargoes bearing leucine-rich nuclear export signals^85^, increased inclusion of a 64-nt exon in both AH and AC. Several nucleoporins were similarly affected. *NUP54*, part of the central transport channel of the nuclear pore^86^, reduced inclusion of its 55-nt exon, whereas *NUP214*, required for efficient *CRM1*-dependent export^87^, trended toward higher inclusion. *NUP42*, which partners with *GLE1* to promote Hsp70 mRNA export^88^, produced an alternative isoform whose consequence is not established. *AKAP8L* (*HAP95*), which associates with RNA helicase A to facilitate constitutive transport element-dependent export of unspliced RNA^89^, reduced inclusion of a 137-nt exon in both disease groups.

Together, these data reveal that ALD is accompanied by coordinated splicing changes across a network of RNA-processing, surveillance, and export factors. Because the overall mRNA levels of these genes were not significantly altered in AH or AC livers, this layer of regulation was invisible to conventional differential gene expression analysis, yet it was captured by ExPert. We propose that these collective isoform switches dampen mRNA export, providing a potential explanation for the newly observed nuclear accumulation of poly(A) RNA in ALD.

### ExPert recovers affected genetic programs in patients with primary mitochondrial disease

We next asked whether ExPert could identify the affected machinery in a disease with established genetic causes. Primary mitochondrial diseases (PMDs) are a heterogeneous group of genetic disorders that affect 1 in 5,000 live births, with significant neurologic morbidity in childhood, and have known current cures^90^. PMD arises from gene variants that cause mitochondrial dysfunction, including lesions in the mitochondrial electron transport chain (ETC) complexes and their supporting machinery^91^. We profiled balanced PBMCs from four patients with PMD, and five controls with non-PMD conditions by scRNA-seq, then encoded every cell with ExPert and asked which perturbations best explain its state (**Figure 6A**; **Methods**). As in ALD, ExPert’s ranking was uncorrelated with differential expression (Spearman ρ = −0.04; **Figure 6B**). ExPert’s top calls were genes that cause PMD, nearly all scored as loss: mitochondrial ETC subunits, mitochondrial ribosomal proteins, regulators of mtDNA maintenance, and mitochondrial dynamics. Among them were *NDUFAB1* and *NDUFB7* (mitochondrial complex I), *UQCRC1* (mitochondrial complex III), *ATP5F1A* (mitochondrial complex V), and *TFAM* (mtDNA maintenance). The same genes sat near zero by differential expression, paradoxically with a slight bias toward positive fold-change. These patients carry variants expected to impair oxidative phosphorylation (OXPHOS), so the modest increase in OXPHOS-related transcripts would be consistent with compensation rather than with a primary driver of disease. By contrast, ExPert scored those same top genes as strong loss of function, which is consistent with the genetics, biochemistry, physiology, and clinical presentation of these diseases.

**Figure 6.**
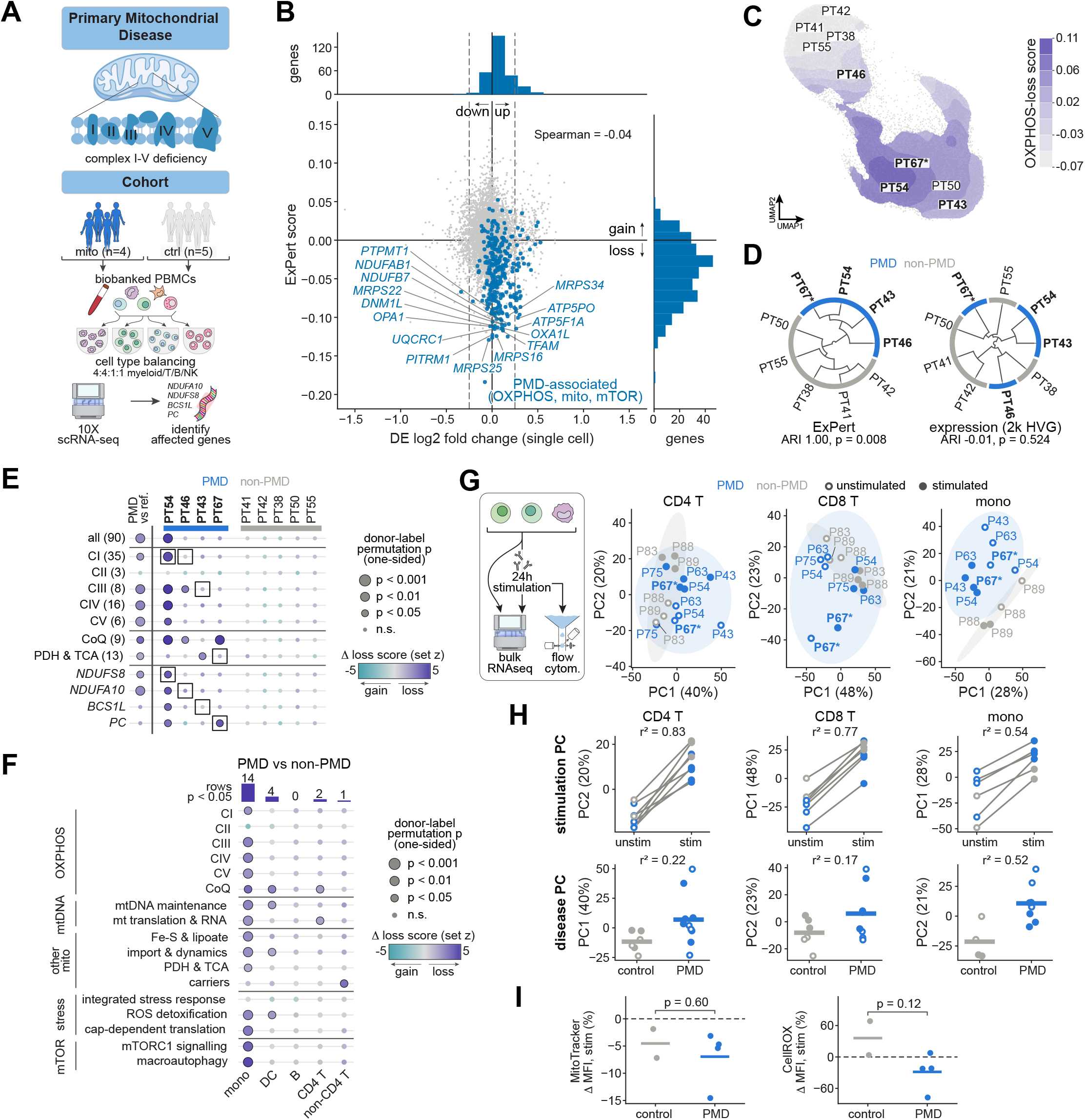
ExPert reveals a monocyte-restricted mitochondrial loss-of-function signature in primary mitochondrial disease. (A) Study design. Biobanked peripheral blood mononuclear cells (PBMCs) from pediatric patients with primary mitochondrial disease (PMD; n = 4) and from reference donors without a PMD diagnosis (non-PMD; n = 5) were balanced by cell type (4:4:1:1 myeloid:T:B:NK) and profiled by single-cell RNA-seq (scRNA-seq; 28,513 cells post-QC). Each patient’s mitochondrial lesion (NDUFS8, NDUFA10, BCS1L, or PC) was provided by the clinical team. OXPHOS, oxidative phosphorylation; complex I–V (CI–CV), respiratory chain complexes I–V. (B) Single-cell differential expression (DE) against ExPert prediction for PMD versus reference monocytes (cells pooled; n = 11,540 cells from 4 PMD patients and 4,271 cells from 5 reference donors), for the 8,615 genes in the ExPert perturbation bank with DE estimates. x axis, log2 fold change (pseudocount 1 on counts per 10,000); dashed lines, |log2 fold change| = 0.25. y axis, ExPert score: cosine similarity between the PMD-versus-reference shift in the ExPert latent space and each gene’s knockout prototype, signed so that predicted loss of function is negative. Blue, PMD-associated genes (295): OXPHOS complexes, mitochondrial biogenesis (mitochondrial translation, protein import, Fe–S cluster biogenesis, cristae formation), mTORC1 signaling, and 44 candidate PMD genes nominated by the clinical team; gray, all other genes. Labeled, the 15 PMD-associated genes with the strongest predicted loss of function. Margins, distribution of the blue genes along each axis. Spearman’s ρ between log2 fold change and ExPert score across all genes is −0.04. (C) UMAP of 15,811 monocytes from all nine donors in the ExPert latent space. Gray, individual cells; shading, OXPHOS-loss score (filled contours of the score averaged over nearby cells): cosine similarity between each cell’s latent embedding and the mean knockout prototype direction of CI, CIII, CIV and CV, averaged over each cell’s 30 nearest neighbors. The score is not centered on the cohort mean; higher values indicate greater similarity to OXPHOS loss. Donor labels mark each donor’s densest region; bold, PMD patients; asterisk, PT67, initially enrolled as a reference donor and reclassified as PMD after re-review of the clinical genetics. (D) Unsupervised clustering of the nine donors (n = 4 PMD, 5 reference) from monocytes in the ExPert latent space (left) and in expression space (right; mean log-normalized expression of the 2,000 most highly variable genes). For each donor, the profile is its mean minus the mean of all other donors; cosine distance, average linkage. Outer ring, donor group (blue, PMD; gray, non-PMD). Agreement between the two-cluster cut and PMD status is given as adjusted Rand index (ARI) with an exact one-sided p-value over all 126 relabelings of four of the nine donors as PMD. (E) Predicted per-donor loss of function of specific mitochondrial targets in monocytes. Rows, from top: all genes in the group rows below (90); OXPHOS complexes CI–CV, coenzyme Q (CoQ) biosynthesis, and pyruvate dehydrogenase (PDH) and tricarboxylic acid (TCA) cycle genes, restricted to diagnostic-grade genes of the Genomics England PanelApp Mitochondrial disorders panel (v10.25)^115^ and grouped by MitoCarta3.0^116^ pathway (gene numbers in parentheses); and each patient’s lesion gene alone. First column, PMD versus reference donors; other columns, individual donors (one profile per donor; n = 4 PMD, 5 reference). Color, loss score: projection of the donor’s latent shift (relative to the cell-weighted centroid of the five reference donors) onto the mean knockout prototype direction of the gene group, in units of the reference donors’ SD (for the first column, PMD mean minus reference mean). Dot size, one-sided *p* value: for the first column, exact donor-label permutation test (126 relabelings); for individual donors, t prediction interval against the five reference donors (4 degrees of freedom). Outlined dots, *p* < 0.05. Boxes, each patient’s lesion gene and the gene group containing it (PC in PT67, confirmed; candidate genes in the other patients). (F) PMD versus reference donors per PBMC cell type, scored as in the first column of (E). Rows, OXPHOS complexes CI–CV (all MitoCarta3.0 subunits and assembly factors), the PanelApp gene groups (defined as in (E)), cellular stress responses and mTOR signaling (Reactome 2024 and curated modules). Color, PMD mean minus reference mean of the loss score (units of reference SD). Dot size, exact one-sided donor-label permutation *p* value (126 relabelings; fewer where donors with < 100 cells of a type are omitted: PT43 from CD4 T cells, PT38 and PT54 from non-CD4 T cells); outlined dots, *p* < 0.05. Top, number of rows with *p* < 0.05 per cell type. CD8 T and NK cells are not shown (< 100 cells in at least three donors). mono, monocytes; DC, dendritic cells; B, B cells; non-CD4 T, unconventional T cells (mostly non-Vδ2 γδ T cells, with naive CD8 T cells; **Figure S14D**). (G) Left, schematic: sorted CD4 T cells, CD8 T cells and monocytes were cultured for 24 h +/-stimulation (CD3/CD28 for T cells, LPS for monocytes) before bulk RNA-seq and flow cytometry (I). Right, principal-component analysis of bulk RNA-seq from an overlapping cohort (PMD, n = 5; controls, n = 3 patients; one library per patient, cell type and condition), unstimulated (open) and stimulated (filled). Variance-stabilized counts (blind; normalization factors from average transcript lengths and size factors estimated across all libraries) of the 1,000 most variable genes per cell type, among genes with mean normalized count ≥ 10. Ellipses, 95% normal-theory regions per group. Six libraries (both conditions of CD8 T cells from P43 and of monocytes from P75 and P83) were excluded for contamination by other cell types after 24 h in culture, leaving four PMD and three control patients for CD8 T cells and four PMD and two control patients for monocytes. Asterisk, P67 (as in (C)). (H) Principal components from (G) for stimulation (top; lines join each patient’s unstimulated and stimulated libraries) and disease (bottom; points, libraries, open unstimulated and filled stimulated; bars, group means). Of PC1 and PC2, the stimulation PC is the one with the higher r² for stimulation and the disease PC is the other. r², variance of the principal component explained by stimulation or by PMD status (linear model over libraries). (I) Change in geometric mean fluorescence intensity (MFI) of MitoTracker Green (mitochondrial mass; left) and CellROX Orange (reactive oxygen species; right) in monocytes after stimulation, relative to unstimulated cells (control, n = 2; PMD, n = 4 patients). Points, patients; bars, group means; dashed line, no change. Monocytes of P75 and P83 were excluded before analysis for T-cell contamination (as in (G)). PMD patients are shown in blue throughout. Statistical tests: exact one-sided donor-label permutation test ((D), ARI; (E), first column; (F)); one-sided t prediction interval against the five reference donors ((E), individual donors); two-sided Student’s t test ((I)). *p* values are nominal (not corrected for multiple comparisons). Spearman’s ρ in (B) and r² in (H) are descriptive (no test). n denotes donors or patients (biological replicates) unless cells are stated. Samples were excluded from (G)–(I) before analysis for contamination by other cell types (CD8 T cells of P43, predominantly myeloid contamination; monocytes of P75 and P83, predominantly T-cell contamination). See also **Figures S14** and **Table S11**.

Scoring each cell for OXPHOS loss placed three PMD donors with known PMD at the high end of the distribution, together with donor PT67, who had been enrolled in the study as a non-PMD control (**Figure 6C**). The placement of PT67 with the PMD patients was unexpected. PT67 presented with severe neurodegenerative disease in the first months of life that had been attributed to a non-mitochondrial metabolic disorder initially. Re-review of the patient’s genetics identified compound heterozygous variants in pyruvate carboxylase (PC), including a known pathogenic variant. PC is localized to the mitochondria and is required for the conversion of pyruvate to oxaloacetate. Although PC deficiency is typically described as a syndrome with three distinct forms, it clearly falls within the broader family of PMDs. PT54, who showed the most severe clinical phenotype (**Table S11**), scored highest by a clear margin. Hierarchical clustering on ExPert scores then recovered the groups exactly (adjusted Rand index (ARI) 1.00, *p* = 0.008), whereas clustering on the 2,000 most variable genes did not (ARI −0.01, *p* = 0.524; **Figure 6D**).

ExPert also identified specific mitochondrial processes affected in each patient. Scored by gene group, every patient showed loss across the mitochondrial ETC complexes, strongest in PT54 (**Figure 6E**). Scored gene-by-gene, the patients’ lesion genes scored as loss in their own monocytes, most clearly PC in PT67, which scored higher for PC than any other donor, and *NDUFS8* in PT54 (**Figure 6E**). PT67’s lesion lies outside the respiratory chain, yet ExPert scored respiratory-chain loss strongly, similar to the three patients carrying disease-driving variants in the mitochondrial ETC complexes. PC deficiency has been reported to produce a secondary mitochondrial complex IV deficiency, in one case leading to an initial diagnosis of PMD second to a mitochondrial complex IV defect before the PC lesion was found^92^, suggesting that ExPert may also identify secondary pathologic consequences in PMD.

Myeloid cells have been shown to be required for disease progression in mouse models of PMD^93,94^. Interestingly, ExPert’s OXPHOS-loss signal was confined to monocytes. Comparing PMD with non-PMD for each cell type, 14 of the scored sets differed in monocytes, against four in dendritic cells, two in CD4^+^ T cells, one in non-CD4^+^ T cells, and none in B cells (**Figure 6F**). The sets affected in monocytes spanned the mitochondrial ETC complexes, mtDNA maintenance and translation, ROS detoxification, and mTORC1 signalling. Mitochondrial defects can be silent at rest and emerge under metabolic demand, so we purified CD4^+^ T cells, CD8^+^ T cells, and classical monocytes from five PMD patients, three of them from the initial cohort, and three additional controls (**Figure 6G**). We cultured them for 24 hours with or without immune stimulation and performed RNA-Seq to determine whether the PMD patients’ responded similarly. Stimulation dominated the principal axis in both T cell populations (r^2^ = 0.83 and 0.77) and less so in monocytes (r^2^ = 0.54). Disease status separated the donors most strongly in monocytes (r^2^ = 0.52, against 0.22 in CD4^+^ and 0.17 in CD8^+^ T cells; **Figure 6H**). Next, we measured mitochondrial content and reactive oxygen species levels directly in monocytes by flow cytometry (**Figure 6I**). Mitochondrial mass change upon stimulation was comparable between groups (-6.9% against-4.5%, *p* = 0.60), while reactive oxygen species induction trended lower in PMD patient cells (-28.5% against +36.0%, *p* = 0.12).

Altogether, ExPert identified the affected respiratory-chain complex, in some patients the lesion gene itself, and suggests a cell-type specific impairment in patients with PMD. The model correctly identified respiratory chain dysfunction in PT67 even though a PMD was not diagnosed at the time of sample collection. Across all patients, the altered mitochondrial signal came largely from monocytes, which is consistent with mouse data and a novel finding in human disease^93,94^. Patients with PMDs have identified monocyte-associated inflammatory and interferon transcriptional signatures, and changes in monocyte mitochondrial respiratory-chain activity, suggesting that innate immune dysregulation may contribute to the systemic manifestations of mitochondrial disease^95–97^. Notably, previous human studies in mtDNA-driven PMDs have identified myeloid-specific accumulation of dysfunctional mtDNA^98,99^. The consistent monocyte signal suggests myeloid cells may be uniquely sensitive to mitochondrial dysfunction in PMD, regardless of which genome causes disease. Lastly, the strength of the ExPert respiratory-chain signal tracked clinical severity, suggesting ExPert may recover prognostic information about PMD patients’ disease course.

## DISCUSSION

Considerable effort has gone into predicting how a cell responds to a perturbation. Here we asked the inverse question, which perturbation produced an observed state, and built ExPert to answer it. Trained across 22 CRISPRi screens spanning 12 cellular contexts, 19.5 million cells, and 61,445 perturbation–context pairs covering 8,964 perturbations, ExPert more than doubled held-out-context recall over the best existing forward or inverse method at a fraction of their compute, and extended to 19,895 genes, most never seen by the model in training. The same prototype table recovered annotated drug targets, degron-induced protein loss, and combinatorial perturbations it was never trained on. Applied to two diseases, ExPert implicated genes which differential expression did not: isoform-level remodeling of mRNA export in alcohol-associated liver disease, confirmed by poly(A) FISH and RT-PCR, and respiratory-chain complexes in PMD, concordant with genetics and clinical presentation of PMD.

ExPert’s learned prototype embedding was built for inference but may be useful for the forward problem of perturbation response prediction. In principle, the prototypes could be supplied to a forward model in place of a text or annotation embedding, or the reconstruction objective could be reframed during training so each prototype also generates the state it names. Other extensions depend on data that does not yet exist at scale. Variational generative models already combine transcriptome and chromatin accessibility per cell^100^, and the same encoder could take both, but multimodal and spatial CRISPR screens are not yet available at scale, though both are expanding rapidly^17,101–103^. Our model recovered combinations without explicitly training ExPert on combinatorial data; as those data expand, the model could be trained on the combinations directly. Furthermore, ExPert was trained only on CRISPRi knockdown, yet recovered CRISPRa perturbations, albeit at lower accuracy. Activating and repressing perturbations do not produce symmetric responses, and the two carry complementary information about gene function^104,105^, so emerging large-scale CRISPRa data will improve the biological grounding of the model and its generalizability.

In both drug screens and disease samples, ExPert recovered biology that annotations did not contain. It highly ranked *GSK3* for abemaciclib (**Figure 4G**), an off-target the compound’s label omits. In PMD, it scored one of six participants enrolled as a control with the disease group, and re-review of that patient’s genetics reclassified them as PMD, so the discordance reflected an error with the label that ExPert corrected. The respiratory-chain score also tracked clinical severity, which a larger cohort could test as a prognostic marker. Any disease whose drivers are unknown can be queried the same way, since the procedure needs only a transcriptomic cell state and ExPert’s reference prototype table. In principle, the same prediction could be applied to patient tissue to select the most strongly affected cells, where no measured efficiency is available. What limits ExPert’s performance is the extent to which a perturbation’s response is shared between the screened contexts and the patient’s cells. Held-out donors were the hardest split in our benchmark, and performance along this axis should improve as Perturb-seq data from larger donor cohorts accumulate.

ExPert ranks perturbations whose transcriptional signature matches a query. Rank is not proof of causality, and in disease settings the evidence that the top calls are causal must come from outside the model: DNA sequencing in PMD, and the FISH and splicing data in ALD were used herein. Decomposing diseases arising from many simultaneous lesions, such as cancer, may require combinatorial training data, though constrained optimization over inferred networks offers another route to expressing a disease state as a combination of perturbations^106^. ExPert also returns ranks rather than calibrated probabilities, which score-based diffusion models that infer perturbation identity probabilistically could supply^107^.

Practically, the ranking converts an unexplained cell state into a short list of experiments: in both diseases here, the top calls—out of 19,895 candidates—were few enough to test directly, and those tests found biology which differential expression had missed. As perturbation atlases expand to more contexts and donors, more of each response becomes identifiable as shared, and these rankings will only improve.

## AUTHOR CONTRIBUTIONS

Y.G. and L.S. conceived and designed the study with input from B.K.S. and E.P. L.S. designed and performed a majority of the computational experiments and analyses with input from Y.G., E.P., and B.K.S. B.K.S. initiated the conceptual discussion on tackling the inverse problem. C.W.J., H.S., B.K., R.A.G., E.R., D.S.M. and S.E.W. performed PMD sample collection and experiments. D.D., B.P., Z.S., A.M.D. and A.K. performed ALD experiments and splicing analysis. T.B. and M.A. performed degron experiments. E.I.G., J.L.S., and G.P. prepared the figures and tables, manuscript preparation, and provided feedback on study design. J.L.S., I.S., and A.M.A. contributed to the analyses and data curation. J.J. assisted in data curation and manuscript preparation. C.M. assisted L.S. and Y.G. in formulating the computational framework. N.S.C. contributed to assessing the model on PMD. L.S. and Y.G. wrote the manuscript with input from all authors.

## ACKNOWLEDGEMENTS

We thank members of the Goyal lab for helpful discussions and comments on the manuscript. Sequencing was performed by the Center for Synthetic Biology Biofoundry Core Facility (RRID:SCR_026869), which is supported by the Northwestern University Office for Research. We thank the Feinberg Information Technology, Northwestern University Information Technology, and Quest High Performance Computing Cluster at Northwestern University Feinberg School of Medicine for computational support and services. YG thanks the National Science Foundation through the Center for Living Systems Ignite Award (grant no. 2317138) as well. This work was supported by the National Institute for Theory and Mathematics in Biology (NITMB) through the National Science Foundation (NSF; DMS-2235451) and the Simons Foundation (MPTMPS-00005320). B.K.S. was supported by the Northwestern University Clinical and Translational Sciences Institute (NUCATS) T32 program, NITMB, Biohub SPARK award, and Northwestern University startup funds to Y.G. L.S., H.S., and J.J. were supported by funding to Y.G. J.L.S., E.I.G., and G.P. received support from NITMB. Individual fellowship support was provided by the National Institute of Allergy and Infectious Disease of the National Institutes of Health fellowship (F31AI189214)(to E.I.G.), the QCMDB T32 fellowship (to H.S.), the Biotechnology Training Program T32 fellowship (to G.P.), the Northwestern MSTP T32 fellowship (T32GM144295)(to J.L.S.), Schmidt Science Fellows, in partnership with the Rhodes Trust (to R.A.G.), and the Charles F. Kade Fellowship (to D.D.). Y.G. acknowledges grant support from the Pew-Stewart Scholars Program, American Heart Association, Biohub Investigator Award, and Burroughs Wellcome Fund Career Awards at the Scientific Interface. Y.G. acknowledges gift from Schmidt Sciences to support this work. Y.G., A.K., and N.S.C. are Biohub Investigators. Y.G. is a Senior Fellow of Biohub. A.K. acknowledges support from National Institutes of Health (R01AA010154 and R01HL126845), Muscular Dystrophy Association (MDA1072487), and Biohub Investigator Award. The PMD studies were supported by Northwestern Center for Human Immunobiology Pilot Award, Cooperative Centers on Human Immunology (CCHI) Infrastructure and Opportunity Fund (IOF), the National Institute of Neurological Disorders and Stroke R21NS149367, (S.E.W, D.S.M), and the Hartwell Foundation Individual Biomedical Research Award (S.E.W, C.W.J). C. Marr received funding from the European Research Council under the European Union’s Horizon 2020 Research and Innovation Programme (grant agreements 866411, 101113551, and 101213822) and support from the Hightech Agenda Bayern. This work was supported through the Bavarian AI Foundation Model Initiative, funded by the Bavarian State Ministry of Science and the Arts (StMWK). Any opinions, findings, and conclusions or recommendations expressed in this material are those of the author(s) and do not necessarily reflect the views of the National Science Foundation and other grant agencies.

## AI USE

During the preparation of this work, the authors used Anthropic’s Opus 4.8, 5, and 5.5 and Sonnet 4.6 via Claude Code to write and edit code, and via Claude Chat to brainstorm, search the literature, and proofread the manuscript. The authors reviewed and edited AI-generated content as needed and take full responsibility for the content of the published article.

## DATA AND CODE AVAILABILITY

All source datasets generated and analyzed in this study are publicly available from their original publications or this study, listed with accessions in **Table S1A**. The trained ExPert model and its prototype table are available on Hugging Face upon publication.

All original code is publicly available as of the date of publication, distributed across three repositories:

● **ExPert** (github.com/GoyalLab/ExPert) — the main package: model, training and inference code.
● **ExPert-data** (github.com/GoyalLab/ExPert-data) — the store and database construction pipeline, including vocabulary construction, quality control, effect-size estimation, gene panel selection and benchmark fold generation.
● **ExPert-benchmarks** (github.com/GoyalLab/ExPert-benchmarks) — the benchmarking workflow for **Figure 2**, the comparison-method adaptations and the scoring code.

Any figures shown in this study have been reproduced fully using the published code and data. Any additional information required to reanalyze the data reported in this paper is available from the lead contact upon request.

## METHODS

Training a single model across heterogeneous perturbation screens required a common representation of gene expression and perturbation identity. We therefore harmonized 22 single-cell CRISPRi datasets, standardizing gene identifiers, perturbation labels, and quality-control criteria. The resulting database contained 31.8 million cells after filtering and control subsampling, of which 19.5 million perturbed cells spanning 8,964 perturbations formed the pool used for model training and evaluation. Below, we describe database construction and benchmark splits, the ExPert model and its training procedure, and the evaluation of perturbation retrieval across methods.

### Training database construction

#### Data sources

We assembled 22 published Perturb-seq datasets from eight studies: ReplogleWeissman2022^19^ (K562 genome-wide and essential, RPE1), ReplogleWeissman2025^42^ (HepG2, Jurkat), Marson2025^20^ (twelve datasets spanning four donors and three stimulation conditions in primary human CD4⁺ T cells), XAtlas2025^43^ (HCT116, HEK293T), Mali2026^45^ (KOLF2.1J), Wong2026^44^ (TeloHAEC) and Zhuang2026^41^ (hESC). Datasets are published in a machine-readable datasheet that records the source URL, assay metadata, cell line, donor and condition annotations, and the axis declarations used for benchmark fold generation.

Of the 49,555,490 cells in the raw corpus, 31,789,072 (64.1%) were retained after filtering and control subsampling. The table below summarizes the contribution of each exclusion category and the final retained cell count.

| **Filter** | **Cells** | **% of raw** |
|---|---|---|
| Unassigned guide (no perturbation label) | 11,517,431 | 23.2% |
| Control cells beyond the 10,000-per-dataset cap | 1,499,680 | 3.0% |
| Quality control (MAD outlier and mitochondrial) | 4,749,307 | 9.6% |
| Multiple perturbations per cell | 0 | 0.0% |
| **Retained** | **31,789,072** | **64.1%** |

Missing perturbation assignments accounted for the largest reduction and occurred exclusively in the Marson2025 series, where 34.9–42.9% of cells in each dataset lacked a perturbation label. These cells could not provide supervised perturbation labels and were excluded. The ten datasets outside this series retained 13,775,783 of 15,945,019 cells (86.4%); thus, the overall retention rate primarily reflected the availability of guide assignments in the twelve T-cell datasets. Restricting the corpus to cells annotated with a single perturbation removed no additional cells, as the source datasets were single-guide by construction or did not annotate multiplets.

### Gene vocabulary and identifier harmonization

All datasets were mapped onto a single append-only vocabulary of 38,638 genes keyed on Ensembl^117^ gene identifier, formed by taking the union of the feature axes of the raw datasets.

We aligned genes across datasets using stable Ensembl gene identifiers rather than gene symbols. Keying on Ensembl identifiers rather than gene symbols is required by the source data: 313 Ensembl identifiers carry conflicting symbols across datasets (for example *SARS/SARS1*, *KARS/KARS1*, *ADPRHL2/ADPRS*), because the source studies used different GENCODE ^118^ releases. Under symbol keying, each of these splits into two largely empty columns and no error is raised. Three datasets provide no gene-identifier column; for these, symbols were resolved through a 39,108-entry symbol-to-Ensembl map extracted once from the datasets that provide both identifiers. This map is used for annotation lookup only, and no expression data crosses datasets.

### Quality control

Cells were filtered with a median-absolute-deviation (MAD) outlier protocol computed from per-cell scalars^119^. A cell was discarded if it was an outlier at five MADs on any of the log-transformed total counts, log-transformed detected genes, or the fraction of counts in the top 50 genes; an outlier at three MADs on mitochondrial fraction; or above an absolute mitochondrial fraction of 20% for cancer-derived lines or 12% otherwise, assigned from the datasheet. Mitochondrial gene membership was derived from the Ensembl vocabulary rather than from per-dataset symbol flags, which are inconsistent across studies. Only passing cells were stored; the per-cell metrics were retained so that alternative thresholds can be re-derived without reading the expression matrix again.

### Control cells

Up to 10,000 control cells per dataset were sampled, retaining 189,365 in total. The sampled barcodes are stored rather than re-drawn at each build. Sampling without replacement depends on the length and ordering of the candidate array as well as on the seed, so a draw is not reproducible from the seed once the input set changes: with the seed fixed, a 5% change in pool size produced only 2,018 of 10,000 overlapping barcodes. Because the control draw propagates into the QC thresholds and from there into every downstream selection, the barcodes are stored.

### Perturbation effect size

Each perturbed cell was assigned a continuous perturbation-effect-size score with a custom Python implementation of Mixscale^120^, restructured so that global fits are computed once per dataset and pinned. For each dataset, we selected the top 3,000 highly variable genes from stored per-dataset statistics and pinned the normalization and per-gene scaling; fitted a 40-component principal component basis on control cells only; and scored each perturbed cell by projecting it into that basis and subtracting the mean of its 20 nearest control neighbors. Perturbations with fewer than five differentially expressed genes received a score of 0.0. Highly variable gene statistics were computed with a streaming reimplementation of the *seurat_v3*^121^ procedure, verified to give results identical to the reference implementation ^121^.

We persist the raw signed z-score rather than a rescaled value, deferring any scaling to selection time. This preserves both the sign and the tail: per-dataset maxima range from 26.5 to 96,199 against medians of 0.3 to 2.3, reflecting a near-zero-standard-deviation blow-up that is visible and correctable downstream but would be silently clipped by saturating at write time. In total, 28.2 million of 31.8 million cells (88.8%) received a score; the remainder are controls or belong to perturbations below the differential-expression threshold.

### Gene panel selection

The input panel was selected to maximize perturbation separability rather than overall variability. Within each training dataset, cells of each perturbation were split at random into two pseudobulk replicates over the full 38,638-gene vocabulary, with perturbations of fewer than 20 cells giving a single profile. Profiles were normalized to counts per million, log-transformed and expressed relative to the dataset’s control. Each gene measured in a dataset was scored by the ratio of between-to within-perturbation variance across replicates (an unmoderated F-statistic), and the panel is the union of each dataset’s 500 highest-scoring genes (3,096 genes).

### Separability evaluation

Perturbations within each dataset were split at random into two halves. Genes were selected on one half, and each replicate of the other half was matched by cosine similarity, over the selected genes, to reference profiles of all scored perturbations, each reference being the cell-weighted mean of that perturbation’s remaining replicates. Separability is the fraction of replicates whose own perturbation ranks first, averaged over the 22 datasets.

### Compared gene sets

At each size n from 100 to 5,000 genes per dataset we compared (i) the union of each dataset’s top n by F-statistic, the construction used here; (ii) the union of each dataset’s top n highly variable genes, computed per dataset on raw counts of quality-controlled cells by a streaming reimplementation of Scanpy’s seurat_v3^121^ flavor, identical in output, and ranked by standardized variance, with ranks available only for the top 3,000 per dataset; (iii) each dataset scored on its own top n alone, an upper bound that is not deployable because it requires knowing a query’s dataset of origin; (iv) the top n genes measured in every dataset, ranked by mean per-dataset F-statistic percentile; and (v) random genes matched in number to (i). Variance moderation, expression dependence, target-gene masking, and a fold-wise rebuild in which each held-out dataset was rescored under a panel built from that fold’s training datasets alone, are reported in **Supplementary Note 2** and **Figure S1F**.

### Perturbation pool and database composition

A perturbation entered the shared pool if it had at least 10 cells in a dataset of at least 60% of the voting groups (6 of 10) and at least 50 cells after pooling. Voting groups were defined by screen: each Replogle and X-Atlas screen formed its own group (K562 genome-wide, K562 essential, RPE1, HepG2, Jurkat, HCT116, HEK293T), and the Marson2025 datasets formed one group per stimulation condition (resting, 8 h, 48 h; four donors pooled). The single-dataset studies Zhuang2026, Wong2026 and Mali2026 did not vote, but their cells were retained for every pooled perturbation. This yielded 8,964 perturbations. Only single perturbations were retained. Letting all 13 groups vote would have produced 8,652 perturbations, sharing 8,380 with the shipped pool (+272 / −584).

Perturbed cells whose perturbation the pool excluded were written to a separate zero-shot evaluation set, capped at 10 cells per dataset-and-perturbation group.

The resulting database comprises 19,513,869 perturbed cells and 189,365 control cells over 3,096 genes, stored as CSR count matrices with per-cell annotation for dataset, perturbation, label, control status and scaled efficiency. Three subsets are derived from the same build and used for benchmarking: a 1,157-perturbation subset of 2,741,678 cells, a 200-perturbation subset of 671,242 cells, and a zero-shot set of 683,651 cells carrying the 9,874 perturbations with no training cells.

Because each dataset measures only a subset of the vocabulary and the panel is a union, the database carries two per-dataset boolean layers: the genes a dataset measured, and the genes the separability selection assigned to it.

### Benchmark fold generation

Folds are declared in the datasheet rather than inferred from dataset names. Any datasheet column naming a holdout axis defines one, and each distinct value becomes a fold in which every cell carrying that value is held out. Three axes are declared: cell line, donor, and condition.

Each donor fold additionally holds out one cell line, which is consumed from the cell-line axis so that no context is held out twice. Pairing is inverse by size, so that the largest donor takes the smallest cell line and donor folds are not trivially small, and no fold removes more than 35% of cells. This yields twelve folds: four donor-and-cell-line folds, five cell-line folds and three condition folds (**Table S5C**).

Within each fold, the remaining cells were partitioned into training, validation, and in-distribution test sets, with validation and test capped at 10 cells per dataset-and-perturbation group and no cap on the number of perturbations, so that every perturbation is represented in evaluation and no single large dataset dominates it. All remaining cells become training data. Each fold’s four index sets are written as positional indices, with an assertion that they partition every cell exactly. Split generation used seed = 42, test_frac = 0.1, eval_cap = 10, min_cells_per_perturbation = 50, min_cells_per_class = 10 and min_dataset_frac = 0.6.

Folds were generated for the full database and for each benchmark subset. The 1,157-perturbation subset inherits the fold-to-context assignment of the full split set shown above (parent_splits: full, id_split_mode: restricted); The 200-perturbation subset was split independently. Because the subset changes relative dataset sizes, the inverse-by-size pairing assigns D1 to HCT116 rather than TeloHAEC, which appears here as a standalone cell-line fold (**Table S5C**).

### Implementation

The pipeline is a Snakemake^122^ workflow in which no stage loads a complete expression matrix; all primitives operate on row bands of CSR matrices read directly from HDF5, so that memory scales with chunk size rather than with dataset size. Peak resident memory for the heaviest stage is 12.5 GB across all 22 datasets and is close to flat in dataset size (5.2 GB at 545 million non-zero entries; 12.5 GB at 25.4 billion, a 47-fold increase in data for a 2.4-fold increase in memory). Total build time for the final database is 1 h 24 min. Artifacts are content-addressed under two independent configuration hashes, separating per-dataset parameters from the dataset list, so that adding a dataset reuses all existing per-dataset artifacts. Every stage is deterministic given the configuration and seed, with the exception of the control draw, which is why sampled barcodes are stored rather than re-derived. Each stage has a validation gate run against real rather than synthetic data; streaming highly variable gene selection was verified to agree exactly with the reference implementation, and parallel and serial materialization were verified bit-identical.

### Input representation

The model receives a cell’s expression profile over the gene panel and nothing else. Each profile was centered on the mean control profile of its own dataset before encoding, removing the between-screen mean shift without a model. Residual dataset-specific variation, including overdispersion, dropout rate and library-size distribution, is absorbed by a decoder conditioned on the dataset of origin^123^.

Let *x*_i_ ∈ ℝ^G^ be the raw counts of cell *i*, from dataset *d*_i_ with control cells *C*_d_. The encoder receives log-transformed counts centered on the control mean of that dataset:

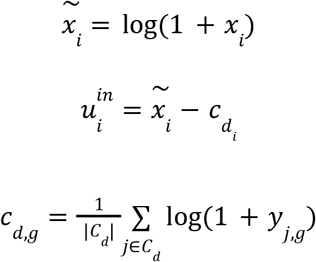

where *y*_j,g_ are the raw counts of control cell *j*. During training the centered input is multiplied element-wise by

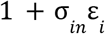

with *ε*_i_ ∼ N(0, *I*) and *σ*_in_ the input noise scale. We apply no library-size normalization to the input, and the reconstruction target remains the raw counts, so centering changes what the encoder sees, not what the decoder has to reproduce. Values for every symbol in this section are given in **Table S4**.

Because the correction is measured rather than fitted, a screen absent from training needs nothing re-estimated before it can be scored. We compute its control mean from its own control cells, or from all available query cells with no control group, requiring at least 10 cells.

### Model architecture

#### Backbone

ExPert uses a variational autoencoder (VAE) backbone in the style of scVI^123^, implemented on the basis of scvi-tools^124^, and extended into a perturbation retrieval model.

#### Encoder

A single projection block maps the *G*-dimensional centered input to *h* hidden units (linear, layer normalization, GELU, dropout at rate *p*_drop_, layer normalization), followed by one residual block at *h* and a closing layer normalization. Gaussian noise of standard deviation *σ*_h_ is added to the hidden representation during training. Two linear heads give the posterior mean *μ*_i_, of dimension *d*_z_, and the log-variance, whose exponential *σ*_i²_ is floored at *ε*_σ_. The latent is sampled during training as

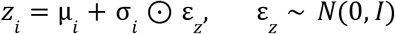

drawn independently of the input noise ε*_in_*, while evaluation and inference use *μ*_i_. No covariate enters the encoder. We used *h* = 4,096 and *d*_z_ = 2,048.

#### Decoder

The decoder receives *z* concatenated once at the input with *E_di_*, the embedding row of that cell’s dataset, where *E* is an orthogonally initialized learnable embedding with one row per dataset and width *d*_ctx_. Dataset identity is the only covariate, and it enters only the decoder. A projection block reduces the concatenated input to *h*_dec_ units, followed by one residual block (layer normalization and GELU throughout, no dropout). Three heads produce, for each gene *g*, expression proportions *ρ*_i,g_ from a softmax over genes, zero-inflation logits *π*_i,g_, and a gene-wise inverse dispersion *θ*_g_ shared across cells. The mean of the negative-binomial component is

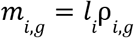

with *l*_i_ the summed counts of cell *i* as its observed library size, so library size is an exact size factor rather than a learned quantity; the mean of the full zero-inflated distribution is (1 − sigmoid(*π*_i,g_)) *m*_i,g_.

#### Latent centering

Because the prototypes receive no dataset information, any per-dataset offset surviving input centering would be fit by the alignment objective. Latents were therefore centered per dataset on the training side, using an exponential moving average of each dataset’s latent mean, so the correction costs no extra pass over the data. During training a running mean per dataset is maintained,

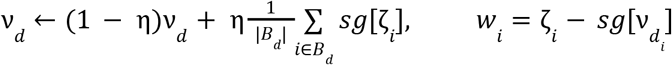

where *η* = 0.05 is the running-mean rate, *ζ*_i_ = Dropout(*z*_i_) at the same rate *p*_drop as the encoder, *B*_d_ is the set of cells of dataset *d* in the batch, and sg denotes stop-gradient. All alignment-side terms and all readouts operate on *w*_i_. At evaluation,

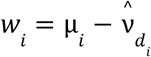

with *ν^*_d_ the stored running mean for datasets seen in training; for a dataset never seen, *ν^*_d_ is the mean latent of its control cells if present and otherwise of all its cells. Reconstruction always decodes the uncentered *z*_i_, so the generative term still sees the full profile.

#### Perturbation prototypes

Each perturbation is represented by one learnable prototype in the centered latent space, and retrieval is a comparison against those prototypes. Each of the 19,895 vocabulary genes owns a row of dimension *d*_z_ in a prototype table *A*, written *a*_p_ for perturbation *p*, initialized i.i.d. from a zero-mean Gaussian of standard deviation *σ*_A_. Only the *L* = 8,964 rows of trained perturbations receive gradients; the remaining rows are never written by the alignment objective and serve as the address space for prior-derived placements and negatives (Prior knowledge as neighborhoods, below). Prototypes live directly in the centered latent space without an adapter, so the readout is a single inner product.

#### Proxy Anchor alignment

Cells are placed relative to prototypes with the Proxy Anchor loss^49^, weighted by *λ*_PA_, using the learnable prototypes as proxies. Let *V* be the cells of a batch carrying a trained perturbation, *y*_i_ the perturbation of cell *i*, and *c*_i_ its effect-size weight (next section). The weight enters twice. It scales the gradient into the cell, through

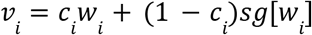

whose forward value is *w*_i_, and it weights that cell’s term in the loss. Cell representations are centered on the batch mean and prototypes on the prototype mean,

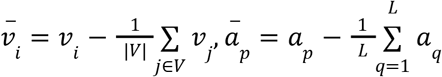

giving cosine similarities *s*_ip_ = cos(*v̄*_i_, *ā*_p_). With scale α = 32 and margin δ = 0.1,

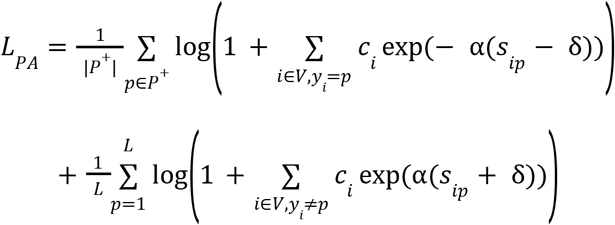

where *P*⁺ is the set of perturbations with at least one cell in the batch. The negative term is a soft maximum over negatives only, so repulsion from wrong prototypes continues after the correct prototype already wins. A margin-relaxation variant and an auxiliary softmax classification head were each tested at full scale and each reduced held-out retrieval; neither is used, and the classification head of the shipped model is untrained.

#### Perturbation effect size

Each cell carries a measured effect size *e*_i_ (Mixscale^120^; see above). Effect sizes are not comparable across screens, with per-dataset medians spanning 0.108 to about 3.0, so a global scale would largely encode which screen a cell came from. We therefore used the within-dataset rank of |*e*_i_| among the cells of that dataset in the batch,

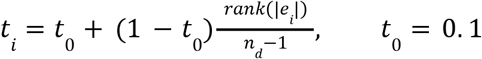

with ranks counted from 0 for the smallest |*e*_i_|, *n*_d_ the number of cells of dataset *d* in the batch, and *t*_i_ = 1 where *n*_d_ < 2, giving a weight in [0.1, 1]. This *t*_i_ is the per-cell weight *c*_i_ in the Proxy Anchor loss above, which is the load-bearing use of the effect size during training.

The same quantity is also predicted. A head *g*_φ_, an MLP of 2,048 to 1,024 units with layer normalization and GELU and a scalar output, regresses *t*_i_ from a detached copy of the cell representation,

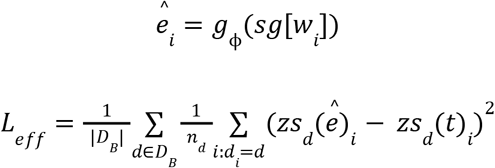

where zs_d_ is a z-score within dataset *d* with the standard deviation floored at 10⁻⁴, and *D*_B_ is the set of datasets in the batch with *n*_d_ ≥ 8 and a non-degenerate target. Up to that floor the loss equals

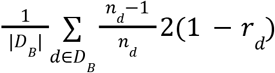

with *r*_d_ the per-dataset Pearson correlation between *ê* and *t*, so it fits the ordering within each screen rather than any absolute scale. Because the input is detached, this term trains the head alone and does not shape the latent. At inference the head’s raw output is one input of the effect-size readout (below).

Effect sizes are not imputed. Cells whose effect size is non-finite sort above every finite value and so take the highest ranks within their dataset in the batch, giving them weights at or just below 1, in arbitrary order among themselves, in both the Proxy Anchor loss and the head’s target. Ranks use arbitrary tie-breaking with no tie averaging.

#### Prior knowledge as neighborhoods

Prior banks supply which genes a perturbation resembles, never where its prototype should sit, so every coordinate in the table remains a combination of prototypes learned from expression.

Three banks enter the shipped model: language-model embeddings of gene descriptions (qwen3-embed-8b^58,59^, 4,096-d, covering all 19,895 rows), STRING interaction profiles^51,52^ (256-d, 93.0% of rows) and DepMap co-essentiality^46,47^ (1,178-d, 88.3%). A bank *b* covers a row where it has a profile for that gene, and *C_b_* denotes the rows it covers.

#### Which prototypes are pulled

The evidence behind a trained prototype is summarized as

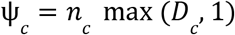

with *n*_c_ the number of training cells of perturbation *c* having a finite effect size and *D*_c_ the number of datasets contributing at least one such cell, and converted to an upward rank fraction over the *L* trained perturbations,

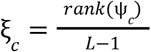

The gate is hard and serves only to define two sets: the prototypes at or below the *q*-th quantile of *ξ* form the pulled set,

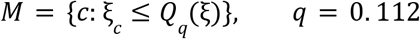

and the fit set *S* is the complement of *M* among the *L* trained prototypes, giving 1,004 pulled and 7,960 fit prototypes. Targets are built from fit prototypes only, so no pulled prototype contributes to its own or any other target. The trained prototypes are also split into five random folds *F*₁,…,*F*₅, and a pulled prototype in fold *F*_f_ draws its neighbors from *S*_f_, the fit prototypes outside *F*_f_. This cross-fit is not needed to keep a prototype out of its own target, which the disjointness of *M* and *S* already guarantees; it is retained from the general implementation, in which the pulled and fit sets may overlap, and its only effect in the shipped model is that each target is built from about four fifths of the fit set.

#### Targets

Each bank is centered on the rows it covers and rescaled to unit mean row norm,

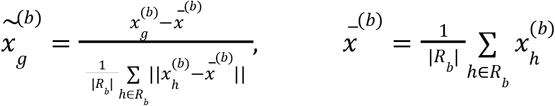

For a pulled prototype *c* ∈ *M* ∩ *F*_f_ covered by bank *b*, the neighborhood *N*_b_(*c*) is the *k* = 25 rows *j* ∈ *S*_f_ ∩ *R*_b_ whose centered bank profiles have the highest cosine similarity to that of *c*. They are weighted by their similarity clamped at zero,

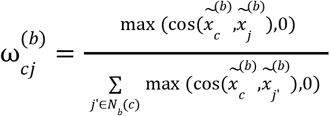

or uniformly by 1/*k* where all similarities are non-positive. The bank’s prediction is the weighted mean of the neighbors’ unit-normalized prototypes,

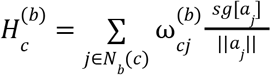

where *a*_j_ is the learned prototype of perturbation *j*. The banks are then fused over those covering row *c*. The text bank covers every row, so at least one bank always applies, and rows missing from STRING or DepMap are fused over fewer banks. The fusion uses three normalizations, each for a different reason. Each bank’s prediction is unit-normalized first, so that the banks enter with equal weight regardless of how concentrated their neighborhoods are. The fused sum is unit-normalized again, because only its direction is a target. That direction is finally scaled to the median prototype norm over *S*, so that the target lies at the scale of the learned prototypes:

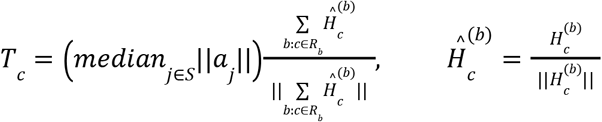

#### Prior pull

Pulled prototypes present in training are rotated toward their targets by

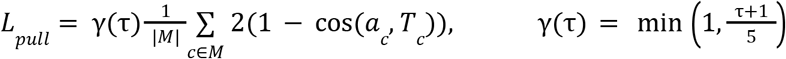

weighted by *λ*_pull_ = 3.0. Here *τ* is the zero-based epoch, so *γ* takes the values 0.2, 0.4, 0.6 and 0.8 over the first four epochs and stays at 1 from the fifth epoch onward. Within *M* every prototype has the same weight (see above). The loss depends on cosine alone, so its gradient has no radial component: prototype norms are untouched, and the scale of *T*_c_ does not affect the loss. Because *T*_c_ carries no gradient, the pull trains prototypes only, and prior structure cannot propagate into the representation of a cell. Because the pull competes with the alignment term rather than overriding it, a gated prototype settles between its cells and its neighbors.

#### Unseen perturbations

A gene with no training cells has no evidence and no learned prototype to blend with, so the target construction above is applied outright and the row is set to the result, *a*_u_ = *T*_u_. An unseen row belongs to no fold and is not gated; its neighbors are drawn from the fit-set prototypes of the first fold, *S*₁, and its placement is scaled to the median prototype norm over *S*. The fill covers all 10,931 untrained vocabulary rows, extending the addressable vocabulary from 8,964 to 19,895 gene perturbations with no retraining and no additional data. The shipped fill was refit after training on the final prototypes with the same construction, except that neighbors are weighted by max(cos, 0)⁴ instead of max(cos, 0); pull targets keep the linear weighting.

The filled rows are also used during training as negatives, under the linear-weighted fill, which is refit each epoch. At each step a fresh set *J* of 256 filled rows is sampled and pushed away from the batch *V* by

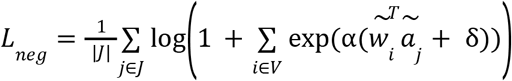

where *w̃*_i_ is the cell’s centered latent, centered on the batch mean and unit-normalized, and ã_j_ is the filled row with a stopped gradient, centered on the mean of the sampled set and unit-normalized:

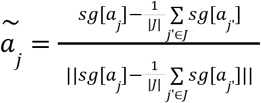

*α* = 32 and *δ* = 0.1 are the Proxy Anchor^49^ scale and margin. There is no positive term, no effect-size weight and no ramp. Because *a*_j_ for a filled row is built from stop-gradient prototypes and is itself detached, the gradient reaches the encoder only and leaves both the filled rows and the prototypes they were built from unchanged. A perturbation that is plausible but absent from the batch is therefore something the encoder is trained to reject, rather than a row that happens never to have been penalized.

#### Generative loss

The generative term is the evidence lower bound (ELBO) over the *G*-gene panel on raw counts, averaged over a batch *B* of cells (as described in scVI^123^),

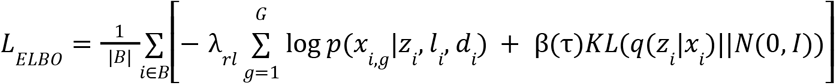

where *p* is the zero-inflated negative binomial (ZINB) of the decoder, conditioned on the cell’s dataset embedding, and *l*_i_ is the observed library size. The reconstruction weight *λ*_rl_ = 0.5 acts inside the ELBO, balancing reconstruction against the KL term; the ELBO as a whole enters *L* with weight 1 and is not rescaled relative to the other terms. The KL weight *β*(*τ*) = 5 × 10⁻⁴ *τ* rises linearly with the zero-based epoch *τ*, following a 200-epoch warm-up toward 0.1 that training stops before completing. It reaches 0.0495 at the final epoch of the production model (*τ* = 99) and 0.0395 at the final epoch of the 12 fold models used for benchmarking, which were trained for 80 epochs (*τ* = 79).

#### Training objective

Five terms are combined with fixed weights,

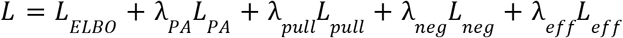

with *λ*_PA_ = 200, *λ*_pull_ = 3 (with its ramp *γ*; see Prior pull), *λ*_neg_ = 1 and *λ*_eff_ = 0.2. Each term reaches a different set of parameters, and the ELBO’s two parts do as well. Its reconstruction part trains the encoder and decoder, and its KL part the encoder. Proxy Anchor trains the encoder and the prototypes; the prior pull trains the 1,004 gated prototypes only; the prior negatives train the encoder only; and the effect-size loss trains the effect-size head only. See **Table S4** for a full list of hyperparameters.

#### Training procedure

The production model (v0.1.0) was trained for 100 epochs of 1,500 batches of 2,048 cells, 150,000 steps in all. Batches were drawn with replacement from the training cells by a weighted random sampler, each cell weighted by (*n* + 1)^−1/2^, where *n* is the number of training cells sharing its perturbation and dataset, with weights normalized to a mean of 1. A perturbation–dataset group is therefore sampled roughly in proportion to √*n*. Batches mix datasets and perturbations, with no structured or contrastive composition. Optimization used AdamW at a constant learning rate of 5 × 10⁻³, weight decay 10⁻⁵ and *ε* = 0.01, with gradients clipped to a global norm of 1.0, in fp32. The prototype embedding and the cross-fit permutation were seeded (seed 0); no global seed was set. Internal validation used a fixed subset of at most 5 cells per perturbation and dataset, drawn once with a fixed seed and held constant across epochs, from which up to 50,000 cells were cached for the per-epoch monitor, macro-averaged top-10 recall excluding datasets using a “Marson2025”-prefix. The validation subsets served only to monitor training; it played no part in checkpoint or hyperparameter selection or in the benchmarks of Figure 2, and the same split was used for every benchmarked model. Early stopping was configured but never fired within the production run, and the final state at epoch 99 was taken rather than the best-monitored checkpoint. The zero-shot fill and the effect-size readout were installed after training (see Unseen perturbations; Perturbation effect size). The 12 fold models followed the same procedure over 80 epochs instead of 100.

#### Inference

A query cell is centered on the control mean of its own dataset at the input, encoded with the posterior mean, and centered in the latent as during training, giving *u*. It is scored against every prototype in one pass by *s_p_* = cos(*u*, *a_p_*). Ranking runs over whichever subset of the table the question admits: the 8,964 trained prototypes for a screened candidate set, or all 19,895 rows when perturbations absent from training are candidates. Scores are signed, so the ranking can be read from either end. The highest cosines name perturbations whose knockdown the cell resembles (loss tail), and the most negative name perturbations whose knockdown it opposes (gain tail). No hubness correction was applied.

Input control means are refreshed from the query’s own control cells where a dataset contributes at least 10, and otherwise the trained baseline is kept. A dataset absent from training has no stored latent mean, so its latent is centered on the mean of its control cells in the same call, or on the mean of all its cells if none are labeled as controls. Scores for such a dataset therefore depend on which of its cells are scored together, and a dataset must not be split across calls. In benchmarking, control means were refreshed once from the pooled control cells and prediction run per whole dataset.

### Datasets and analyses

#### Benchmarking against perturbation-prediction methods

Most published perturbation models are formulated in the opposite direction, mapping a perturbation to an expected expression response. Only RNA fingerprinting^38^ and the non-learned reference floors are natively inverse; CellOT^32^, GEARS^31^, scGPT^34^, scouter^35^ and GenePert^53^ are forward models and must be inverted to be compared. Each inversion is stated explicitly below, and in each case we chose the form most favorable to that method that its published formulation supports. Every method emits the same artifact, a matrix of per-cell scores over the candidate perturbations together with the row positions of the scored cells in the source data, and a single collector scores all matrices with identical code; ground-truth labels are read from the source annotation, never from the method’s output.

#### Benchmark data and folds

All methods were run on subsets of the database described above, over the same 22 datasets and 3,096-gene union panel. The primary benchmark comprises the 1,157 perturbations assayed in at least seven of the ten cellular contexts (2,741,678 cells); a 200-perturbation subset (671,242 cells) is used only to measure how cost scales with the number of candidates. Each of 12 folds holds out one to four datasets: folds 0–3 hold out one T cell donor (all three conditions) together with one cell line, folds 4–8 a cell line, and folds 9–11 a stimulation condition across all donors. Per fold, the 1,157-perturbation subset has on average 2.17 million training, 131,639 in-distribution (ID) test and 17,523 out-of-distribution (OOD) test cells; OOD test cells are capped at 10 per dataset and perturbation.

#### ExPert scores

ExPert models were trained outside the benchmark using the ExPert python package (https://github.com/GoyalLab/ExPert). We trained one model per fold of the split set over all 8,964 perturbations. Benchmark and model folds were paired by held-out dataset set (an exact match, otherwise the smallest model fold whose held-out set contains the benchmark fold’s); all 12 folds of the primary benchmark are covered, and 11 of 12 of the 200-perturbation subset. Benchmark cells present in the paired model’s training set were excluded by cell name. The score is the cosine similarity between a cell’s centered latent and each perturbation’s learned prototype, restricted to the benchmark’s candidates. Because a dataset absent from training is centered on its mean over the cells in the same prediction call, predictions were made on whole datasets.

#### Per-dataset training and dataset-blind evaluation

CellOT^32^, GEARS^31^, scouter^35^, GenePert^53^ and RNA fingerprinting^38^ are single-dataset methods, so we trained one model per dataset on that dataset’s cells over the union panel and evaluated dataset-blind: because a query cell does not reveal which model applies, each test cell is scored by every dataset’s model. Per-cell scores are converted to a softmax over each model’s perturbations, averaged over the models whose class space contains the perturbation, and log-transformed; each perturbation column is then z-scored across cells, which removes per-perturbation baseline offsets without using labels or dataset identity.

#### CellOT

A per-dataset scGen^28^ autoencoder with per-perturbation optimal-transport maps. Cells are scored by −ǁδ_c_ − Δ_p_ǁ, where Δ_p_ is the predicted latent shift and δ_c_ the cell’s latent shift from its own dataset’s controls.

#### GEARS

A per-dataset graph network. Cells are scored by the Pearson correlation between the cell’s deviation from its own dataset’s control mean and each predicted response.

#### scouter

A per-dataset model, scored on its predicted mean response as for GEARS.

#### GenePert

Per-dataset ridge regression (*α* = 0.1) from the target gene’s embedding (Qwen3-Embedding-8B^58,59^; see below) to mean post-perturbation expression, scored as for GEARS.

#### RNA fingerprinting (ensemble)

We followed the package instructions for generating RNA fingerprints^38^ wherever possible. Per-dataset SCTransform^125^, differential-expression selection and fingerprint dictionary, queried per cell using the provided package (GitHub commit: 4004c8d) against each dataset’s dictionary individually and combined as above.

#### RNA fingerprinting (merged)

The per-dataset dictionaries merged into one, keeping for each perturbation the fingerprint of its most confident dataset after per-dataset scale normalization, with one query per cell.

### Single-model comparisons

scGPT^34^, XGBoost^54^ and linear regression train one model per fold across all datasets. scGPT was fine-tuned from scGPT_human (15 epochs of 2,500 steps, batch size 64, learning rate 10⁻⁴, mixed precision). For each test dataset and candidate, it predicts a profile from 32 control cells with the candidate’s target-gene token flagged, and test cells are scored by cosine similarity to these profiles over the 3,096 panel genes (ID test cells uniformly subsampled to 25,000 per fold). Because scGPT can express only perturbations whose target gene has a token, it was run on a copy of the data carrying one additional column per target gene outside the panel (measured expression where available, Poisson draws otherwise), making 712 of 1,157 perturbations expressible; the added columns are excluded from scoring. XGBoost 38 (multi:softprob; 150 rounds, depth 4, learning rate 0.2, column subsample 0.3, row subsample 0.5, early stopping after 20 rounds on the validation split) and linear regression (one-vs-rest logistic regression by stochastic gradient descent, α = 10⁻⁵, 30 iterations) take CPM-normalized, log1p-transformed expression minus the cell’s own dataset control mean; XGBoost training was capped at 500,000 cells for memory.

### Baselines

Random emits uniform scores. Prior scores every cell with the log training frequency of each perturbation (additive smoothing). Centroid scores the cosine similarity between a cell’s control-relative log-normalized profile and each perturbation’s mean training profile, with that perturbation’s target gene masked. Negative target scores each perturbation by how far its target gene falls below the dataset’s control mean in the query cell; for the 882 of 1,157 perturbations whose target gene is not among the input genes, the same score is computed on a decoy gene (an input gene that is not the target of any candidate, drawn at random with a fixed seed and distinct for each perturbation), so that every perturbation competes on equal terms. Negative target is shown only in the target-masking analysis (**Figure S3C**).

### Metrics

Score matrices are converted to log-probabilities, and perturbations a method cannot express receive a score of negative infinity. The rank _r_ of the true perturbation among the 1,157 candidates counts ties as half; the top-ranked call breaks ties by one fixed random order of the candidates. Metrics are computed separately for each held-out dataset in each fold (a dataset by fold combination) and averaged over the combinations available for every compared method. Within a combination, each perturbation present contributes one vote: top-k recall is the fraction of its queries whose true perturbation ranks among the top-k, mean reciprocal rank (MRR) is the mean of 1/(*r* + 1), and F1 combines its recall with its precision among all queries called as that perturbation.

Queries are scored at three levels. For pseudobulks (**Figures 2A–2C**), the log-probabilities of all cells of one perturbation in one dataset are averaged and the pooled vector is ranked once. For single cells, each cell is ranked separately and cells are weighted by their knockdown effect size (a Mixscale^120^–style perturbation score; cells with non-positive or missing scores excluded) when metrics are averaged within a perturbation (**Figure 2E**). For the effect-size analysis (**Figure 2D**), single cells are scored unweighted after keeping those with effect size of at least *t* ∈ {0, 1, 2, 3}, where t = 0 keeps all cells; here MRR is averaged over cells. As a control (**Figure S3E**), a random sample of the same number of cells, drawn regardless of effect size, is scored at each threshold (mean of five draws). The per-cell association between effect size and retrieval is the Spearman correlation, within each combination with at least 20 cells, between each cell’s effect size and its outcome (top-1 hit, top-10 hit or reciprocal rank).

Fold change over random divides each value by its expectation under random ranking of C = 1,157 candidates: 1/*C* for F1, *k/C* for top-k recall and *H_C_/C* for MRR, where *H_C_* is the *C*-th harmonic number. The increase over the next-best method is, for each fold, ExPert’s mean over that fold’s combinations divided by the highest such mean among all other methods, summarized as the median across the 12 folds; ExPert and the per-fold next-best method were compared with a two-sided Wilcoxon signed-rank test over folds.

### Compute cost and scaling

Costs were read from the scheduler’s accounting database (SLURM^126^) and attributed to method, benchmark size and stage by workflow rule, counting only completed jobs, each by its last completed attempt. Compute is CPU core-hours plus GPU-hours at a one-to-one exchange, the cluster’s billing weight. **Figures 2E–2I** report evaluation only; XGBoost, linear regression and GenePert train and evaluate in one job, so their costs include training and are marked with asterisks. ExPert’s evaluation cost was measured on fold 0 (eight CPU cores) and multiplied by the number of folds, because the scheduled runs also loaded the training data, which inference does not require.

Scaling compares the 200-and 1,157-perturbation subsets per fold, excluding scGPT, GEARS and CellOT, whose readout, hardware or code changed between the two sizes. For each quantity, the exponent *b* in cost ∝ *C^b^* is the log-log slope between the two sizes (0, flat; 1, linear). The summary quantity is the evaluation burden, compute-hours multiplied by jobs per fold, so that a method keeping per-job cost flat by spreading work over more jobs still scales poorly.

### Implementation

The benchmark is one Snakemake workflow over benchmark size, fold, method and split, with a single target that produces every method’s score matrix and the scoring rollup; each rule declares its own pinned conda environment, which lets incompatible dependencies coexist (for example, CellOT on torch 1.11.0 and scGPT on torch 2.5.1), and ExPert runs in its own environment. A complete 12-fold pass comprises 24,029 scheduler jobs for the 1,157-perturbation subset and 6,386 for the 200-perturbation subset. One versioned configuration file pins the data build, methods, hyperparameters and per-rule resources; the source data path is content-addressed, and split generation and the tie-break order are seeded. Software versions are listed in the publicly available GitHub repository GoyalLab/ExPert. All computation was performed on a Slurm-managed HPC cluster.

### Zero-shot placement of untrained perturbations

The model vocabulary covers 19,895 genes. The 8,964 trained perturbations have reliable training data (**Methods**); the remaining 10,931 genes have no training cells. Each untrained gene u receives a prototype from the prior graph. In each prior bank (gene descriptions embedded with qwen3-embed-8b^58,59^, DepMap^46,47^ and STRING^51,52^; see Gene priors), the bank is centered and scaled to unit mean row norm, and the 25 trained genes most similar to u by cosine are taken. The bank’s placement is the weighted mean of those neighbors’ unit-normalized learned prototypes, with weights proportional to the fourth power of the prior similarity and negative similarities set to zero. Placements are averaged over the banks that cover u, normalized, and scaled to the median prototype norm. The fourth power applies to untrained genes only; the prior pull on trained prototypes during training uses linear similarity weights (**Methods**, prior pull). Placements were computed once on each frozen model and stored with it; no model was retrained. The exponent and neighborhood size were selected on the 12 fold models by the summed change in zero-shot recovery, in and out of distribution, and in recovery of trained perturbations, relative to linear weights at k = 25 (**Figure S6**; **Table S6**).

### Zero-shot retrieval (Figure 3B)

Of the 10,931 untrained genes, 9,874 were perturbed in at least one screen of the corpus. Their cells were held in a separate database that no model saw in training. A query is the pseudobulk mean latent of between 5 and 25 cells of one perturbation in one screen. Queries were encoded by each of the 12 fold models (see **Table S5**) and ranked by cosine similarity against all 19,895 prototypes, trained and zero-shot. For each model and group of screens, the queries of one perturbation were averaged into a single query. Top-10 recall was weighted by the within-screen percentile of the perturbation’s mean knockdown effect size and expressed as fold over chance (10/19,895). A screen is in distribution (ID) for a model if its context was in that model’s training data and out of distribution (OOD) if the model held its context out; each context is held out by exactly one model. As a null, the zero-shot prototypes were permuted across untrained genes, 20 times per model. The summary row “all trained” ranks the held-out cells of the 8,964 trained perturbations, under the same cap and score, against the same 19,895 candidates. Values are medians over the 12 models (**Table S6**).

### Placement of zero-shot prototypes (Figure 3C)

On the production model, the 8,964 trained prototypes were clustered by Ward linkage on unit vectors and the dendrogram drawn in leaf order around a circle. Each of the 10,931 zero-shot prototypes was placed beside its most similar trained prototype by cosine, ordered by decreasing similarity within each trained leaf. Clade statistics use a cut of the trained tree into 60 clades. Nearest-neighbor shares count, for each prototype, the zero-shot prototypes among its 10 nearest prototypes by cosine, against the share expected if kind did not affect proximity (10,931/19,894).

### Prior pull during training (Figure 3D)

Three models were trained with both K562 screens held out (fold 4) under identical settings except the prior pull: no pull; pull restricted to the 1,004 prototypes with the lowest training support, as in the production model (see Augmented learned prototypes); and the same pull applied with equal weight to every prototype. Each was scored by pooled top-10 recall of 10,061 held-out K562 pseudobulk queries of trained perturbations, up to 25 cells each, against the 8,964 trained prototypes. Run-to-run variation was estimated from two independently trained replicates of one configuration by a paired bootstrap over queries, and each configuration was compared with the no-pull model by a two-sided z test against that distribution. For the support analysis (**Figure S6**), queries were binned by decile of their perturbation’s training support, training cells multiplied by training datasets, and per-query changes in top-10 hits relative to the no-pull model were summed per bin, with 95% confidence intervals from 2,000 bootstrap resamples of queries. The post-hoc comparison moved the same 1,004 prototypes of the trained no-pull model toward their graph targets by a static blend, unit((1 − β)a + βT), for β from 0.05 to 1.0, counting top-10 changes on the moved prototypes’ own queries and on all other queries.

### Gene priors

Each prior is a matrix with one row per gene; genes absent from a source are not covered by it. Gene symbols were matched exactly to the model vocabulary.

#### Cancer dependency maps (DepMap)

CRISPR gene-effect scores^46,47^ (DepMap 24Q4, CRISPRGeneEffect) across 1,178 cell lines. Genes with more than 20% missing values were removed and remaining missing values set to the gene’s mean (17,787 genes). Two versions removing the dominant essentiality axis were also built: double standardization, in which cell-line and gene means were removed and rows and columns then alternately scaled to zero mean and unit variance for 20 iterations (DepMap, used by the production model), and removal of the top two singular components after centering cell lines (DepMap, drop 2).

#### Protein-protein interactions

Protein-protein interactions (PPI) were downloaded from STRING^51,52^ v12.0 (https://string-db.org, human, taxon 9606, file 9606.protein.links.detailed.v12.0.txt.gz, accessed July 8, 2026). STRING protein identifiers were mapped to gene symbols with the matching protein info file (9606.protein.info.v12.0.txt.gz). Interactions were kept when both genes were in the model vocabulary (19,895 genes) and treated as undirected, one edge per gene pair. This left 6,531,794 scored edges over 18,550 genes. Three PPI priors were built from these edges, each using a different STRING score:

**STRING**: combined score ≥ 400; 888,859 edges over 18,511 genes. (production)

**STRING experimental**: experimental channel score ≥ 150; 799,924 edges over 17,805 genes.

**STRING co-expression**: co-expression channel score ≥ 150; 862,941 edges over 17,305 genes.

For each prior, the retained edges formed a symmetric weighted adjacency matrix *A* (weight = score / 1,000). *A* was symmetrically normalized as *D^-1/2^ A D^-1/2^*, where *D* is the diagonal degree matrix, so that highly connected genes do not dominate. The normalized matrix was then embedded into 256 dimensions by randomized singular value decomposition (5 power iterations, fixed seed), using *U·S* as the gene embedding. Genes with similar interaction neighborhoods therefore get similar embeddings. A gene with no retained edge has no embedding in that prior. Each prior was paired with a shuffled control, in which gene labels were permuted across embedding rows.

#### Morphology

JUMP^114^ Cell Painting (cpg0016) assembled, batch-corrected profiles of CRISPR knockouts and ORF overexpression, with each gene’s vector the median over its wells (CRISPR, 7,977 genes over 599 features; ORF, 12,602 genes over 722 features).

#### Proteome

A protein intensity matrix downloaded from the ProteinTalks^127^ data portal (db.prottalks.com, accessed September 13, 2026): data-independent acquisition mass spectrometry of 2,856 human tissue samples (1,105 tumor, 1,088 adjacent non-tumor, 549 normal and 114 fetal, from 80 major tissue types), quantifying 13,609 proteins. A gene’s prior is the abundance profile of its protein across samples, so the prior is measured independently of whether the gene was perturbed. Intensities were log2-transformed (zero or missing values treated as missing) and centered per sample on the median. Proteins observed in fewer than 30% of samples were removed, and protein groups mapping to the same gene were averaged, leaving 7,746 genes. Each gene was z-scored across all samples (Proteome, global) or within each major tissue type (Proteome, within tissue; tissue types with fewer than 10 samples excluded). Missing values were then set to zero, and the matrix was embedded by randomized SVD into 256 dimensions.

#### L1000^39,40^

Consensus shRNA knockdown signatures over the 978 landmark genes (GEO: GSE106127), z-scored per landmark within each cell line and averaged per target gene (4,648 genes).

#### Genomic position

GENCODE^118^ release 46 gene records encoded as a chromosome one-hot, sine and cosine encodings of the gene start at periods of 0.1–100 Mb, and gene length, strand and gene type (53 dimensions). Blocks were scaled to unit mean norm and the metadata block down-weighted by 0.2.

#### Shuffled controls

For every prior, a twin was built with gene labels permuted across rows.

### Gene descriptions and text embeddings (Figure 3E)

For each gene, a language model wrote a functional description of at most 256 words from records retrieved for that gene: the NCBI^128^ Gene summary; UniProt^129^ function, subcellular location, pathway, interaction, subunit, disease, catalytic activity, keyword and domain annotations; the top ten STRING^52^ interaction partners with confidence; DepMap^46,47^ dependency and disease associations from Open Targets^130^; and PubMed literature. Three writers were used, Claude Sonnet 4.5, Claude Opus 4.7 and Claude Opus 4.8, with the instructions reproduced in **Supplementary Note 4**. Each description set was embedded by 13 text encoders. BioLORD-2023-M^131^, SapBERT^132^ (from PubMedBERT, full text) and BiomedBERT^133^ (abstract and full text) were applied by mean pooling of the last hidden states over at most 512 tokens with L2 normalization. OpenAI text-embedding-3-large^134^, at 3,072 and 1,024 dimensions, and text-embedding-ada-002^134^ were queried through the API. Qwen3-Embedding^58,59^ (0.6B, 4B and 8B), Llama-Embed-Nemotron-8B^135^, Llama-Nemotron-Embed-1B^136^, BGE-M3^137,138^ and MedCPT-Article^139^ completed the set. Embeddings were obtained by mean pooling for BioLORD, SapBERT and PubMedBERT, from the “CLS” token for BGE-M3 and MedCPT, and from the final token for the Qwen3-Embedding and Nemotron models. The published GenePT^50^ embeddings (version 2: ada-002 embeddings of NCBI summaries, and version 3: text-embedding-3-large embeddings of NCBI and UniProt summaries; Zenodo: 10833191) were included unchanged. The production model uses Opus 4.8 descriptions embedded with Qwen3-Embedding-8B (4,096 dimensions).

### Prior quality and zero-shot screen (Figure 3F)

Each of 48 priors was scored in two ways. Biological content, independent of ExPert, was the lift in KEGG^57,110^ pathway co-membership among each gene’s 10 nearest neighbors. The bank was centered and restricted to genes annotated in KEGG 2021 (Human), and for each annotated gene the share of its 10 cosine neighbors sharing at least one pathway was divided by the share expected over all gene pairs. To remove the effect of coverage, 2,000 annotated genes were subsampled and the lift averaged over five draws. Zero-shot recovery used one fold-4 model trained without any prior information. Each prior alone placed the 9,874 measured untrained genes under the production placement, including the fourth-power weights, and was scored as in **Figure 3B** against all 19,895 prototypes, pooling all screens. As a null, each prior’s gene-to-vector assignment was permuted three times. Associations were tested by two-sided Spearman correlation.

### Prior combinations (Figure 3G)

The ten best single priors by zero-shot recovery, with at most three text priors, formed the pool. Every nonempty subset (1,023) was fused as in the production model, with unit-normalized placements averaged at equal weight, and scored as above; each subset’s null fused the members’ permuted placements. The candidate genes were split in half at random, the best subset of each size was selected on one half and scored on the other, and the two directions were averaged. The production combination falls outside the pool and was scored separately (**Table S7**).

### Recovery of combinatorial perturbations (Figures 3H–3J)

Double-perturbation pseudobulks from four screens not used in training were encoded by the production model: CRISPRi (Replogle et al.^60^), Cas13d (CaRPool-seq^61^) and CRISPRa (Norman et al., 2019^62^; ProPer-seq^63^). Each double was scored as the cosine of its control-centered latent shift to the unit prototypes, ranked in descending order for loss-of-function screens and ascending order for CRISPRa. RNA fingerprinting^38^ was fitted to genome-wide K562 Perturb-seq^19^ with the RNAFingerprinting package, using only the training cells of the production model, so that both methods saw the same K562 data. The 8,347 perturbations of that screen in ExPert’s training set were used, with at most 100 cells per perturbation drawn at random (794,828 of 1,491,224 training cells; median 163 per perturbation) and up to 2,000 control cells. Expression was restricted to the screen’s measured genes within ExPert’s 3,096-gene input panel. It was normalized by SCTransform, with one negative-binomial model fitted on a subsample and applied to all cells, so every fingerprint lies in the same residual space. Features were the union, over perturbations, of genes differentially expressed against controls (Seurat FindMarkers, *P* < 0.05, at most 500 per perturbation; 1,706 genes). The dictionary was learned with LearnDictionary on these features, giving 8,347 fingerprints, all of them trained ExPert perturbations. Against the published genome-wide dictionary (4,098 fingerprints over 8,141 features), same-gene fingerprints agreed with a median correlation of 0.67 (shuffled-gene null, 0.004) on shared genes and features. Candidates were ranked by posterior inclusion probability, with ties broken by correlation. The centroid baseline ranked candidates by the Pearson correlation between the double’s control-relative profile and each perturbation’s mean training profile, with the targeted genes excluded; the random baseline is computed analytically. All methods ranked the same 8,347 candidates, on doubles whose two genes were both candidates and both measured as single perturbations in that screen (134 loss-of-function, 133 gain-of-function). A double counts as recovered at k if either gene (**Figures 3H** and **3I**) or both genes (**Figure S5B,C**) rank within the top k.

### Drug screens, target annotation and response scoring (Figure 4)

We queried three drug screens with ExPert (v0.1.0); labels, prototypes and readout were unchanged from training, and no part of the model was retrained, fine-tuned or recalibrated. sci-Plex3^21^ covers 188 drugs at four doses (0.01–10 µM) in A549, K562 and MCF7 cells after 24 h; we sampled at most 150 cells per drug by dose by cell line and kept every vehicle cell as controls. The FlexPlex screen^38^ covers four HDAC inhibitors (AR-42, tacedinaline, ricolinostat and ACY-738) at eight doses (0.1–10 µM) in K562 cells and we used all 15,626 cells for the analysis. Tahoe-100M^64^ covers 379 small molecules at three doses (0.05, 0.5 and 5 µM) across 50 cancer cell lines.

Scoring drug target recovery requires gene-level targets for each drug, something the dataset specific annotations may not always provide. For instance sci-Plex3 annotates each compound only with a protein family which includes many genes, some of which may not be targeted by the drug. We therefore matched each drug to ChEMBL^140^ by name or synonym, checking non-exact matches by molecular formula, and took its targets from ChEMBL’s curated mechanism of action records, narrowing protein family targets to members inhibited within one log unit of the drug’s most potent target in that family. For drugs without a mechanism record, we used single protein targets with pChEMBL ≥ 7 within one log unit of the most potent target. For 30 drugs without a usable ChEMBL target we took targets from the literature cited in drug vendor sheets (Selleck chemical drug vendor sheets accessed October 2026; **Table S8C**). We excluded two drugs whose only ChEMBL targets were not credible: curcumin, a promiscuous compound with potent activities recorded against more than a dozen unrelated protein classes, and cyclocytidine, a cytarabine prodrug whose sole target record comes from a cell-based lamin A splicing screen rather than a binding assay. This left 176 annotated drugs. 23 drugs had no target among the 3,802 perturbations shared by ExPert and RNA fingerprinting, 20 of them because their ChEMBL targets lay outside this set and 3 because they had no ChEMBL target; for these we added the genes of their sci-Plex3 protein family. In total, 179 of the 188 drugs had at least one annotated target, and 111 had a target among the 3,802 perturbations shared by ExPert’s trained prototypes and the RNA fingerprinting dictionary. Tahoe-100M annotates drugs at the gene level, so there we used the dataset’s own annotation.

For sci-Plex3 and Tahoe-100M we pseudobulked cells per plate, cell line, drug, and dose, keeping those with at least 10 cells in sci-Plex3 and 30 in Tahoe-100M, and passed them through the ExPert encoder, which maps each pseudobulk to a 2,048-dimensional embedding in the latent space shared with the prototypes, with mitochondrial genes masked while centring each cell line on its own controls. Each response was the change in latent state (Δz) relative to matched controls from the same cell line (vehicle cells in sci-Plex3, DMSO wells in Tahoe-100M; FlexPlex is described below). We scored Δz against the trained prototypes with the model’s shipped readout: the top 10 positions are the plain cosine ranking, and deeper positions merge two further rounds, each of which projects out the rank-3 subspace spanned by the previous round’s 25 highest-scoring prototypes and re-scores the residual against the whole table. A drug effect need not be a knockdown of its target — it may also activate it — so we ranked prototypes in both the loss-of-function and the gain-of-function direction.

### Comparison with RNA fingerprinting in sci-Plex3 and FlexPlex

For sci-Plex3 we compared ExPert with RNA fingerprinting, a training-centroid baseline and a random baseline, all ranking the same 3,802 candidate perturbations shared by ExPert’s trained prototypes and the RNA fingerprinting dictionary. All methods were scored on the same cells. We ran RNA fingerprinting separately on each of the three cell lines with its published genome-wide K562 dictionary, restricted to ExPert’s input genes and the 3,802 candidates. Each run took every vehicle cell of that line and up to 200 cells per drug by dose, kept genes with a mean of at least 0.01 UMI per cell, and pooled each condition’s replicate plates, giving one ranking per cell line by drug and dose; candidates were ordered by their posterior inclusion probability, with ties broken by the absolute correlation between the drug response and each candidate’s fingerprint. For K562 this reproduces the published fingerprinting run exactly. ExPert, the training centroids and the random baseline read exactly these cells, grouped the same way, so every method has one input per cell line, drug, and dose combination. The training centroids ranked candidates by the Pearson correlation between a condition’s expression change from vehicle and each perturbation’s expression change in the training corpus, restricted to the model’s input genes. The random baseline drew one uniform rank per condition.

We scored only responsive conditions, in which the drug moved the cells’ embedding further than control samples vary among themselves. Control variation has to be measured at the depth of a drug condition, so we split each line’s vehicle cells into pseudobulks of about 200 cells, the size of a drug condition (median 200), measured how far each one’s embedding lay from the mean embedding of the other vehicle pseudobulks of the same line, and took the 95th percentile of these distances. A drug condition was responsive if its distance from its cell line’s vehicle exceeded this threshold. We applied the same responsiveness filter to every method, per cell line, drug, and dose. A drug counted as recovered at rank cutoff k when any of its responsive conditions placed an annotated target within the top k. We compared methods on the 110 drugs with at least one responsive condition that every method could score (**Figure 4B**), and by dose on the drugs responding at each dose (60, 63, 82 and 96 drugs at 0.01, 0.1, 1 and 10 µM; **Figure 4C**).

We also scored the 20 HDAC inhibitors against their complexes rather than a single annotated subunit, because HDAC1–3 act within multi-subunit corepressor complexes. Each condition was scored against the 48-gene HDAC machinery, comprising the SIN3, CoREST, NuRD and NCoR complexes, the HDAC1–3 catalytic subunits and CORUM^112^–core partners of the HDAC family (**Table S8A**). The 23 HDAC inhibitors with a responsive condition give 92 drug by dose pairs, scored for every method on the same 69 conditions per dose across the three cell lines (**Figure 4D**). CORUM-core partners are genes that share CORUM complexes of at most 20 members with a drug target gene. We scored each gene *g* as *S_g_*= *n_g_/√(N_T_·D_g_)*, where *n_g_*is the number of these complexes containing *g*, *N_T_* is the total number of these complexes, and *D_g_* is the number of all CORUM complexes containing *g*, and kept genes with *S_g_* ≥ 0.15. We set the size limit and threshold on an independent set of 35 subunits of the four class I HDAC corepressor complexes ^141,142^ (**Table S8B**). Complex portals were last accessed on October, 26th, 2026. For each concentration, we measured their similarity to AR-42 as the Spearman correlation between their rankings over the same 3,802 candidates, computed identically for ExPert and RNA fingerprinting (**Figure 4E**).

In the FlexPlex screen^38^, we measured the similarity of ricolinostat and ACY-738 to the pan-HDAC inhibitor AR-42 as the Spearman correlation between their rankings at the same concentration over the same 3,802 candidates, computed in the same way for ExPert and RNA fingerprinting (**Figure 4E**). Here we encoded each cell rather than pseudobulks, took each drug by dose response as the mean embedding of its cells minus that of the pooled untreated (NT) cells, kept conditions with at least 30 cells, and ranked the candidates by plain cosine against the trained prototypes. ExPert’s input genes were restricted to the 1,408 genes RNA fingerprinting uses, so neither method saw genes the other could not; RNA fingerprinting’s rankings were its own per-condition output, ordered as above.

### Target and pathway recovery in Tahoe-100M

As in sci-Plex3, we scored only responsive conditions, those whose ǁΔzǁ exceeded the 95th percentile of DMSO pseudobulks (25% of conditions), and evaluated the 140 drugs that had an annotated target among the trained prototypes, at least three responsive conditions, and a known direction of action (130 inhibitors, 10 activators). We defined a target’s partners as the other genes in any CORUM complex or Reactome 2024 pathway that contains the target and has 3 to 30 members. Because a target can belong to many sets, we capped each drug at 25 partners, taking genes from the smallest, most specific sets first. We excluded the targets themselves, so that partner recovery tests something separate from target recovery. Five drugs had no partners and were left out of the partner comparison only. For each drug, we computed the fraction of its responsive conditions that placed an annotated target, or a partner, in the top 10 (**Figure 4F**). As a null, we repeated this for 20 random gene sets of the same size, replacing each gene with a random prototype of similar popularity, defined by its mean cosine across all responsive drugs (20 equal-sized bins), so that genes that score highly for every drug could not inflate the comparison.

For the worked examples (**Figure 4G, H**), we scored each drug by dose against the trained prototypes with corrections for Tahoe-100M’s plate layout. All 50 cell lines share a well, and each block of about 90 drugs is spread over three plates (one per dose), with a drug at the same well position on all three. The well position is its order within the plate in the public sample table. A drug’s response in a cell line was its latent state minus that of the DMSO pseudobulk from the same plate by cell line, minus the mean response of all drugs on that plate by cell line. The drug by dose score of a prototype was the mean cosine over cell lines, converted to a z score across the drugs on the same plate, minus the mean z score of the drugs at the same well position in the other blocks. **Figure 4G** shows the rank of each target among the 8,964 trained prototypes at every dose, averaged over the gene pair. For **Figure 4H**, a drug by dose was responsive when its expression response reproduced across the drug’s other doses better than the responses of other drugs did. Reproduction was the maximum cosine to its other doses, over the 3,000 genes whose responses varied most. It was compared with 2,000 draws of other drugs on other plates, and a dose counted as responsive at Benjamini-Hochberg *q* < 0.05. A drug’s score profile was the minimum over its responsive doses of the position-corrected z score.

We measured each drug’s glucocorticoid response in two ways, from ExPert’s prototype scores and directly from gene expression (**Figure 4H**). From ExPert’s scores, we took the mean Pearson correlation between a drug’s score profile across the 8,964 trained prototypes and those of four reference glucocorticoids (dexamethasone, triamcinolone, betamethasone dipropionate and budesonide), computed for drugs with at least two responsive doses. Independently of ExPert, we measured the same response in gene expression, as the increase in expression of eight glucocorticoid target genes (*FKBP5, TSC22D3, PER1, DUSP1, KLF9, CEBPD, DDIT4* and *ERRFI1*) at 5 µM. For each gene, we computed the log fold change against DMSO from the same plate by cell line, converted it to a z score across the 379 drugs, and averaged the eight z scores.

### Recovery with zero-shot prototypes in the candidate bank (Figure S8)

To test whether prototypes for perturbations never seen in training add or dilute signal, we repeated the sci-Plex3 and Tahoe-100M analyses with the 10,931 zero-shot prototypes appended to the 8,964 trained ones (19,895 candidates). Zero-shot prototypes are placed from the prior graph as the mean of their 25 nearest trained neighbours weighted by cosine⁴. Because the candidate pools differ in size and in which conditions they can score, we expressed recovery as fold over random. For sci-Plex3 (**Figure S8A**), random was computed exactly per condition as the probability that a uniform ranking places at least one of the drug’s m annotated targets present in a pool of N candidates within the top-k, 1 − C(N−m, k)/C(N, k), averaged over each pool’s own scorable conditions (2,011, 3,450 and 4,117 conditions from 84, 144 and 172 drugs for the 3,802-candidate, trained and padded pools). For Tahoe-100M (**Figure S8B**), the null was the size-and popularity-matched random gene sets of **Figure 4F**. Intervals are 95% bootstrap intervals over drugs (2,000 resamples).

### Analysis of unrecovered drug targets (Supplementary Note 5)

To probe whether drugs whose annotated target ExPert did not recover were true misses, or instead reflected drug biology absent from their annotation, we used the responsive drugs, readout and complex partner sets of the target-recovery analysis above. A drug’s target was recovered when it ranked in the top ten in more of the drug’s responsive conditions than expected from size-and popularity-matched random gene sets (one-sided binomial test, *p* < 0.05); the remaining 142 of 155 drugs were missed. Misses in which a CORUM-core partner of the target ranked in the top 10 more often than chance were not considered further, as partner recovery is assessed above (**Figure 4F**). Each remaining miss was assigned to the first criterion it met: (i) the target was recovered in the gain-of-function direction; (ii) the drug resembled another drug class, (iii) did not resemble its own class, or (iv) resembled its own class, measured as drug-class similarity, the mean Pearson correlation of the drug’s profile with the annotated members of the class, called at z ≥ 3 against 2,000 random drug sets of the same size from other plates, in both ExPert scores and gene expression; or (v) the same prototypes ranked highly at every active dose (FDR < 0.05 against pseudo-drugs assembled from other drugs’ wells on the same plates), counted only when a top-five prototype was a documented secondary target or a member of the target’s pathway whose loss is expected to mimic the drug. We then checked every flagged miss against the primary literature and removed explanations it did not support. During review, we also accepted two glucocorticoids, dexamethasone and budesonide, whose ExPert scores ranked activation of *FKBP5*, a direct glucocorticoid receptor target, first and third. 13 of the 142 misses showed biology consistent with the drug’s known pharmacology (**Table S9**). Misses that met none of the criteria were not reviewed, so this is a lower bound.

### Prototype neighborhoods between drug targets (Figure S9, Supplementary Note 6)

We built a nearest-neighbor graph over the 8,964 trained prototypes. We joined each prototype to its eight nearest neighbors by cosine similarity, with edge length 1 − cosine. For every pair of the 156 Tahoe-100M drugs with a trained target, we found the shortest route through the graph from any target of one drug to any target of the other; 11,744 of the 12,090 pairs were connected. For each gene we counted the drug pairs whose path passed through it, excluding the drugs’ own targets. Because highly connected prototypes lie on many paths by construction, we compared each count with the count when each drug’s targets were replaced by size-and popularity-matched random genes (200 permutations, 20 popularity bins), and called a gene a bridge when it was traversed by at least 50 drug pairs at least twice as often as expected. 102 genes met this criterion, 89 of them at *p* < 0.05; **Figure S9A** shows the 24 with the largest excess over the null. To ask whether known biology also connects these genes, we built the same graph and ran the same test from each of seven independent gene priors (co-essentiality from DepMap^46,47^, literature-derived gene descriptions, STRING^51,52^ interactions, GO^109^ biological process, L1000 signatures^39,40^, ProteinTalks^127^ and cell morphology^114^); 9 of the 24 were bridges at twice the matched-random rate in at least one prior (DepMap 6, gene descriptions 4, STRING 3, GO 2, L1000 1, morphology 1), and no single prior supported all 9. Genomic position was scored the same way as a negative control.

### Degron tagged lines (Figure 4I)

Degron-tagged iPSC lines were developed as previously described^143^. Briefly, the AAVS1_CAG_OsTIR1(F74G)_PuroR construct was knocked into the AAVS1 safe-harbor locus of KOLF2.2J hiPSCs to give the OCP74G parental line (AID 2.0), and degron tags were then introduced at the C-terminus of each endogenous target by Cas9 RNP nucleofection with a plasmid HDR donor bearing gene-specific homology arms; homozygous clones were identified by PCR genotyping and inducible depletion confirmed by western blot. Each of the 40 lines carries one tagged gene: 29 with miniAID (mAID) and 10 with nanoAID (nAID), the PRMT5 line unrecorded. To create the degron village, 40 degron cell lines were dissociated by ACCUTASE (Stemcell Technologies), pooled together with equal numbers, and 5 × 10⁵ cells were seeded onto Matrigel-coated 6-well plates in Stemflex medium (Invitrogen) supplemented with 10 µM Y27632 (Stemcell Technologies). The next day, the cells were refreshed with fresh Stemflex medium without Y27632. 48 hours later, the cells were treated with DMSO or 1 µM 5-Ph-IAA (MedChemExpress) for 24 hours.

### Western blotting (Figure 4I)

Depletion kinetics were measured in a single experiment, separate from the pooled screen. Two independently derived NAT10-mAID clones (C#4 and C#18) and the OCP74G parental line were treated with 1 µM 5-Ph-IAA for 1, 6 or 24 h, or with DMSO. Cells were washed once with cold DPBS, scraped off the plate, transferred to a microcentrifuge tube and centrifuged at 500g at 4 °C for 5 min. Cells were mixed with 1× RIPA buffer (Boston BioProducts, BP-115) containing 1× Protease Inhibitor Cocktail (Thermo Scientific, 78430) by vortexing every 5 min, and lysed on ice for a total of 30 min. Protein concentrations were determined by BCA assay (Thermo Fisher, 23225). Protein was mixed with 4× Laemmli reducing sample buffer (Boston BioProducts, BP-110R) and boiled at 95 °C for 5 min. Equal amounts of protein were loaded onto 7.5% Tris-glycine gels, and proteins were transferred to a PVDF membrane (Bio-Rad, 1620174) by wet transfer at 4 °C overnight. Membranes were blocked in 5% milk in TBS-T (20 mM Tris, 150 mM NaCl, 0.1% Tween 20, pH 7.6) for 1 h at room temperature, incubated with primary antibody (1:1000) overnight at 4 °C, washed three times for 5 min in TBS-T, incubated with secondary antibody (anti-rabbit IgG (H+L), Promega, W4011; 1:10,000) in blocking buffer for 1 h at room temperature, and washed three times for 10 min in TBS-T. Membranes were developed with SuperSignal West Femto substrate (Thermo Fisher, 34094) and imaged on an iBright system. Primary antibodies were *NAT10* (Cell Signaling, 3041S) and *GAPDH* (Cell Signaling, 2118) as the loading control.

### Auxin-inducible degron screen (Figure 4J)

We pooled 40 clones, each carrying an auxin-inducible degron on one gene, treated them with DMSO or auxin for 24 h, and profiled nuclei in two separate 10x Multiome gene-expression libraries (26,956 DMSO and 31,475 auxin nuclei after quality control, which removed nuclei with more than 20% mitochondrial counts). We assigned nuclei to clones by their 20-bp clone barcode, allowing one mismatch (3,971 assigned nuclei), and scored the 25 clones with at least 10 nuclei in each condition. We pseudobulked each clone and condition, masked mitochondrial genes and passed the pseudobulks through the ExPert encoder with DMSO as the control baseline. Because the two conditions were separate libraries, every clone shares an auxin shift that reflects the library and treatment rather than the depletion; we therefore took each clone’s response as its auxin-minus-DMSO change minus the mean change over all clones. A clone was responsive when this response, measured in expression, exceeded the 95th percentile of split-half differences between its own DMSO nuclei (50 splits, rescaled to the depth of the comparison); 18 of 25 clones passed. We ranked all 19,895 prototypes — the 8,964 trained ones and the 10,931 zero-shot ones, so that the three clone genes never perturbed in training (*MED28*, *NUP62* and *SOX2*) compete on the same list — in both directions with the readout above. For each clone, we compared the rank of its own depleted gene with a null rank: the median rank that the genes depleted in the other clones reached in the same clone’s ranking, which reflects how highly an unrelated gene ranks in that response. We tested whether clones ranked their own gene above this null by a one-sided paired sign-flip permutation test on log₁₀ rank (20,000 permutations) and a paired t-test on log₁₀ rank (**Figure 4J**).

### Human liver samples and datasets

Single-nucleus RNA-seq (snRNA-seq) of human liver was obtained from Chembazhi et al. (GEO GSE281574)^73^. The data comprise liver tissue from 15 individuals: five healthy donors, five patients with alcohol-associated cirrhosis (AC) and five with alcohol-associated hepatitis (AH). For each condition, the nuclei were pooled into two libraries, one of three individuals and one of two. The processed data carry pool labels only, so the pool was the replicate in all snRNA-seq analyses. We analyzed 26,807 nuclei of five cell types: hepatocytes, liver sinusoidal endothelial cells (LSEC), Kupffer cells, hepatic stellate cells (HSC) and T cells; 205 plasma cells were excluded.

Bulk RNA-seq of 16 livers (healthy, n = 4; AC, n = 6; AH, n = 6) from the same study and biobank^73^ was used for differential expression and alternative splicing. Which bulk livers contributed to the snRNA-seq pools is not recorded, so the two datasets are not treated as independent. Liver samples for RNA-FISH, nuclear morphometry and RT-PCR came from the same biobank (patient explants and donor resections, Johns Hopkins Hospital)^73^.

### ExPert readout of the snRNA-seq data

The trained ExPert model (v0.1.0) was applied without retraining. For each pool and cell type, one pseudobulk profile was formed as the mean raw counts of its nuclei; mitochondrial and metallothionein transcripts (MT-, MT1, MT2A, MT3, MT4) were set to zero. All pseudobulks were encoded in one call of predict(), with each cell type’s input centered on its healthy pools (center_by = cell type, control_label = healthy pools) and no hubness correction; the number of nuclei behind each pseudobulk was passed as its depth. All scores below were computed from the returned latents. From each pool’s latent, the unweighted mean latent of its cell type over all six pools was subtracted, removing an offset shared by all conditions (without this step, disease and healthy reads correlated at r = 0.95–1.00). Pools were then combined into condition by cell type groups (nucleus-weighted mean latent). Each group was scored by the cosine similarity of its latent to the 8,964 trained perturbation prototypes: a high score means that the state resembles loss of that gene (predicted loss), a low score that it resembles the opposite (predicted gain). Where a standardized score is shown, scores were z-scored over the prototypes of that condition. ALD denotes AC and AH together; for rankings, the AC and AH hepatocyte top-200 lists were combined.

For the mRNA export machinery (**Figure 5L**), the grouped mRNA export set was divided into six families (nuclear pore complex, TREX and TREX-2, Ran cycle, exon junction complex and cap-binding proteins, SR proteins, exportins). For each family, the predicted-loss z of all its members in ALD (mean of AC and AH) was compared with healthy by a two-sided paired Wilcoxon signed-rank test over the members, unadjusted; in **Figure S12H**, AC and AH were tested separately and Benjamini-Hochberg adjusted over families and disease groups.

This configuration was chosen from a systematic comparison of input centering, latent centering, grouping, readout and prototype bank, scored on known ALD biology, on signal shared with healthy livers (a negative control) and on agreement between pools. For **Figure 5B**, the latents of all nuclei (input centered per cell type on healthy nuclei, latent centered as above, unit-normalized) were embedded with UMAP (umap-learn; 30 neighbors, minimum distance 0.3, cosine metric), and the 25 highest-scoring prototypes of AC and of AH (scores averaged over the five cell types) were placed in the embedding with the fitted transform.

### Pseudobulk differential expression of the snRNA-seq data

Raw counts were summed per pool and cell type (pools with ≥ 30 nuclei). Genes with ≥ 10 counts in ≥ 2 pseudobulks were tested with PyDESeq2^144,145^ (design ∼condition, per cell type over the six pools; Wald test of AC or AH versus healthy, Benjamini-Hochberg false discovery rate, apeGLM-shrunken log2 fold changes). Differentially expressed genes (DEGs) were defined as adjusted *p* < 0.05 and |log2 fold change| ≥ 2. Genes were ranked by the absolute Wald statistic (log2 fold change divided by its standard error); this “DE rank” was used for **Figure 5C** and for the top-100 DEGs of **Figure 5D**. The replicate was the pool. Agreement between ExPert’s ranking and the DE rank (**Figure 5C**) was measured by Spearman’s ρ.

### Pathway enrichment

Gene sets were Reactome 2024 pathways (Enrichr library) with 10–500 members among the scored genes and two curated sets: mRNA export (nuclear pore complex; TREX, TREX-2 and export adaptors; export receptors, exportins and Ran cycle; Reactome transport of mature transcripts) and mRNA splicing (Reactome major and minor spliceosome pathways and all SR proteins). For ExPert (**Figure 5E**), the 200 highest-scoring prototypes in each direction (predicted loss or gain) were tested by one-sided hypergeometric test against all 8,964 trained prototypes; for DE (**Figure 5F**), the 200 most downregulated and upregulated genes were tested against all genes tested for DE. Fold enrichment was computed as (hits + 0.5)/(expected + 0.5) and shown as log2. Viral and disease-named pathways were excluded, and a pathway whose hit genes overlapped those of a higher-ranked pathway by ≥ 50% of the smaller hit set was folded into it. **Figure 5E** shows mRNA export, mRNA splicing and the 13 most significant remaining pathways; **Figure 5F** the 15 most significant; rows are ordered by fold enrichment in ALD. **Figures 5E, 5F** and **S12D**, E report nominal *p* values; all non-redundant pathways with nominal *p* < 0.05 are listed in **Table S6** together with Benjamini-Hochberg *q* values over all gene sets and both directions.

### Bulk RNA-seq: expression and alternative splicing

Bulk paired-end RNA-seq^73^ was aligned to the human genome (hg38, gencode-v43) with STAR v2.7.10a^146^ and counted with featureCounts^147^ (Subread v2.0.4) using uniquely mapping reads. Gene-level counts were modeled with DESeq2^144^ (design ∼group, healthy as reference). In **Figures 5G** and **5K**, genes were called differentially expressed with the thresholds of the snRNA-seq analysis (adjusted *p* < 0.05 and |log2 fold change| ≥ 2). Alternative splicing was analyzed with rMATS-turbo^148,149^ (skipped exon, mutually exclusive exons, alternative 3′ and 5′ splice sites and retained intron); a gene was considered significantly spliced if it had an event with FDR < 0.01 and |ΔPSI| ≥ 0.10. In **Figure 5K**, overlap between ExPert’s top 20 hepatocyte predictions and splicing was assessed at the level of gene families (mRNA export machinery families as in **Figure 5L**, otherwise the gene-symbol root) and compared with 5,000 random 20-gene sets drawn from the genes scored by both (empirical p).

### ExPert scores of the bulk livers (Figure S12A, B)

Each bulk liver was encoded as one pseudobulk (DESeq2-normalized counts, the mitochondrial and metallothionein mask above, rescaled to 5,000 counts, depth 500) in one predict() call with the healthy livers as the input baseline. The mean latent over all livers was subtracted, livers were averaged per condition, and conditions were scored against the trained prototypes and tested for enrichment as above.

### Zero-shot readout (Figure S13)

The condition latents of hepatocytes (centered as above) were scored against all 19,895 prototypes of the model, including the 10,931 zero-shot prototypes placed from prior knowledge (**Figure S6**; cos⁴-weighted mean of the 25 nearest trained prior neighbors). Scores are shown as the change from healthy hepatocytes, in units of the standard deviation of the healthy scores over all prototypes.

### RT-PCR splice assay

Total RNA was extracted from human liver samples with TRIzol. cDNA was synthesized with Maxima Reverse Transcriptase (Thermo Fisher, EP0741), and alternatively spliced regions were amplified with Taq 5X Master Mix (NEB, M0285L) and primers spanning the flanking exons. Products were resolved on 5% polyacrylamide gels and quantified as background-adjusted integrated band volumes. For two-product assays, the percent spliced in was PSI (%) = 100 x V₊/(V₊ + V₋), where V₊ and V₋ are the volumes of the inclusion and exclusion products. For the four-product UPF3A assay, the percent spliced out (PSO) was calculated for the 197-bp product as 100 x V₁₉₇/ΣV. In **Figure 5M**, values are shown as the change from the healthy median, and healthy and ALD (AC and AH pooled) livers were compared by two-sided Welch’s t-test.

### RNA-FISH and microscopy

Formalin-fixed, paraffin-embedded (FFPE) human livers were sectioned at 5 µm, deparaffinized in xylene and rehydrated through graded ethanol (100%, 95%, 80% and 50%) to water. Antigen retrieval was performed in citrate buffer (10 mM sodium citrate, 0.05% Tween-20, pH 6.0) at 85–95 °C for 15–20 min. Poly(A) RNA was detected by HCR Gold RNA-FISH (Molecular Instruments) with Sample Readiness Probes targeting the poly(A) tail (01SRP123) and X3-specific hairpins labeled with a 647-nm fluorophore, according to the manufacturer’s protocol. Nuclei were stained with NucBlue Fixed Cell ReadyProbes (Thermo Fisher, R37606), autofluorescence was quenched with Vector TrueVIEW (SP-8400), and slides were mounted in CC/Mount (Sigma-Aldrich, C9368).

Images were acquired on a Zeiss LSM 900 confocal microscope with the Airyscan 2 module (Institute for Genomic Biology core facility) with 40× or 63× objectives, as z-stacks of 11 optical sections spanning 8 µm around the plane of maximal nuclear signal. Airyscan data were processed by joint deconvolution in ZEN Blue 3.6 and analyzed in Fiji^150^ with a custom macro. Nuclei were segmented with the higher of the Otsu and 60th-percentile nuclear-stain thresholds, after watershed, excluding objects touching the image edge, smaller than 30 µm² or with compactness < 0.5. Poly(A) signal was thresholded at the 90th intensity percentile, and puncta were assigned to the nuclear or cytoplasmic compartment by their centroid. Per imaging frame, the fraction of poly(A) puncta in nuclei, r = N◻/(N◻ + Nₑ), was converted to the nuclear-to-cytoplasmic ratio N/C = r/(1 − r) = N◻/Nₑ (**Figure 5I**). For display (**Figure 5H**), poly(A) intensities were shown in percentile bins. Each imaging frame was treated as an observation, since frames could not be assigned to individual livers, and groups were compared by two-sided Mann-Whitney U test with Benjamini-Hochberg adjustment over the four comparisons shown (**Figures 5I** and **S12F**). Nuclear diameter (**Figure 5J**) was measured on the segmented hepatocyte nuclei as the average diameter in micrometers. Because nuclei of one liver are not independent, groups were compared on per-liver medians by two-sided Mann-Whitney U test (**Figures 5J** and **S12G**).

### Mitochondrial patient collection and storage

Two 4 mL blood samples were collected in green sodium heparin tubes from consented patients in the Neurogenetics Clinic at the Ann & Robert H. Lurie Children’s Hospital of Chicago, in accordance with an approved institutional IRB (NU IRB# 2017-1331). Samples were processed 2–5 hours after collection. Briefly, samples were diluted 1:1 with fresh dPBS + 2% FBS (cPBS), and PBMCs were isolated by layering samples on top of 3.5 mL Ficoll in a SepMate-15 tube (StemCell #85415) and spinning 1200g x 10 min at RT. Cells underwent one round of RBC lysis (3 min RT, BioLegend 420302) and washing with cPBS (300g x 8 min at 4°C) prior to counting with AOPI using a Cellometer K2 and controlled freezing in CryoStor CS-10 (StemCell 07930) at a concentration between 1×10⁶ and 3×10⁶ cells/mL. PBMCs were frozen at −80°C and transferred to vapor-phase liquid nitrogen storage the following day. Typical yields ranged from 5×10⁶ to 1.5×10⁷ total cells.

### Single-cell RNA-seq of PBMCs

Nine patients (four with PMD and five reference donors; see “Donor groups” below) were processed in two batches. Frozen samples were agitated in a 37°C water bath for 1–2 minutes and transferred to 10 mL of pre-warmed HPLM + 10% FBS (cHPLM; Gibco A48991), pelleted (300g x 5 min), resuspended in 1 mL cHPLM, and counted on a Cellometer K2 with AOPI. Additional vials were thawed as needed to obtain ∼2×10⁶ live cells per patient. Cells were resuspended in 190 µL staining buffer (BioLegend 420201) and incubated with 10 µL Human TruStain FcX (BioLegend 422301) for 10 min at 4°C. They were then stained for 30 min at 4°C with anti-human CD3-BV510 (BioLegend 317332), CD20-PE-Cy7 (BioLegend 302312), and CD56-APC (Invitrogen 17-0567-41), all at 1:50. Cells were washed, resuspended in 4 mL Tyto buffer with DAPI (1:1000), and sequentially sorted on a MACSQuant Tyto (Miltenyi) into live T cells (CD3⁺), myeloid cells (CD3⁻CD20⁻CD56⁻), B cells (CD3⁻CD20⁺), and NK cells (CD3⁻CD20⁻CD56⁺).

Sorted populations were recombined for each patient at a 4:4:1:1 ratio (T:myeloid:B:NK; up to 1×10⁵ cells per patient) to enrich for less abundant populations. Each patient pool was hashtagged with a distinct TotalSeq-B hashtag antibody (BioLegend; 50 ng per 1×10⁵ cells in 50 µL) for 25–30 min at 4°C and washed three times in staining buffer. Hashtagged samples were pooled and resuspended in PBS + 0.5% BSA at 1–2×10⁶ cells/mL. Libraries were generated using the 10x Genomics GEM-X Universal 3’ Gene Expression v4 kit on a 10x Genomics Chromium X Controller, according to the manufacturer’s instructions, with each pool loaded across two lanes. Gene expression libraries were first sequenced at shallow depth on an Illumina NextSeq 2000 (P1 flow cell) to estimate cell recovery per library and balance pooling. Gene expression and hashtag libraries were then sequenced together on an Illumina NextSeq 2000 (P4 flow cell), and gene expression reads from both runs were combined for analysis.

Reads were aligned to GRCh38 (10x Genomics reference 2024-A) and quantified with Cell Ranger multi (v9.0.1)^151^ using the nf-core/scrnaseq pipeline (v4.1.0)^152^. Quality control, demultiplexing and annotation were performed in R (v4.4.0) with Seurat (v5.3.0)^153^. Cells were demultiplexed by hashtag using HTODemux^154^, and heterotypic doublets were identified using DoubletFinder (v2.0.6)^155^. Cells were removed if HTODemux called them doublets or negatives, if DoubletFinder called them doublets, or if they had >10% mitochondrial reads, <500 or>15,000 UMIs, or <200 or>4,500 detected genes. Data were normalized with SCTransform (v2)^125,156^, integrated across patients by reciprocal PCA^121^, and clustered on the first 30 integrated dimensions (resolution 0.5). Clusters were annotated using a combination of manual review of commonly expressed cell-type markers alongside automated labeling using SingleR (v2.8.0)^157^ against the Monaco Immune reference (celldex v1.16.0)^158^.

### Donor groups (Table S11)

The four PMD patients carried variants in *NDUFS8* (PT54), *NDUFA10* (PT46), *BCS1L* (PT43) or *PC* (PT67). The five reference donors were two siblings with a non-mitochondrial diagnosis (non-PMD; PT41, PT42) and three undiagnosed patients (PT38, PT50, PT55). PT67 was originally enrolled as a reference donor; after ExPert placed this patient with the PMD patients, the clinical genetics were re-reviewed and compound heterozygous variants in PC, including a known pathogenic variant, were identified, so PT67 was assigned to the PMD group for all analyses shown.

### ExPert readout of the single-cell data

The trained ExPert model (v0.1.0) was applied without retraining. Mitochondrially encoded and metallothionein transcripts (MT-, MT1, MT2A, MT3, MT4) were removed from the input. All 28,513 cells were encoded in one call of predict(), with each cell type’s input centered on the cells of that type from the five reference donors (center_by = cell type, control_label = reference cells), no latent centering and no hubness correction. All scores below were computed from the returned latents and the 8,964 trained perturbation prototypes, by cosine similarity: a high score means that the state resembles loss of that gene (predicted loss of function).

#### Group profile (Figure 6B)

For monocytes, the PMD profile was the mean latent of all PMD monocytes minus the mean latent of all reference monocytes (cells pooled across donors) and was scored against every prototype. It was compared with differential expression of the same cells: the log₂ fold change of PMD versus reference monocytes, computed on counts per 10,000 with a pseudocount of 1 (log₂[(mean PMD + 1)/(mean reference + 1)]), for the 8,615 prototype genes expressed in monocytes. Agreement was measured by Spearman’s ρ between the log₂ fold change and the ExPert score (signed so that predicted loss is negative). PMD-associated genes were those in curated OXPHOS-complex, mitochondrial-biogenesis (Reactome mitochondrial translation, mitochondrial protein import, Fe–S cluster biogenesis, cristae formation) and mTORC1-signaling sets, plus 44 candidate PMD genes nominated by the clinical collaborators.

#### OXPHOS-loss score (Figure 6C)

For each cell, the score was the cosine similarity between its latent and the normalized mean of the prototypes of the complex I, III, IV and V genes (curated modules), averaged over the cell’s 30 nearest neighbors in the latent space (cosine). The score was not centered on the cohort, so higher values mean greater similarity to OXPHOS loss. Monocytes were embedded with UMAP (umap-learn; 30 neighbors, minimum distance 0.3, cosine metric) on their unit-normalized, mean-centered latents.

#### Donor clustering (Figure 6D)

For each donor, the monocyte profile was the donor’s mean latent minus the mean of the other eight donors’ mean latents. Profiles were clustered hierarchically (cosine distance, average linkage). For a model-free comparison, donors were clustered in the same way on their mean log-normalized expression (counts per 10,000, log1p, the same masked genes removed) of the 2,000 most highly variable monocyte genes. Each tree was cut into two clusters, and agreement with PMD status was measured by the adjusted Rand index (ARI), with an exact one-sided *p* value from the ARI distribution over all 126 ways to label four of the nine donors as PMD.

#### Gene-set loss scores (Figures 6E and 6F)

Gene sets were (i) the diagnostic-grade (green) genes of the Genomics England PanelApp Mitochondrial disorders panel^115^ (v10.25; 311 genes, 244 of them trained prototypes after mapping to current HGNC symbols), each assigned to the first matching MitoCarta3.0^116^ MitoPathway group (complexes I–V, coenzyme Q biosynthesis, Fe–S cluster and lipoate, mtDNA maintenance, mitochondrial translation and RNA metabolism, pyruvate dehydrogenase [PDH] and tricarboxylic acid [TCA] cycle, protein import and dynamics, small-molecule carriers); (ii) the MitoCarta3.0 OXPHOS complexes I–V (all subunits and assembly factors); (iii) each patient’s lesion gene alone; and (iv) Reactome 2024 pathways and curated modules for the stress response (integrated stress response, cap-dependent translation) and mTOR signaling. For each donor and cell type with at least 100 cells, the donor’s shift was its mean latent minus the cell-weighted mean latent of the five reference donors (for a reference donor, the shift relative to the other four was rescaled to the same five-donor baseline). Each shift was projected onto the unit mean prototype direction of the gene set and expressed in units of the standard deviation of the five reference donors’ projections. **Figure 6E** shows the PanelApp-restricted complexes, CoQ and PDH/TCA groups, their union (“all”) and the lesion genes in monocytes; **Figure 6F** shows the MitoCarta3.0 complexes, the PanelApp groups and the stress and mTOR sets in each cell type.

PMD patients were compared with reference donors by the difference between the group means divided by the reference standard deviation, tested by an exact one-sided donor-label permutation test (all 126 relabelings of four of nine donors as PMD, recomputing the reference standard deviation each time; fewer relabelings where donors with < 100 cells of a type were omitted). Individual donors were scored as z = (projection − reference mean)/reference SD, with a one-sided *p* value from a t prediction interval (4 degrees of freedom, z/√(1 + 1/5)). For reference donors this z is computed within the reference set and serves as a calibration.

### Stimulation assays

Eight patient samples (five with PMD, three controls) were processed in four batches; three of the PMD patients (P43, P54, P67) are also in the single-cell cohort. Samples were thawed and prepared for Tyto sorting as described above. Antibodies used were anti-human CD3-BV510 (BioLegend 317332), CD19-FITC (Invitrogen 11-0199-42), CD8a-PE-Cy7 (Invitrogen 25-0088-42), CD14-APC (Invitrogen 17-0149-42), and CD4-APC-eFluor 780 (Invitrogen 47-0047-41), all at 1:100. Cells were washed, resuspended in 2 mL Tyto buffer with DAPI (1:1000), and sequentially sorted into live CD4⁺ T cells (CD19⁻CD3⁺CD4⁺), CD8⁺ T cells (CD19⁻CD3⁺CD8⁺), and classical monocytes (CD19⁻CD3⁻CD14⁺).

Sorted cells were resuspended at 1×10⁶ cells/mL in cHPLM with 1% Antibiotic-Antimycotic (Gibco), supplemented with recombinant human IL-2 (50 U/mL; teceleukin) for T cells. Each population was split into two wells of a 96-well plate and rested for 30 min at 37°C, 5% CO₂. T cells were stimulated with ImmunoCult Human CD3/CD28 T Cell Activator (StemCell 10971; 25 µL/mL) and monocytes with LPS (100 ng/mL), with matched unstimulated wells receiving vehicle. Input was matched between patient and control samples as closely as possible and initial plating densities were recorded. After 24 hours, cells were collected by vigorous pipetting, while remaining adherent cells were lifted with CellStripper for 15 min at 37°C and pooled. Cells were counted, aliquoted for flow cytometry (below), and the remainder was lysed in 350 µL Buffer RLT Plus (Qiagen) with 1% β-mercaptoethanol, vortexed for at least 30 s, and stored at −80°C for RNA-seq.

### Bulk RNA-seq of sorted cells

RNA was isolated from RLT Plus lysates (approximately 8×10³–1×10⁵ cells per sample; median ∼2.3×10⁴) using the RNeasy Plus Micro Kit (Qiagen 74034). RNA was stored at −80°C, and concentration and integrity were quantified with a High Sensitivity RNA kit (Thermo Fisher) on a Qubit 2.0 Fluorometer (Thermo Fisher) and Agilent RNA ScreenTape on a 4200 TapeStation system (Agilent). Libraries were prepared using the NEBNext Single Cell/Low Input RNA Library Prep Kit for Illumina (NEB E6420S) according to the manufacturer’s instructions. Samples were grouped into three tiers by estimated input, and cDNA amplification cycles were adjusted per tier (9, 11, and 15 cycles). Libraries were indexed with NEBNext Multiplex Oligos for Illumina (Dual Index Primers Set 1; NEB E7600S), pooled, and sequenced using the NextSeq 1000/2000 XLEAP 100 cycle kit on a NextSeq 2000 sequencer (Illumina) with a paired-end read configuration (28 bp for Read 1 and 90 bp for Read 2), targeting 16 million read pairs per sample.

FASTQ files were generated using BCL Convert v4.4.7 (Illumina). Samples were processed using the nf-core/rnaseq pipeline version 3.27.0 implemented in Nextflow 26.04.4 using Singularity 3.8.1 with the minimal command: nextflow run nf-core/rnaseq-r ‘3.27.0’-profile nu_genomics –fasta ‘GRCh38-2024-A.fa’ --gtf ‘gencode.v44.primary_assembly.annotation.gtf’ --gencode true --extra_trimgalore_args ‘--length 15’, with all libraries set as unstranded (salmon^159^ gene-level quantification). Cross-population contamination was estimated by summing normalized counts of myeloid (CD14, LYZ, S100A8, S100A9, FCN1, VCAN) and T-cell (CD3E, CD3D, CD2, TRAC, CD247) marker panels, scaled to their median in monocyte and T-cell libraries respectively. Six of 48 libraries were excluded on this basis: the unstimulated and stimulated monocyte libraries from patients 75 and 83, which were mostly T-cell RNA, and both CD8⁺ T-cell libraries from patient 43 (stimulated, ∼90% myeloid signal; unstimulated, myeloid signal plus the lowest quality on every metric). The remaining monocyte libraries come from four PMD and two control patients; all remaining PMD monocyte samples were biobanked and all control samples freshly collected.

Principal-component analysis (**Figures 6G** and **6H**) was performed per cell type with PyDESeq2^145^, following DESeq2’s tximport workflow ^144,160^: per-gene, per-library normalization factors were computed once over all 48 libraries from the salmon average transcript lengths and median-of-ratios size factors and kept when the libraries were subset; genes with a mean normalized count ≥ 10 within the cell type were retained; counts were variance-stabilized (blind to the design); and PCA (centered, unscaled) was run on the 1,000 genes with the highest variance-stabilized variance. Of PC1 and PC2, the stimulation PC was the one with the higher r² for stimulation (linear model over libraries) and the disease PC was the other; PC signs were oriented so that stimulated and PMD libraries score higher. r² is the variance of the PC explained by stimulation or by PMD status. The collaborators’ R implementation (DESeq2 v1.46.0) gave the same PCs and r^2^ within 0.05.

### Flow cytometry

For each cell type and condition, matched cell numbers (up to 2×10⁴ cells per panel, scaled down for lower yields but held constant within a cell type and batch) were washed in phenol red-free HBSS and stained in 100 µL of one of two dye panels in protein-free HBSS for 25 min at 37°C, 5% CO₂: Panel 1, MitoTracker Green FM (20 nM) and TMRE (50 nM); Panel 2, MitoSOX Green (1 µM) and CellROX Orange (5 µM). Cells were washed once in HBSS (500g x 5 min at RT), resuspended in room-temperature flow buffer with DAPI (1:1000), kept at RT, and acquired on a BD LSRFortessa within 1 hour of staining. Residual fluorescence from sort antibodies was assessed at acquisition; CD8 signal, but not CD4 or CD14, persisted after 24 hours of culture. Dye MFIs were quantified on live (DAPI⁻) singlets in FlowJo v10.

For **Figure 6I**, the change on stimulation was computed per patient as the percent change of the geometric MFI of stimulated over unstimulated monocytes, for MitoTracker Green (mitochondrial mass) and CellROX Orange (reactive oxygen species). Monocytes of P75 and P83 were excluded before analysis for T-cell contamination (the same wells whose bulk libraries were excluded, above), leaving two control and four PMD patients. Groups were compared by a two-sided Student’s t test.

