## Supplementary material for "ExPert infers causal perturbations from cell states across biological and clinical contexts": SI Figures, Table Captions, Notes

# SI 1

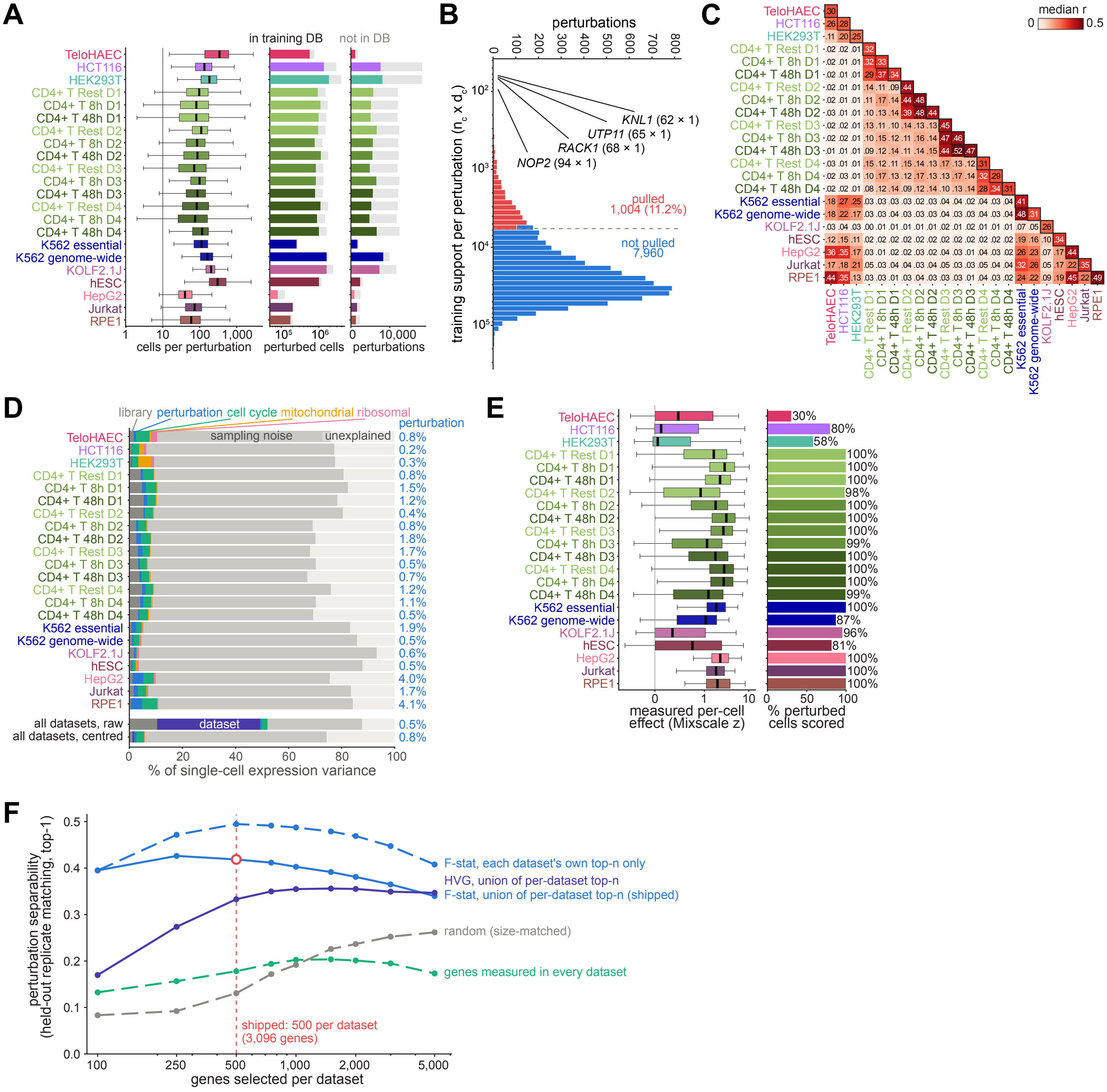
A
in training DB
not in DB
TeloHAEC
HCT116
HEK293T
CD4+ T Rest D1
CD4+ T 8h D1
CD4+ T 48h D1
CD4+ T Rest D2
CD4+ T 8h D2
CD4+ T 48h D2
CD4+ T Rest D3
CD4+ T 8h D3
CD4+ T 48h D3
CD4+ T Rest D4
CD4+ T 8h D4
CD4+ T 48h D4
K562 essential
K562 genome-wide
KOLF2.1J
hESC
HepG2
Jurkat
RPE1
1 10 100 1,000
cells per perturbation
10⁵ 10⁶
perturbed cells
0 10,000
perturbations
B
perturbations
0 100 200 300 400 500 600 700 800
training support per perturbation (n_c x d_c)
KNL1 (62 × 1)
UTP11 (65 × 1)
RACK1 (68 × 1)
NOP2 (94 × 1)
pulled
1,004 (11.2%)
not pulled
7,960
C
median r
0 0.5
D
library perturbation cell cycle mitochondrial ribosomal
perturbation
sampling noise
unexplained
0.8% 0.2% 0.3% 0.8% 1.5% 1.2% 0.4% 0.8% 1.8% 1.7% 0.5% 0.7% 1.2% 1.1% 0.5% 1.9% 0.5% 0.6% 0.5% 4.0% 1.7% 4.1%
all datasets, raw
all datasets, centred
dataset
0.5%
0.8%
0 20 40 60 80 100
% of single-cell expression variance
E
30% 80% 58% 100% 100% 100% 98% 100% 100% 100% 99% 100% 100% 100% 99% 100% 87% 96% 81% 100% 100% 100%
0 1 10
measured per-cell effect (Mixscale z)
0 50 100
% perturbed cells scored
F
perturbation separability (held-out replicate matching, top-1)
F-stat, each dataset's own top-n only
HVG, union of per-dataset top-n
F-stat, union of per-dataset top-n (shipped)
random (size-matched)
genes measured in every dataset
shipped: 500 per dataset
(3,096 genes)
100 250 500 1,000 2,000 5,000
genes selected per dataset

**Figure S1. Composition and properties of the training corpus, related to Figure 1**

(A) Cells per perturbation in each dataset of the store. Left, quality-control-passed perturbed cells per perturbation (log scale; box, median and interquartile range; whiskers, 1.5x interquartile range; outliers not shown); dashed line, the 10-cell minimum a perturbation needs in a context to count toward selection. Middle, perturbed cells and right, perturbations per dataset, split into those in the training database (colored) and not (gray). Datasets are colored and ordered by context as in **Figure 1A**; $CD4^+$ T cell datasets are labeled by stimulation condition and donor (D1–D4).

(B) Training support of the 8,964 selected perturbations, defined as training cells by number of datasets with training cells (log scale). Red, the 1,004 perturbations (11.2%) with the lowest support, whose prototypes are pulled toward the mean of their prior neighbors during training (**Figure S2**); blue, the remaining 7,960; dashed line, cutoff. The four lowest-support perturbations are labeled with training cells by datasets.

(C) Similarity of the response to the same perturbation across datasets. Pseudobulk profiles (log counts per million, minus the dataset's control profile, 3,096 input genes) were compared by Pearson correlation between the two datasets' responses to each shared perturbation, over genes measured in both; each cell gives the median over perturbations that are reproducible in both datasets (correlation ≥ 0.2 between two random halves of the perturbation's cells). Diagonal (boxed), split-half reliability: median correlation between two random halves of each reproducible perturbation within the dataset. The matrix is symmetric; the lower triangle is shown. $CD4^+$ T cell datasets are grouped by donor.

(D) Sources of single-cell expression variance. Per dataset, up to 300 selected perturbations with ≥20 cells (20 cells each) and 2,000 control cells, 3,096 input genes, log counts per 10,000. Variance was partitioned sequentially: library depth (regression on log total counts over the input genes), perturbation identity (one-way analysis of variance on the residual, adjusted for the number of groups), cell cycle (regression on S and G2/M scores from the cell-cycle genes of Tirosh et al.[163] in the input panel), and the fractions of counts from mitochondrial and from ribosomal protein genes. The remainder is split into sampling noise, estimated by count splitting (each cell's counts divided binomially into two halves; variance not shared between the halves; an upper bound under the log transform), and unexplained variance. Bottom, all datasets pooled with equal cells per dataset, before (raw) and after subtracting each dataset's control mean (centered, as for the model input), with dataset entered after library depth. Right, percentage explained by perturbation.

(E) Measured per-cell effect size (Mixscale[120] score: projection of a cell onto its perturbation's differential-expression direction, in units of control-cell standard deviations; symmetric log scale). Left, distribution per dataset (20,000 cells sampled per dataset; box, median and interquartile range; whiskers, 5th–95th percentiles). Right, percentage of perturbed cells with a score (perturbations with fewer than five differentially expressed genes are not scored).

(F) Selection of the input genes. Perturbation separability (held-out replicate matching: genes are selected on a random half of each dataset's perturbations, and each held-out pseudobulk replicate is matched by cosine similarity to its own perturbation among all reference profiles of the dataset; top-1 accuracy, mean over 22 datasets) against the number of genes selected per dataset. F-stat, ratio of between- to within-perturbation variance per gene; HVG, highly variable genes (seurat_v3); random, size-matched to the F-stat union. Solid lines, a single gene set shared by all datasets (union of per-dataset sets); dashed lines, sets that are not such a union (each dataset's own genes only, genes measured in every dataset, random). Open circle and dashed vertical line, the configuration used (500 genes per dataset; a union of about 3,170 genes in the sweep, 3,096 in the shipped input panel).

# SI 2

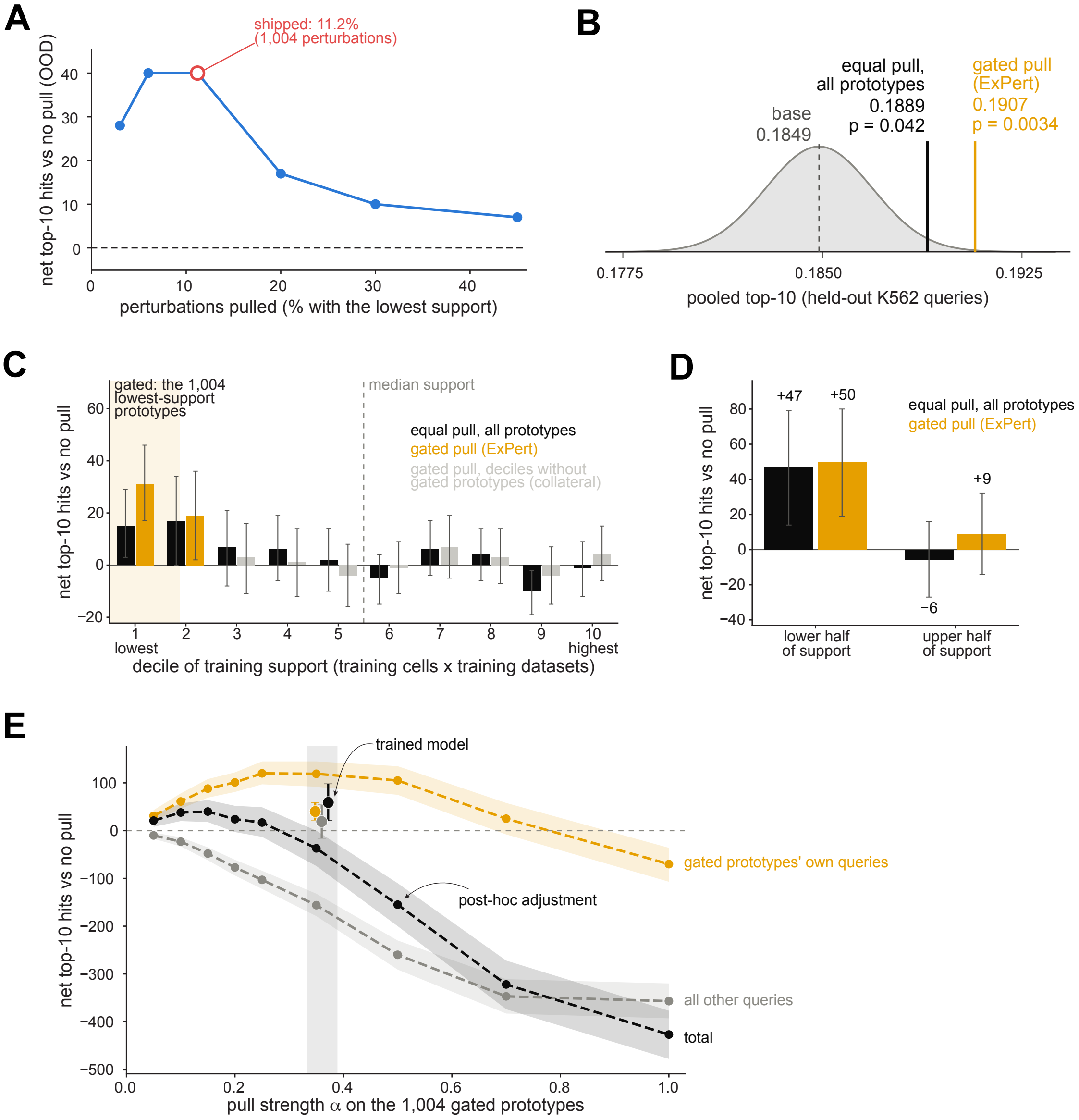

A
shipped: 11.2%
(1,004 perturbations)
net top-10 hits vs no pull (OOD)
perturbations pulled (% with the lowest support)
B
equal pull,
all prototypes
0.1889
p = 0.042
gated pull
(ExPert)
0.1907
p = 0.0034
base
0.1849
0.1775
0.1850
0.1925
pooled top-10 (held-out K562 queries)
C
gated: the 1,004
lowest-support
prototypes
median support
equal pull, all prototypes
gated pull (ExPert)
gated pull, deciles without
gated prototypes (collateral)
net top-10 hits vs no pull
lowest
highest
decile of training support (training cells x training datasets)
D
+47
+50
+9
−6
equal pull, all prototypes
gated pull (ExPert)
net top-10 hits vs no pull
lower half
of support
upper half
of support
E
trained model
post-hoc adjustment
gated prototypes' own queries
all other queries
total
net top-10 hits vs no pull
pull strength α on the 1,004 gated prototypes

**Figure S2. The prior pull during training: size of the pulled set, its effect, its gain by training support, and the pull applied after rather than during training, related to Figure 3**

(A) Size of the pulled set. Offline sweep on the model trained without the pull (fold-4 held-out contexts, 10,061 query groups): for each of six set sizes, the lowest-support fraction of perturbations was blended toward its prior-predicted position, and the net change in top-10 hits relative to no pull is shown at the best of nine blend strengths. Open circle, the size used (11.2%, 1,004 perturbations).

(B) Effect of pulling prototypes toward their prior-graph neighbors during training. Pooled top-10 recall on 10,061 held-out K562 queries (trained perturbations) for models trained with K562 held out: without pull (base; dashed line), with pull restricted to the gated prototypes (**Figure S1B**) (ExPert; orange), and with equal pull on all prototypes (black); n = 1 training run per configuration. Gray distribution, run-to-run variation around base: normal with SD = 0.002, estimated by a paired bootstrap over queries between two independently trained replicate runs of one configuration. p-values, two-sided Z-test of each configuration against the run-to-run distribution (gated pull, $p$ = 0.0034; equal pull, $p$ = 0.042).

(C) Net change in top-10 hits relative to the model trained without the pull, on 10,061 held-out K562 pseudobulk queries of trained perturbations (models trained with K562 held out), by decile of the training support of each query's perturbation (training cells by number of training datasets, ranked as the gate of the trained model ranks them), from 1 (lowest) to 10 (highest). Equal pull on all prototypes (black) and gated pull (ExPert; orange). Shaded, the queries of the 1,004 gated (lowest-support) prototypes, which fill decile 1 and 37% of decile 2 (1,383 queries). Gray, gated pull in deciles without gated prototypes, where any change is collateral: the whole network is trained with the pull (see E). Dotted line, the median support.

(D) The same summed over the lower and upper five deciles. Equal pull: lower half +47 (95% CI 14–79), upper half −6 (−27 to 16); gated pull: +50 (19–80) and +9 (−14 to 32).

(E) Net change in top-10 hits relative to the model trained without the pull, on the same queries, split into the queries of perturbations whose prototype is among the 1,004 gated (pulled) prototypes (orange), all other queries (gray), and in total (black). Dashed lines, post-hoc adjustment: after training without the pull, the gated prototypes are blended toward their prior-predicted positions with strength α (α = 0.05–1.0). Points, the model trained with the gated pull, at its implied strength (median α = 0.36; shaded, interquartile range over the pulled prototypes). At the post-hoc strength closest to the one training reached (α = 0.35; trained median 0.36), the post-hoc adjustment gains 119 hits on the gated prototypes' own queries but loses 156 elsewhere (net −37; 95% CI −74 to 0); the trained model gains 40 on the gated prototypes' queries and 19 elsewhere (net +59; 95% CI 21–98). Light post-hoc adjustments are net positive (+21 to +40 at α = 0.05–0.15). Bands and error bars, 95% confidence intervals (bootstrap over queries).

In (C) and (D), error bars, 95% confidence intervals (bootstrap over queries, 2,000 resamples).

# SI 3

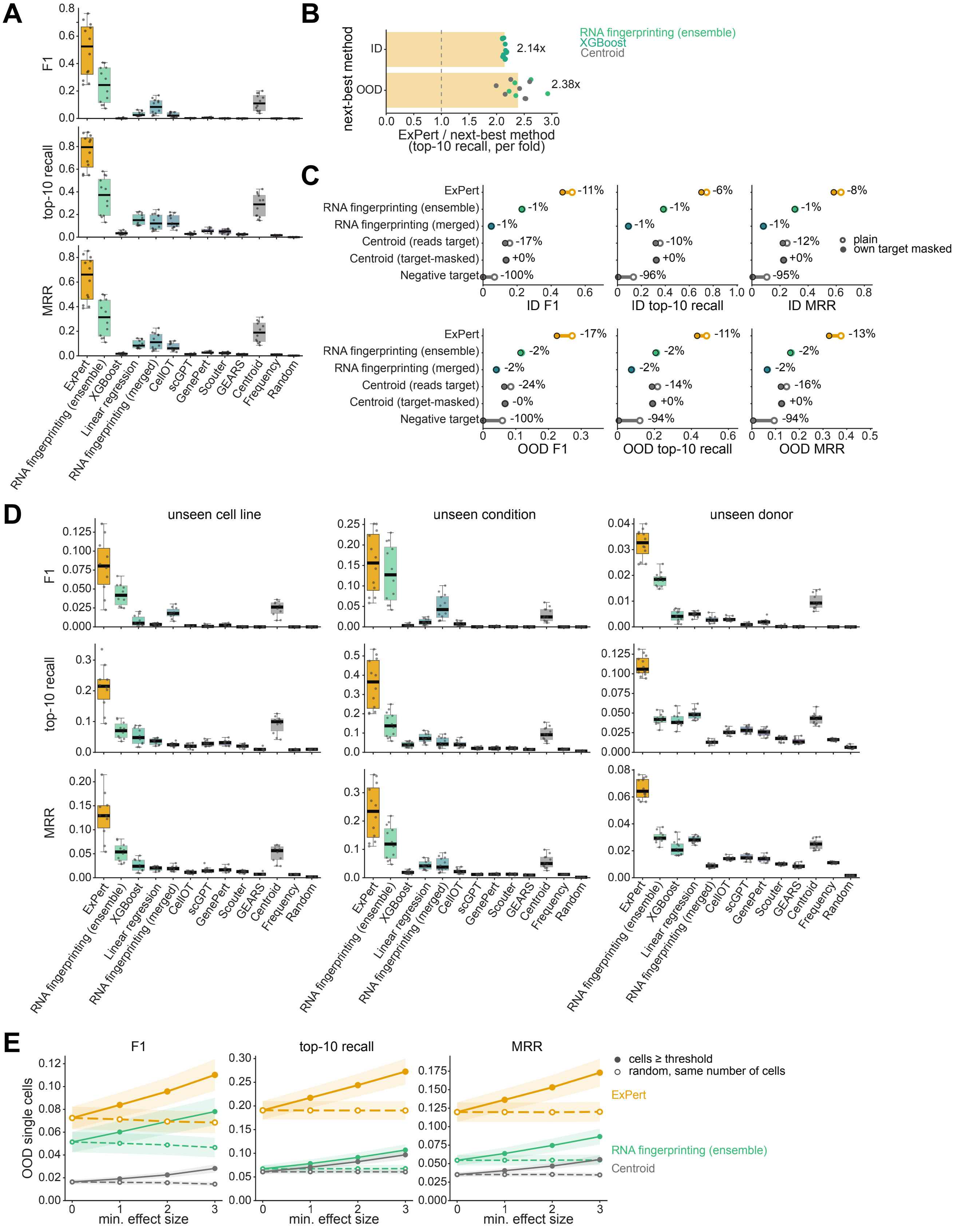
A
F1
top-10 recall
MRR
ExPert
RNA fingerprinting (ensemble)
XGBoost
Linear regression
RNA fingerprinting (merged)
CellOT
scGPT
GenePert
Scouter
GEARS
Centroid
Frequency
Random
B
next-best method
ID
OOD
2.14x
2.38x
RNA fingerprinting (ensemble)
XGBoost
Centroid
ExPert / next-best method
(top-10 recall, per fold)
C
ExPert
RNA fingerprinting (ensemble)
RNA fingerprinting (merged)
Centroid (reads target)
Centroid (target-masked)
Negative target
-11%
-1%
-1%
-17%
+0%
-100%
-6%
-1%
-1%
-10%
+0%
-96%
-8%
-1%
-1%
-12%
+0%
-95%
plain
own target masked
ID F1
ID top-10 recall
ID MRR
-17%
-2%
-2%
-24%
-0%
-100%
-11%
-2%
-2%
-14%
+0%
-94%
-13%
-2%
-2%
-16%
+0%
-94%
OOD F1
OOD top-10 recall
OOD MRR
D
unseen cell line
unseen condition
unseen donor
F1
top-10 recall
MRR
E
F1
top-10 recall
MRR
OOD single cells
min. effect size
cells ≥ threshold
random, same number of cells
ExPert
RNA fingerprinting (ensemble)
Centroid

**Figure S3. Retrieval by held-out context, robustness to target-gene masking, and the effect of pooling and knockdown strength, related to Figure 2**

(A) OOD performance on pseudobulks for held-out conditions (n = 12 dataset by fold combinations); held-out cell lines and donors are shown in **Figure 2C**. Rows show F1, top-10 recall, and MRR over 1,157 candidate perturbations, with each perturbation weighted equally. Boxes show the median and interquartile range (IQR); whiskers extend to 1.5× IQR; points, individual dataset by fold combinations.

(B) Per-fold ratio of ExPert's top-10 recall to that of the best other method in the same fold, in distribution (ID) and out of distribution (OOD), on single cells (weighted by knockdown effect size, each perturbation weighted equally; mean over the fold's dataset combinations). Bars show the median across the 12 folds (value beside each bar); points, individual folds, colored by the method that was second best in that fold; dotted line, parity.

(C) Effect of masking the CRISPRi target gene in each query, on pseudobulks over all 1,157 perturbations. For each perturbation, the expression of its own target gene in the query cells was removed (ExPert, set to zero in the control-centered input; Fingerprint, SCTransform residual set to zero; Centroid and Negative target, set to the dataset's control mean); trained models were unchanged. Open circles, unmasked; filled circles, masked; numbers, relative change. Centroid (reads target), nearest-centroid baseline over all input genes; Centroid (target-masked), the same baseline with each perturbation's own target gene excluded; Negative target, ranks perturbations by the drop of their target gene below the control mean (perturbations whose target is not an input gene are scored on a random non-target input gene). Values are means over n = 230 (ID) and n = 34 (OOD) dataset by fold combinations. For Fingerprint, perturbations whose target gene is an input were rescored on all OOD cells and on 5,000 randomly sampled ID cells per fold; all other perturbations, which masking cannot affect, keep their published scores.

(D) **Figure 2C** on single cells: OOD performance per held-out dataset by fold combination, split by held-out context (unseen cell line, n = 10; unseen condition, n = 12; unseen donor, n = 12), scored on single cells with each cell weighted by its knockdown effect size. Rows, F1, top-10 recall and MRR over 1,157 candidate perturbations; boxes and points as in (A).

(E) Control for **Figure 2D**. OOD single-cell performance on the cells at or above each minimum knockdown effect size (0, 1, 2 and 3) (solid lines, filled circles) and on a random sample of the same number of cells drawn regardless of effect size (dashed lines, open circles; mean of five draws), for ExPert, Fingerprint (ensemble), and the nearest-centroid baseline. Metrics are computed per perturbation with cells weighted equally. Data are represented as mean ± SEM over n = 34 OOD dataset by fold combinations, each with at least 30 cells at every threshold.

# SI 4

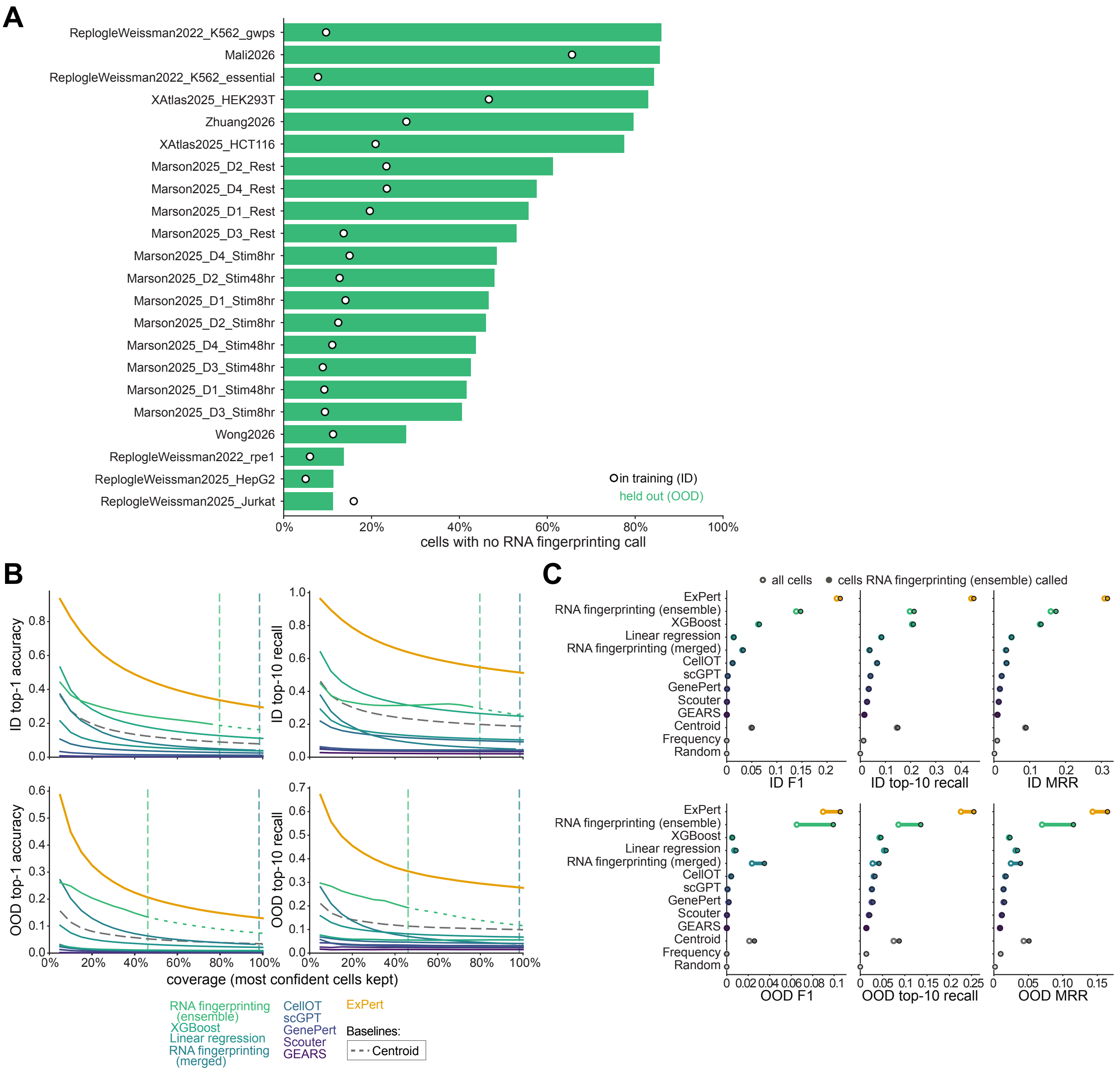
A
ReplogleWeissman2022_K562_gwps
Mali2026
ReplogleWeissman2022_K562_essential
XAtlas2025_HEK293T
Zhuang2026
XAtlas2025_HCT116
Marson2025_D2_Rest
Marson2025_D4_Rest
Marson2025_D1_Rest
Marson2025_D3_Rest
Marson2025_D4_Stim8hr
Marson2025_D2_Stim48hr
Marson2025_D1_Stim8hr
Marson2025_D2_Stim8hr
Marson2025_D4_Stim48hr
Marson2025_D3_Stim48hr
Marson2025_D1_Stim48hr
Marson2025_D3_Stim8hr
Wong2026
ReplogleWeissman2022_rpe1
ReplogleWeissman2025_HepG2
ReplogleWeissman2025_Jurkat
in training (ID)
held out (OOD)
0%
20%
40%
60%
80%
100%
cells with no RNA fingerprinting call
B
ID top-1 accuracy
ID top-10 recall
OOD top-1 accuracy
OOD top-10 recall
coverage (most confident cells kept)
RNA fingerprinting (ensemble)
XGBoost
Linear regression
RNA fingerprinting (merged)
CellOT
scGPT
GenePert
Scouter
GEARS
ExPert
Baselines:
Centroid
C
all cells
cells RNA fingerprinting (ensemble) called
ExPert
RNA fingerprinting (ensemble)
XGBoost
Linear regression
RNA fingerprinting (merged)
CellOT
scGPT
GenePert
Scouter
GEARS
Centroid
Frequency
Random
ID F1
ID top-10 recall
ID MRR
OOD F1
OOD top-10 recall
OOD MRR

**Figure S4. Fingerprint assignment comparison, related to Figure 2**

(A) Percentage of query cells that RNA Fingerprinting (ensemble) left unassigned (no per-dataset fingerprint model called a perturbation), per dataset, when the dataset was held out (OOD; bars) and when it was in training (ID; open circles), pooled over folds.

(B) Retrieval as a function of coverage on single cells, in distribution (ID; top) and out of distribution (OOD; bottom). Within each dataset by fold combination, each method's query cells were ranked by its confidence (difference between the top-1 and top-2 log probabilities), with cells that Fingerprint left unassigned ranked last; at each coverage, the most confident fraction of cells was kept. Left, top-1 accuracy; right, top-10 recall; both pooled over the kept cells of all dataset by fold combinations, each cell weighted by its knockdown effect size. Dashed vertical lines, the fraction of cells Fingerprint (ensemble; 79.7% ID, 46.1% OOD) and Fingerprint (merged; 98.5% ID, 98.3% OOD) assigned; beyond it, curves are dotted (unassigned cells). Centroid, dashed gray. Random and Frequency are not shown, as their scores do not depend on the cell.

(C) Performance on all cells (open circles) and on only the cells that Fingerprint (ensemble) assigned (filled circles), ID (top) and OOD (bottom). F1, top-10 recall, and MRR over 1,157 candidate perturbations, per perturbation (cells weighted by knockdown effect size, each perturbation weighted equally). Values are means over n = 230 (ID) and n = 34 (OOD) dataset by fold combinations.

## SI 5

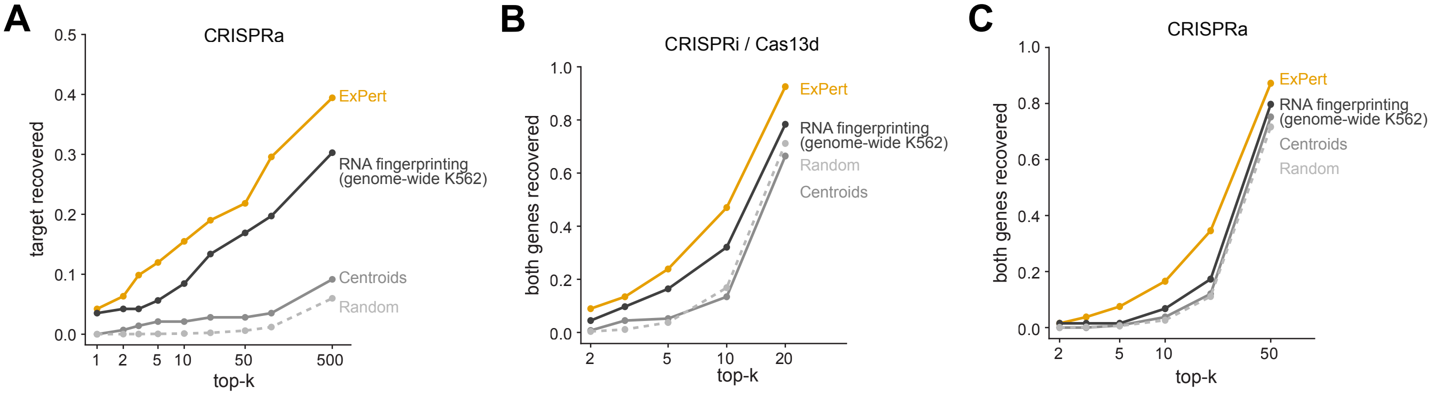

A
CRISPRa
target recovered
top-k
ExPert
RNA fingerprinting (genome-wide K562)
Centroids
Random
B
CRISPRi / Cas13d
both genes recovered
top-k
ExPert
RNA fingerprinting (genome-wide K562)
Random
Centroids
C
CRISPRa
both genes recovered
top-k
ExPert
RNA fingerprinting (genome-wide K562)
Centroids
Random

**Figure S5. Recovery of single CRISPRa perturbations and extended double perturbation recovery, pooled and per screen, related to Figure 3**

(A) Recovery of single CRISPRa perturbations. ExPert was trained on loss-of-function screens only, so gain-of-function single perturbations are themselves a transfer. Fraction of single perturbations whose targeted gene ranks within the top k of the 8,347 candidate perturbations of **Figures 3H** and **3I** (ranked from the gain end for ExPert). ExPert (orange); RNA fingerprinting custom-trained on genome-wide K562 Perturb-seq (dark gray); nearest centroid of each perturbation's training cells (gray); random ranking (light gray, dotted). n = 142 single perturbations whose targeted gene is a candidate, from two CRISPRa screens (Norman et al.[62], n = 78 of 105; ProPer-seq[63], n = 64 of 84).

(B) Recovery of combinatorial loss-of-function perturbations when the candidates are restricted to the genes targeted in each screen's own double perturbations. Fraction of double perturbations for which both targeted genes rank within the top k ($k \geq 2$) of 20 (CRISPRi screen[60]) or 24 (Cas13d screen[61]) candidate perturbations scorable by all methods; curves end at the largest evaluated k (2, 3, 5, 10, 20, 50) within the smaller candidate set (k = 20 here, k = 50 in E). Methods and colors as in (A). n = 134 double perturbations from a CRISPRi screen (Replogle et al.[60], n = 10) and a Cas13d screen (CaRPool-seq[61]; n = 124), as in **Figure 3H**.

(C) Recovery of combinatorial gain-of-function perturbations, as in (C), among 56 (Norman et al.[62]) or 64 (ProPer-seq[63]) candidate perturbations. n = 133 double perturbations from the CRISPRa screens of (A) (Norman et al.[62], n = 77; ProPer-seq[63], n = 56), as in **Figure 3I**.

None of the screens were used in training.

## SI 6

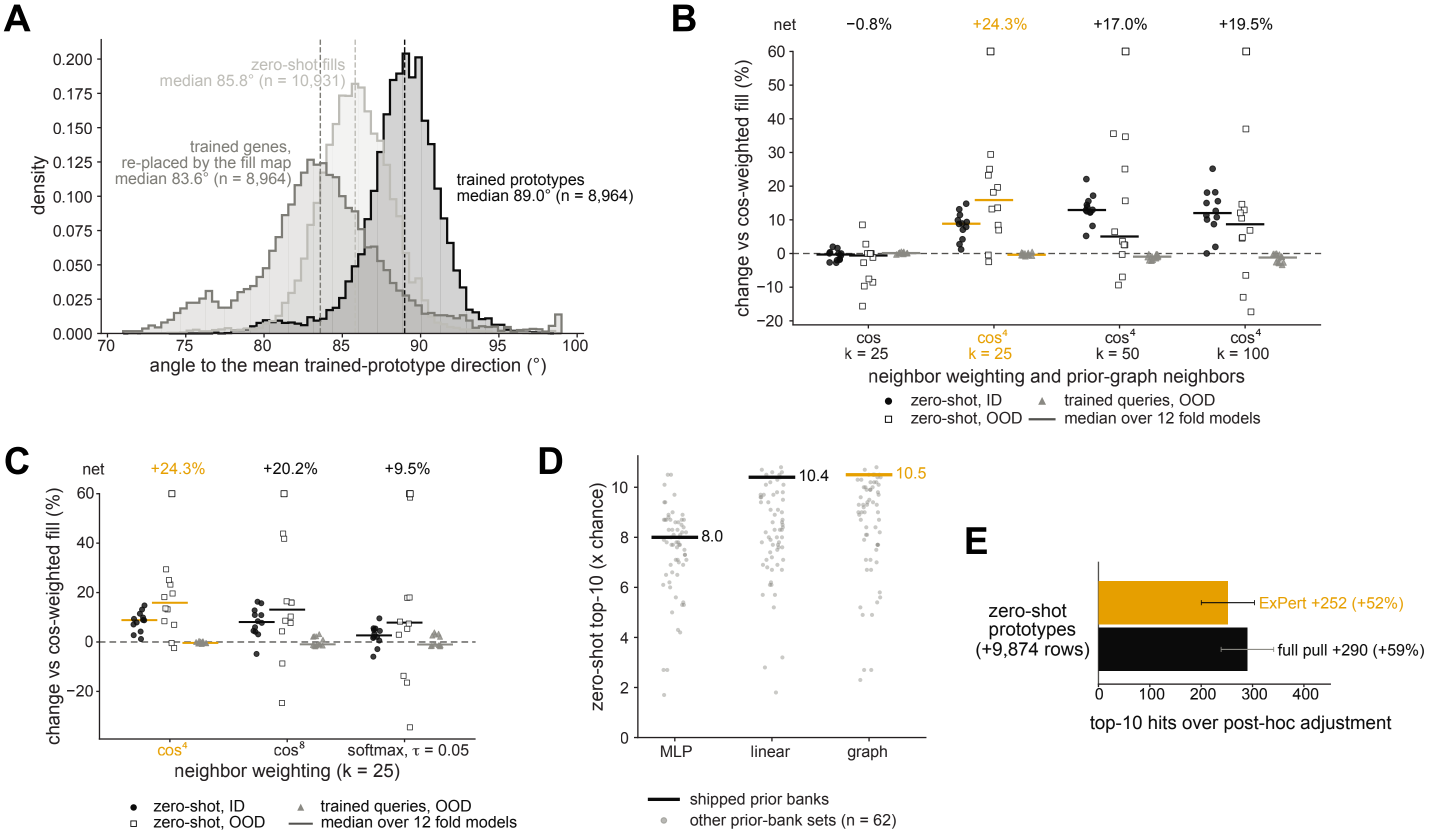

A
zero-shot fills
median 85.8° (n = 10,931)
trained genes,
re-placed by the fill map
median 83.6° (n = 8,964)
trained prototypes
median 89.0° (n = 8,964)
density
angle to the mean trained-prototype direction (°)
B
net −0.8% +24.3% +17.0% +19.5%
change vs cos-weighted fill (%)
cos k = 25
cos⁴ k = 25
cos⁴ k = 50
cos⁴ k = 100
neighbor weighting and prior-graph neighbors
zero-shot, ID
zero-shot, OOD
trained queries, OOD
median over 12 fold models
C
net +24.3% +20.2% +9.5%
change vs cos-weighted fill (%)
cos⁴
cos⁸
softmax, τ = 0.05
neighbor weighting (k = 25)
zero-shot, ID
zero-shot, OOD
trained queries, OOD
median over 12 fold models
D
zero-shot top-10 (x chance)
8.0
10.4
10.5
MLP
linear
graph
shipped prior banks
other prior-bank sets (n = 62)
E
zero-shot
prototypes
(+9,874 rows)
ExPert +252 (+52%)
full pull +290 (+59%)
top-10 hits over post-hoc adjustment

**Figure S6. Zero-shot prototype geometry, fill weighting and map, and training with zero-shot prototypes as negatives, related to Figure 3**

(A) Angle of each prototype to the mean direction of the trained prototypes in the latent space of the final model: trained prototypes (n = 8,964; median 89.0°), zero-shot prototypes placed by the graph fill from their trained prior-graph neighbors (fills; n = 10,931; median 85.8°), and the trained perturbations placed by the same map as if they were unseen (n = 8,964; median 83.6°). Dashed lines, medians. Trained perturbations placed by the map shift at least as far as the fills (median 83.6° against 85.8°), so the shift reflects averaging over neighbors rather than inaccurate priors.

(B) Number of prior-graph neighbors k of the graph fill, which places each untrained perturbation at the weighted mean of the prototypes of its k nearest trained neighbors in each prior graph, each neighbor weighted by the fourth power of its prior similarity ($\cos^4$; averaged over the prior banks). Fills were rebuilt on the frozen fold models (n = 12; **Table S5**) and scored as in **Figure 3B**; each value is the change relative to the fill weighted by cosine similarity at k = 25 on the same model. Filled circles, zero-shot top-10 recall of unseen perturbations in contexts seen in training (ID); open squares, in the held-out context (OOD); gray triangles, top-10 recall of held-out queries of trained perturbations with the fills among the candidates (OOD). Points, fold models; bars, medians; numbers above the panel (net), the sum of the three medians. Left, the cosine-weighted fill stored in each model (cos, k = 25; net −0.8%), whose spread around zero is the noise of rebuilding the fill. $\cos^4$ at k = 25 (orange) was chosen by the net change: +8.8%, +15.9% and −0.4% (net +24.3%) vs +12.9%, +5.0% and −0.9% at k = 50 (+17.0%) and +12.0%, +8.7% and −1.2% at k = 100 (+19.5%). Values beyond the axis (one fold model, fold 8) are drawn at its edge and printed.

(C) Neighbor weighting of the graph fill at k = 25, relative to cosine weighting, drawn as in (B); beyond the axis are fold 8 (all three weightings) and, for softmax, also fold 11. Net change: $\cos^4$ +24.3%, $\cos^8$ +20.2%, softmax +9.5%. $\cos^4$ (orange; shipped): zero-shot +8.8% ID (12 of 12 fold models improved; two-sided sign test, $p = 0.0005$) and +15.9% OOD (10 of 12; $p = 0.04$), trained perturbations −0.4%; $\cos^8$: +8.1% (11 of 12) and +13.1% (10 of 12), −0.9%; softmax over cosine similarity (temperature $\tau$ = 0.05): +2.7% (10 of 12) and +7.8% (9 of 12), −1.0%. Weightings tested on the fold-4 model only (K562 held out; aux): $\cos^2$, softmax with $\tau$ = 0.02, cosine times neighbor support (−15% and −38% zero-shot ID and OOD; squared, −40% and −79%) and cosine times held-out recall (+2.5% and −5.7%; squared, −6.5% and −5.7%).

(D) Zero-shot recovery by fill map: a multilayer perceptron (MLP), a linear map (ridge regression, $\alpha$ = 0.5) and the graph fill (orange), each mapping the prior banks onto the trained prototypes. Offline scan made before the weighting in (C) was chosen, so the graph fill here is cosine-weighted at k = 25. Fold-4 model trained without priors (K562 held out); zero-shot score, top-10 recall of the unseen perturbations among the untrained prototypes relative to chance (one pooled query per perturbation), so values are not on the scale of **Figure 3B**. Lines and values, the shipped prior banks (text, DepMap[46,47] and STRING[51,52]; 8.0×, 10.4× and 10.5×); gray points, each of the other 62 combinations of prior banks (combinations with co-expression, which is built from the screens, are excluded).

(E) Training with the zero-shot prototypes as negatives reduces their cost to trained perturbations. When the 9,874 zero-shot prototypes of perturbations measured in a screen (graph fill, $\cos^4$, k = 25) join the 8,964 trained prototypes as candidates, held-out queries of trained perturbations lose top-10 hits (10,061 queries; models trained with K562 held out); a model trained without priors and filled after training (post-hoc adjustment) loses 488. Bars, top-10 hits recovered over the post-hoc adjustment by models trained with the zero-shot prototypes as negatives: ExPert, whose pull is gated to the lowest-support prototypes (orange; +252, 52% of the post-hoc loss), and a model pulling every prototype with equal weight (full pull; black; +290, 59%). All are filled by the same graph map from one shared set of prior banks. Error bars, 95% confidence intervals: the bootstrap intervals (2,000 resamples over queries) of each model and of the post-hoc adjustment, combined in quadrature.

# SI 7

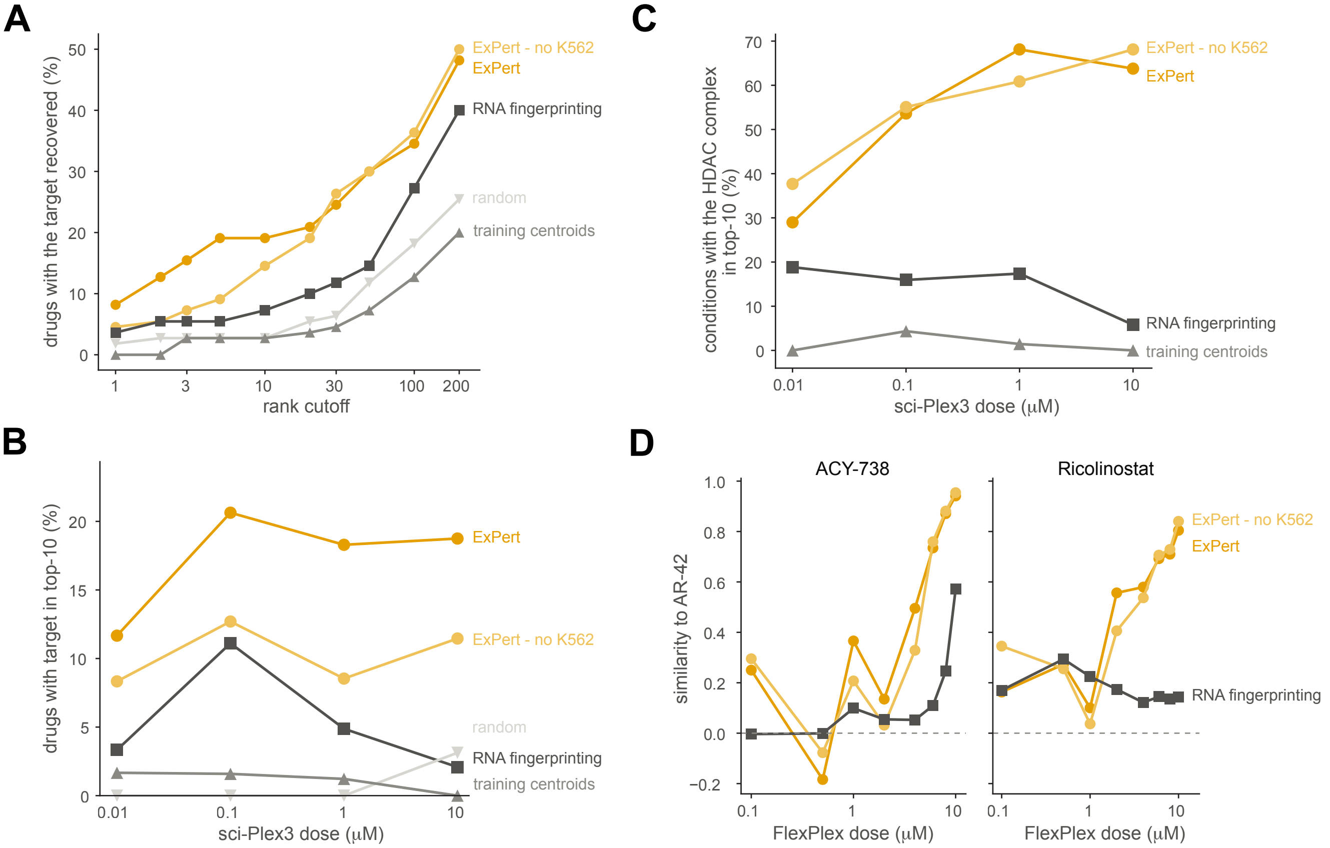

A
drugs with the target recovered (%)
rank cutoff
1
3
10
30
100
200
0
10
20
30
40
50
ExPert - no K562
ExPert
RNA fingerprinting
random
training centroids
B
drugs with target in top-10 (%)
sci-Plex3 dose (μM)
0.01
0.1
1
10
0
5
10
15
20
ExPert
ExPert - no K562
random
RNA fingerprinting
training centroids
C
conditions with the HDAC complex in top-10 (%)
sci-Plex3 dose (μM)
0.01
0.1
1
10
0
10
20
30
40
50
60
70
ExPert - no K562
ExPert
RNA fingerprinting
training centroids
D
ACY-738
Ricolinostat
similarity to AR-42
FlexPlex dose (μM)
0.1
1
10
−0.2
0.0
0.2
0.4
0.6
0.8
1.0
ExPert - no K562
ExPert
RNA fingerprinting

**Figure S7. Drug-target recovery by an ExPert model trained without any K562 data, related to Figure 4**

**Figure 4B–E** redrawn including an ExPert model trained without any K562 data (“ExPert - no K562” alongside ExPert v0.1.0: the same architecture and label space, trained with the two K562 screens[19] (ReplogleWeissman2022 K562 genome-wide and essential) held out, so the model has seen no K562 cell. The perturbations themselves are not unseen: 3,800 of the 3,802 candidate genes have training cells from other screens. ExPert - no K562 is scored on exactly the conditions of **Figure 4** (the responsiveness of each condition is judged in the v0.1.0 embedding), so the ExPert v0.1.0, RNA fingerprinting, training-centroid and random (A, B) curves are identical to **Figure 4** and only the light-orange curve is new. Light orange, ExPert - no K562; dark orange, ExPert (v0.1.0); other colors as in **Figure 4**.

(A) sci-Plex3[21] annotated drug-target recovery as a function of rank cutoff k, over the same 110 compounds as **Figure 4B**.

(B) The same recovery at k = 10 by labelled dose (n = 60, 63, 82 and 96 drugs at 0.01, 0.1, 1 and 10 μM, as in **Figure 4C**).

(C) HDAC inhibitors, scored against the 48-gene HDAC machinery over the same 92 drug by dose pairs and 69 conditions per dose as **Figure 4D**.

(D) Similarity of ricolinostat and ACY-738 to AR-42 in the FlexPlex screen of K562 cells[38], computed as in **Figure 4E**.

Statistics. Comparisons of the top-10 hit use an exact two-sided McNemar test on the discordant drugs; rank comparisons use a one-sided paired sign-flip permutation test on $\log_{10}$ rank (20,000 permutations). No error bars are plotted. Replicates as in **Figure 4**.

## SI 8

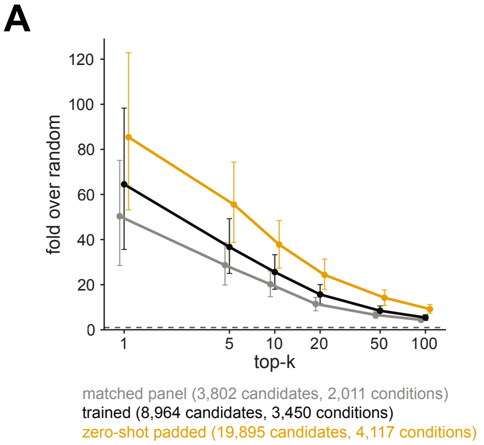

A
fold over random
120
100
80
60
40
20
0
1
5
10
20
50
100
top-k
matched panel (3,802 candidates, 2,011 conditions)
trained (8,964 candidates, 3,450 conditions)
zero-shot padded (19,895 candidates, 4,117 conditions)


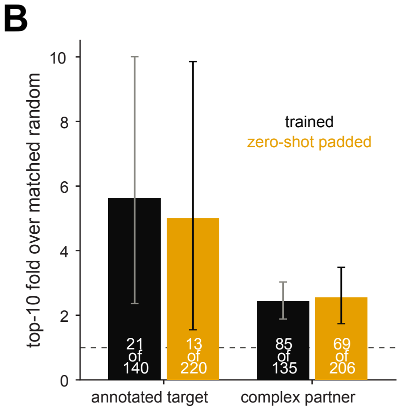

B
top-10 fold over matched random
10
8
6
4
2
0
trained
zero-shot padded
21 of 140
13 of 220
85 of 135
69 of 206
annotated target
complex partner

**Figure S8. Drug-target recovery with zero-shot prototypes added to the candidate bank, as fold over random, on sci-Plex3 and Tahoe-100M, related to Figure 4**

(A) sci-Plex3[21]: recovery of each condition's annotated drug targets in the top k (MT-masked pseudobulk, deflated readout, the better of the loss and gain tails), as fold over random, for three candidate pools: the panel matched to fingerprinting (3,802 candidates; gray), the trained prototypes (8,964; black), and the trained prototypes padded with the zero-shot prototypes (19,895; orange). Random is exact per condition, the probability that a uniform ranking places at least one of the drug's annotated targets in the pool within the top k, averaged over each pool's own scorable conditions; pools that can score different numbers of conditions (2,011, 3,450 and 4,117, from 84, 144 and 172 drugs) are therefore comparable. Dotted line, random (fold = 1). At the top 10, padding raises recovery from 25.6- to 37.8-fold over random. Error bars, 95% bootstrap intervals over drugs (2,000 resamples).

(B) Tahoe-100M[64], drugs of known mechanism: the fraction of each drug's responsive conditions that place an annotated target (left) or a complex partner of a target (right) in the top 10, relative to size- and popularity-matched random gene sets (as in **Figure 4F**), for the trained bank (black) and the zero-shot-padded bank (orange). Numbers in bars, drugs with at least one such condition, of the drugs scored. Dotted line, matched random (fold = 1). Padding makes 80 (annotated targets) and 71 (complex partners) more drugs scorable but recovers fewer (13 against 21 for annotated targets, 69 against 85 for complex partners), at a similar fold over the null (5.0× against 5.6×, and 2.5× against 2.4×). Error bars, 95% bootstrap intervals over drugs (2,000 resamples).

Zero-shot prototypes are the shipped graph fill ($cos^4$ weights, k = 25; **Figure S6**).

## SI 9

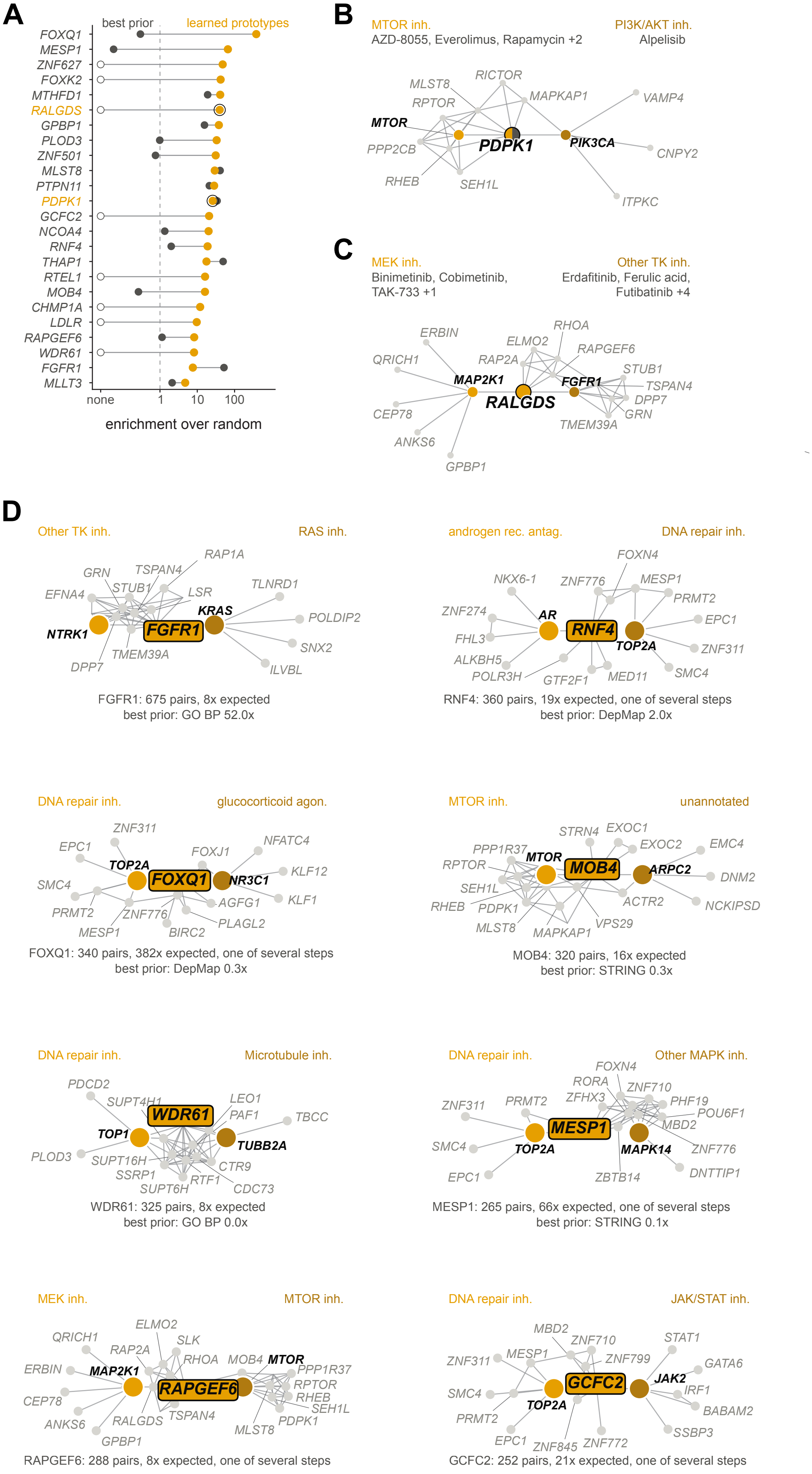
A
best prior
learned prototypes
FOXQ1
MESP1
ZNF627
FOXK2
MTHFD1
RALGDS
GPBP1
PLOD3
ZNF501
MLST8
PTPN11
PDPK1
GCFC2
NCOA4
RNF4
THAP1
RTEL1
MOB4
CHMP1A
LDLR
RAPGEF6
WDR61
FGFR1
MLLT3
none
1
10
100
enrichment over random
B
MTOR inh.
AZD-8055, Everolimus, Rapamycin +2
PI3K/AKT inh.
Alpelisib
RICTOR
MLST8
MAPKAP1
VAMP4
RPTOR
MTOR
PDPK1
PIK3CA
PPP2CB
CNPY2
RHEB
SEH1L
ITPKC
C
MEK inh.
Binimetinib, Cobimetinib,
TAK-733 +1
Other TK inh.
Erdafitinib, Ferulic acid,
Futibatinib +4
ERBIN
RHOA
ELMO2
RAPGEF6
QRICH1
RAP2A
STUB1
MAP2K1
FGFR1
TSPAN4
RALGDS
DPP7
CEP78
GRN
TMEM39A
ANKS6
GPBP1
D
Other TK inh.
RAS inh.
RAP1A
GRN
TSPAN4
TLNRD1
EFNA4
STUB1
LSR
KRAS
POLDIP2
NTRK1
FGFR1
SNX2
TMEM39A
DPP7
ILVBL
FGFR1: 675 pairs, 8x expected
best prior: GO BP 52.0x
androgen rec. antag.
DNA repair inh.
FOXN4
NKX6-1
ZNF776
MESP1
PRMT2
ZNF274
AR
EPC1
FHL3
RNF4
TOP2A
ZNF311
ALKBH5
SMC4
POLR3H
GTF2F1
MED11
RNF4: 360 pairs, 19x expected, one of several steps
best prior: DepMap 2.0x
DNA repair inh.
glucocorticoid agon.
ZNF311
NFATC4
EPC1
FOXJ1
TOP2A
KLF12
SMC4
FOXQ1
NR3C1
AGFG1
KLF1
PRMT2
ZNF776
PLAGL2
MESP1
BIRC2
FOXQ1: 340 pairs, 382x expected, one of several steps
best prior: DepMap 0.3x
MTOR inh.
unannotated
STRN4
EXOC1
EXOC2
EMC4
PPP1R37
MTOR
RPTOR
MOB4
ARPC2
DNM2
SEH1L
ACTR2
NCKIPSD
RHEB
PDPK1
VPS29
MLST8
MAPKAP1
MOB4: 320 pairs, 16x expected
best prior: STRING 0.3x
DNA repair inh.
Microtubule inh.
PDCD2
LEO1
SUPT4H1
WDR61
PAF1
TBCC
TOP1
TUBB2A
PLOD3
SUPT16H
CTR9
SSRP1
RTF1
SUPT6H
CDC73
WDR61: 325 pairs, 8x expected
best prior: GO BP 0.0x
DNA repair inh.
Other MAPK inh.
FOXN4
RORA
ZNF710
ZFHX3
PHF19
ZNF311
PRMT2
POU6F1
MESP1
MBD2
SMC4
TOP2A
MAPK14
ZNF776
EPC1
ZBTB14
DNTTIP1
MESP1: 265 pairs, 66x expected, one of several steps
best prior: STRING 0.1x
MEK inh.
MTOR inh.
ELMO2
QRICH1
SLK
RAP2A
RHOA
MOB4
MTOR
ERBIN
MAP2K1
PPP1R37
RPTOR
RAPGEF6
RHEB
CEP78
SEH1L
TSPAN4
PDPK1
ANKS6
RALGDS
MLST8
GPBP1
RAPGEF6: 288 pairs, 8x expected, one of several steps
DNA repair inh.
JAK/STAT inh.
MBD2
ZNF710
STAT1
ZNF311
MESP1
ZNF799
GATA6
JAK2
SMC4
GCFC2
IRF1
TOP2A
BABAM2
PRMT2
SSBP3
EPC1
ZNF845
ZNF772
GCFC2: 252 pairs, 21x expected, one of several steps

**Figure S9. Prototype bridges between drug classes, related to Figure 4**

(A) Genes that the prototype graph places between the targets of different drugs. A k-nearest-neighbor graph (k = 8, cosine) is built over the 8,964 trained prototypes, and the shortest path between the annotated target sets of every pair of the 156 Tahoe-100M drugs with a target is traced (12,090 pairs). A gene's enrichment is the number of drug pairs whose path runs through it, relative to the number expected when each drug's targets are replaced by size- and popularity-matched random genes (200 permutations). 102 genes carry at least 50 pairs at twice the expected number; shown are the 24 with the largest excess, ordered by fold over expected (log scale; dotted line, expected; all $p$ = 0.005, the permutation floor). Orange, the learned prototypes; gray, the best of seven priors (GO Biological Process, gene-description text, STRING[51,52], DepMap co-essentiality[46,47], L1000[39,40], ProteinTalks[127], cell morphology[114]), each rebuilt into the same graph (genomic-position control not shown); open circles at "none", no prior routes a drug pair through the gene. Nine of the 24 are also bridges in at least one prior's graph (at least twofold), six of them in DepMap, more than in any other single prior. Circled and named in color, the bridges in (B) and (C); a two-colored name (PDPK1) marks one a prior also finds.

(B) PDPK1 is the only intermediate on the shortest path between the targets of the mTOR inhibitors (MTOR) and of the PI3K inhibitor alpelisib (PIK3CA). Large nodes, drug targets (tint, side of the path); small gray nodes, their nearest prototypes; edges, kNN edges, wider for higher cosine. Center marker split orange and gray, found by the prototypes and by DepMap (34-fold). Headers, Tahoe-100M mechanism class and drugs on each side.

(C) RALGDS is the only intermediate between the targets of the MEK inhibitors (MAP2K1) and of the drugs annotated to FGFR1 (Tahoe-100M class "Other TK inhibitor", mostly FGFR inhibitors). No biological prior routes a drug pair through it.

(D) The same neighborhood for the eight other bridges among the ten with the largest excess, one drug-class pair per bridge, with the bridge boxed. Subtitles, drug pairs through the gene and fold over expected, the best prior's fold, and "one of several steps" where the bridge is not the only intermediate.

See **Supplementary Note 1**.

# SI 10

## A

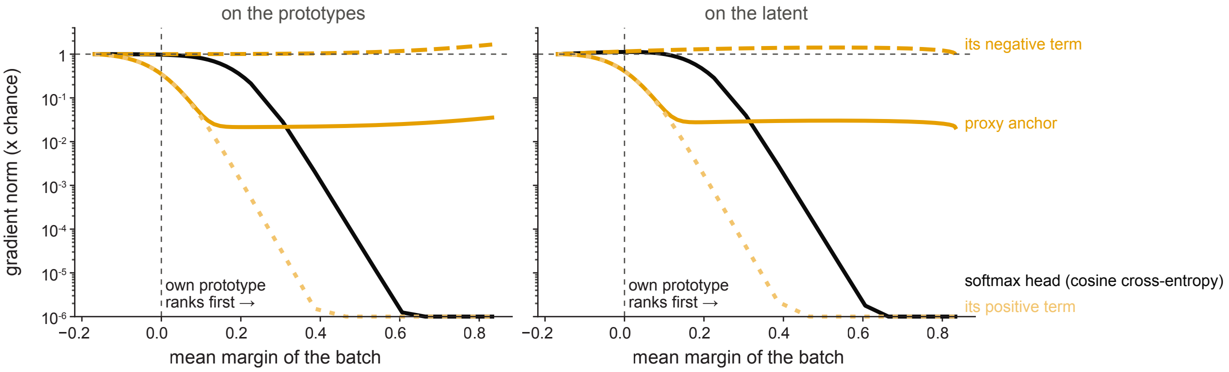

on the prototypes
on the latent
gradient norm (x chance)
1
$10^{-1}$
$10^{-2}$
$10^{-3}$
$10^{-4}$
$10^{-5}$
$10^{-6}$
own prototype
ranks first →
−0.2
0.0
0.2
0.4
0.6
0.8
mean margin of the batch
its negative term
proxy anchor
softmax head (cosine cross-entropy)
its positive term

**Figure S10. Gradient remaining for Proxy Anchor and for a softmax head once the correct prototype wins, related to Figure 1**

Controlled geometry at the shape of the production model: batches of 16,384 cells, a 512-dimensional latent and 8,964 trained prototypes. Each cell's latent is a mixture of its own prototype and random noise, and raising the prototype's share moves the batch from chance (mean margin −0.17) to every cell ranking its own prototype first by a wide margin. Margin, cosine similarity to the cell's own prototype minus that to the most similar other prototype; dotted vertical line, mean margin 0 (about half the cells rank their own prototype first); dotted horizontal line, chance geometry; y axis floored at $10^{-6}$. Lines, gradient norm on the prototypes (left) and on the latent (right) relative to chance geometry, for the softmax head of the cross-entropy models (cosine logits scaled by 30; black) and for Proxy Anchor at its shipped settings ($\alpha = 32$, $\delta = 0.1$; orange), with its two terms: the positive term, which pulls each cell toward its own prototype (dotted), and the negative term, which pushes it away from the other prototypes (dashed). The softmax head keeps its gradient while cells barely win (53% on the prototypes and 68% on the latent when every cell first ranks its own prototype first, margin 0.16) but loses six orders of magnitude by a margin of 0.6, because its normalization includes the correct class; Proxy Anchor's positive term saturates the same way. The negative term normalizes over the other prototypes only and keeps at least its full gradient at every margin, so once its positive term has saturated (margin $\gtrsim 0.15$) Proxy Anchor retains 2–4% of its starting gradient at every larger margin. Performance of the two objectives: **Table S3A** and **S3D**.

## SI 11

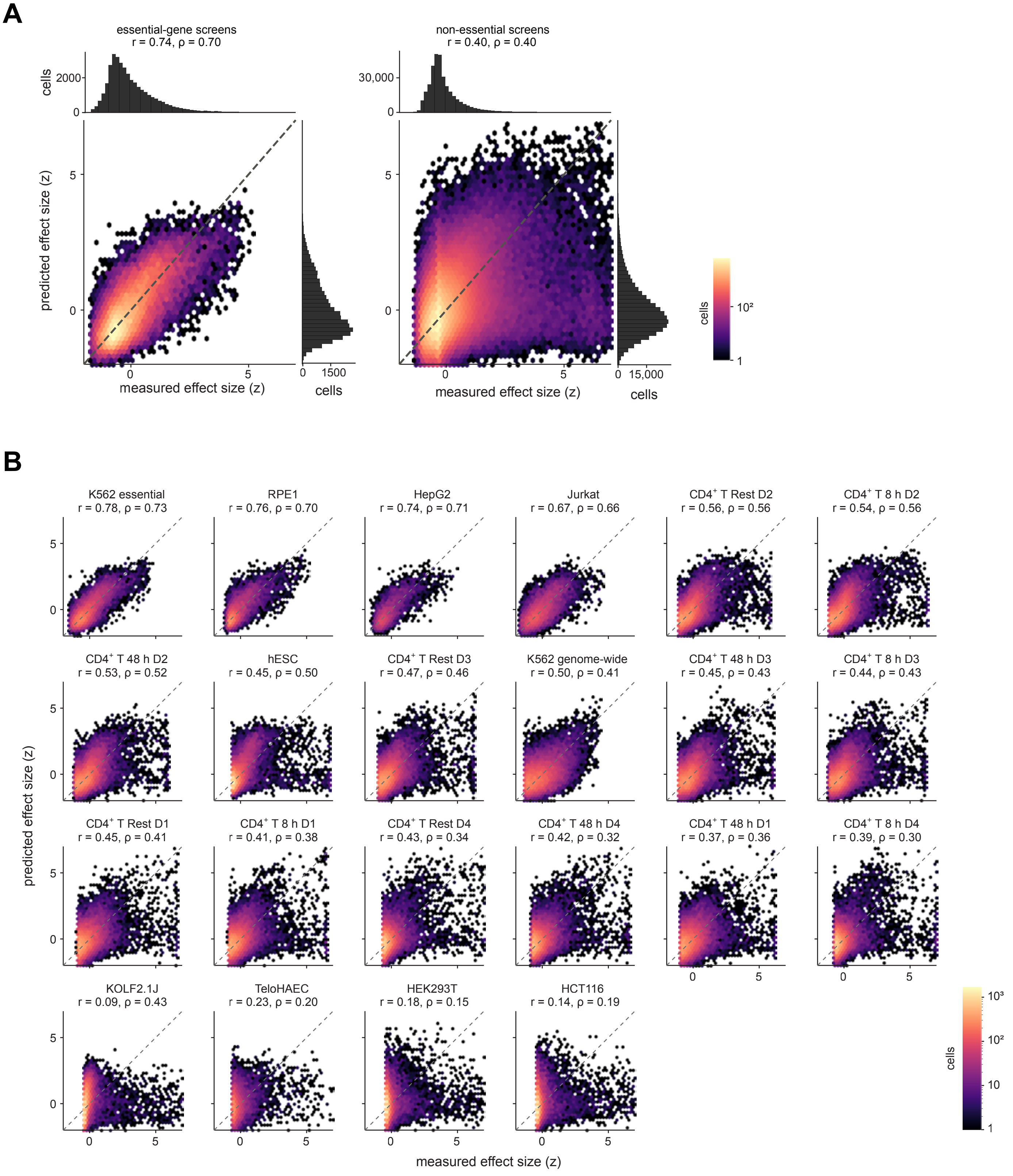

A
essential-gene screens
r = 0.74, ρ = 0.70
non-essential screens
r = 0.40, ρ = 0.40
cells
2000
30,000
predicted effect size (z)
measured effect size (z)
1500
15,000
10²
B
K562 essential
r = 0.78, ρ = 0.73
RPE1
r = 0.76, ρ = 0.70
HepG2
r = 0.74, ρ = 0.71
Jurkat
r = 0.67, ρ = 0.66
CD4⁺ T Rest D2
r = 0.56, ρ = 0.56
CD4⁺ T 8 h D2
r = 0.54, ρ = 0.56
CD4⁺ T 48 h D2
r = 0.53, ρ = 0.52
hESC
r = 0.45, ρ = 0.50
CD4⁺ T Rest D3
r = 0.47, ρ = 0.46
K562 genome-wide
r = 0.50, ρ = 0.41
CD4⁺ T 48 h D3
r = 0.45, ρ = 0.43
CD4⁺ T 8 h D3
r = 0.44, ρ = 0.43
CD4⁺ T Rest D1
r = 0.45, ρ = 0.41
CD4⁺ T 8 h D1
r = 0.41, ρ = 0.38
CD4⁺ T Rest D4
r = 0.43, ρ = 0.34
CD4⁺ T 48 h D4
r = 0.42, ρ = 0.32
CD4⁺ T 48 h D1
r = 0.37, ρ = 0.36
CD4⁺ T 8 h D4
r = 0.39, ρ = 0.30
KOLF2.1J
r = 0.09, ρ = 0.43
TeloHAEC
r = 0.23, ρ = 0.20
HEK293T
r = 0.18, ρ = 0.15
HCT116
r = 0.14, ρ = 0.19
10³
10

**Figure S11. Per-cell effect size predicted by the model, related to Figure 1**

Measured against predicted knockdown effect size for held-out perturbed cells of the production model (cells not used in training; up to 20,000 per dataset, 383,263 in total). Measured effect size is the absolute Mixscale score: the cell's expression projected onto its own perturbation's differential-expression direction, relative to control cells; a negative score is a shift against that direction and counts as an effect of the same size. Predicted effect size is the model's effect-size readout: the effect-size head trained with the model, combined linearly with how confidently the cell matches a perturbation (its largest cosine similarity to any perturbation prototype, the gap to the second largest, the share of the best match among the ten most similar prototypes, and the distance of its control-centered latent from the controls). The five weights were fitted on 32,677 training cells, separate from the held-out cells shown here. Both measured and predicted effect size are z-scored within each dataset. Hexagons, number of cells (log scale); dashed line, identity; r, Pearson correlation; ρ, Spearman correlation.

(A) Cells pooled from the four essential-gene screens (K562 essential, RPE1, HepG2 and Jurkat; left; 34,916 cells; r = 0.74, ρ = 0.70) and from the other 18 screens (non-essential; right; 348,347 cells; r = 0.40, ρ = 0.40), each z-scored within its dataset first. Histograms, cells per bin of measured (top) and predicted (right) effect size. Measured effect size is right-skewed, with a long tail of strongly affected cells; the prediction is closer to symmetric and does not reach that tail.

(B) Each dataset, ordered by the mean of r and ρ. The readout tracks effect size best in the essential-gene screens (r = 0.67–0.78, ρ = 0.66–0.73) and in the $CD4^+$ T cell screens (r = 0.37–0.56, ρ = 0.30–0.56), and least in screens where few cells respond (12–20% of cells with an absolute Mixscale score above 2; TeloHAEC r = 0.23, ρ = 0.20; HCT116 r = 0.14, ρ = 0.19; HEK293T r = 0.18, ρ = 0.15). In KOLF2.1J a few extreme scores depress r (0.09) while the ranking holds (ρ = 0.43). Mean over the 22 datasets, r = 0.46, ρ = 0.44.

The effect-size head alone reaches mean r = 0.40, ρ = 0.40 over the 22 datasets (pooled essential-gene screens r = 0.68, ρ = 0.65; pooled non-essential screens r = 0.36, ρ = 0.35).

## SI 12

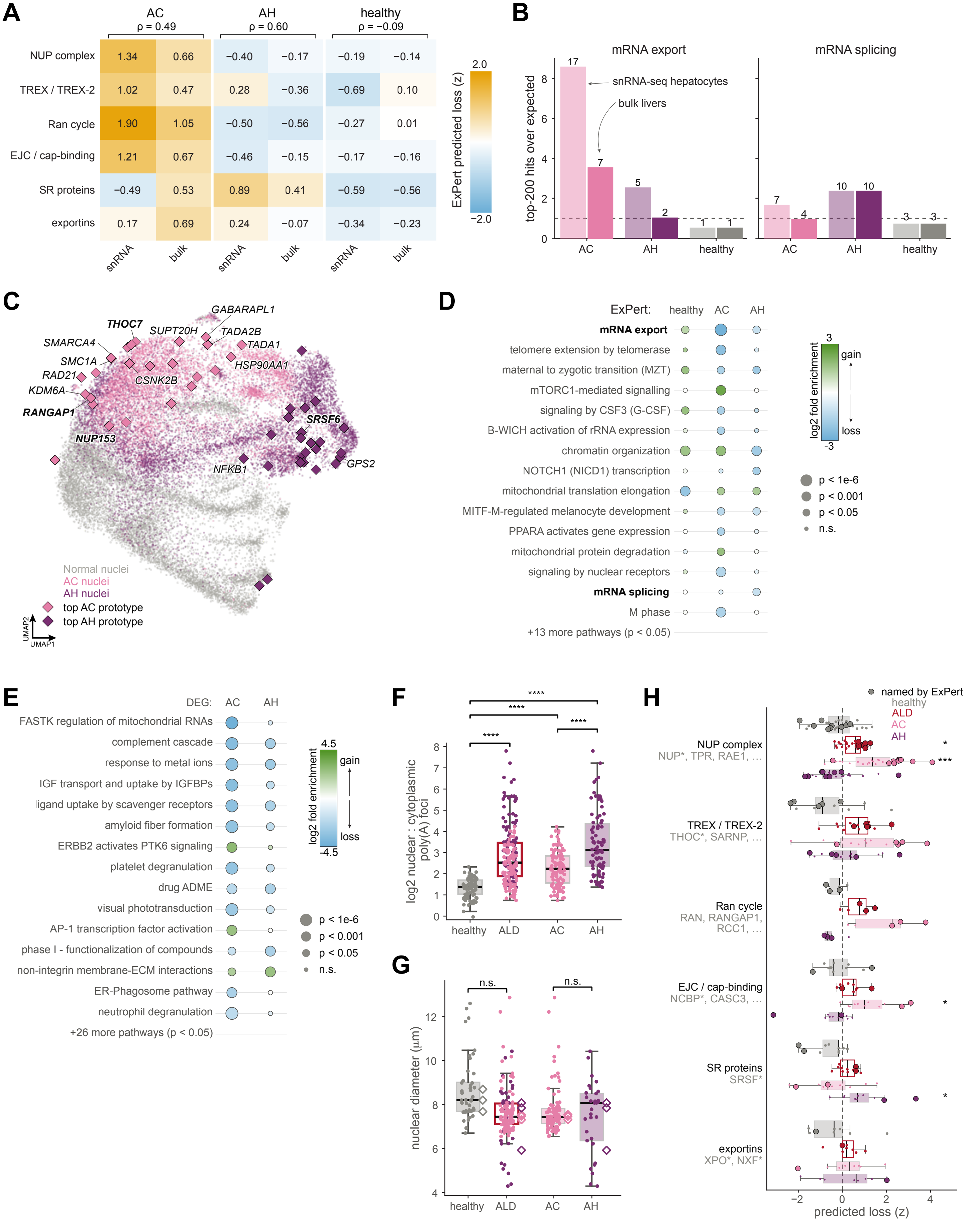

A
AC ρ = 0.49
AH ρ = 0.60
healthy ρ = −0.09
NUP complex 1.34 0.66 −0.40 −0.17 −0.19 −0.14
TREX / TREX-2 1.02 0.47 0.28 −0.36 −0.69 0.10
Ran cycle 1.90 1.05 −0.50 −0.56 −0.27 0.01
EJC / cap-binding 1.21 0.67 −0.46 −0.15 −0.17 −0.16
SR proteins −0.49 0.53 0.89 0.41 −0.59 −0.56
exportins 0.17 0.69 0.24 −0.07 −0.34 −0.23
snRNA bulk snRNA bulk snRNA bulk
ExPert predicted loss (z)
2.0
−2.0
B
mRNA export
mRNA splicing
top-200 hits over expected
snRNA-seq hepatocytes
bulk livers
AC
AH
healthy
C
THOC7
GABARAPL1
SUPT20H
TADA2B
SMARCA4
TADA1
SMC1A
HSP90AA1
RAD21
CSNK2B
KDM6A
RANGAP1
SRSF6
NUP153
NFKB1
GPS2
Normal nuclei
AC nuclei
AH nuclei
top AC prototype
top AH prototype
UMAP2
UMAP1
D
ExPert: healthy AC AH
mRNA export
telomere extension by telomerase
maternal to zygotic transition (MZT)
mTORC1-mediated signalling
signaling by CSF3 (G-CSF)
B-WICH activation of rRNA expression
chromatin organization
NOTCH1 (NICD1) transcription
mitochondrial translation elongation
MITF-M-regulated melanocyte development
PPARA activates gene expression
mitochondrial protein degradation
signaling by nuclear receptors
mRNA splicing
M phase
+13 more pathways (p < 0.05)
log2 fold enrichment
3 gain
loss -3
p < 1e-6
p < 0.001
p < 0.05
n.s.
E
DEG: AC AH
FASTK regulation of mitochondrial RNAs
complement cascade
response to metal ions
IGF transport and uptake by IGFBPs
ligand uptake by scavenger receptors
amyloid fiber formation
ERBB2 activates PTK6 signaling
platelet degranulation
drug ADME
visual phototransduction
AP-1 transcription factor activation
phase I - functionalization of compounds
non-integrin membrane-ECM interactions
ER-Phagosome pathway
neutrophil degranulation
+26 more pathways (p < 0.05)
log2 fold enrichment
4.5 gain
loss -4.5
p < 1e-6
p < 0.001
p < 0.05
n.s.
F
****
log2 nuclear : cytoplasmic poly(A) foci
healthy ALD AC AH
G
n.s.
nuclear diameter (µm)
healthy ALD AC AH
H
named by ExPert
healthy
ALD
AC
AH
NUP complex
NUP*, TPR, RAE1, …
TREX / TREX-2
THOC*, SARNP, …
Ran cycle
RAN, RANGAP1, RCC1, …
EJC / cap-binding
NCBP*, CASC3, …
SR proteins
SRSF*
exportins
XPO*, NXF*
*
***
predicted loss (z)

**Figure S12. ExPert scores of the bulk RNA-seq livers reproduce the snRNA-seq hits, and Figure 5 split by condition (AC and AH), related to Figure 5**

(A, B) ExPert scores of bulk RNA-seq livers. Bulk RNA-seq of 16 livers (4 healthy, 6 AC, 6 AH) from the same biobank as the snRNA-seq donors (donor overlap unknown, so the two cohorts are not independent). Each liver was encoded as one pseudobulk with the healthy livers as the input baseline, the latents were centered over all livers and averaged per condition, and each condition was scored against the 8,964 trained prototypes as in **Figure 5**.

(A) Mean predicted-loss z of the mRNA export families (as in **Figure 5L**) in snRNA-seq hepatocytes and in the bulk livers, per condition. Color and values in each cell, mean z (orange, predicted loss; blue, predicted gain); z is not relative to healthy, so healthy columns are nonzero; brackets, Spearman correlation between the snRNA-seq and bulk columns over the six families. Over the 77 individual export genes the correlation is −0.03 (AC; $p$ = 0.83), 0.46 (AH; $p = 2 \times 10^{-5}$) and 0.37 (healthy; $p$ = 0.0008).

(B) Members of mRNA export and of mRNA splicing among the 200 strongest predicted losses, relative to the number expected by chance (hypergeometric test, background the 8,964 trained prototypes), in snRNA-seq hepatocytes (light) and the bulk livers (dark), per condition. Bars colored by condition (pink AC, plum AH, gray healthy); dotted line, as expected by chance. Numbers, hits; bold, $p < 0.05$. In AC, mRNA export is enriched in both (17 and 7 hits against 2.0 expected; $p = 8 \times 10^{-12}$ and 0.004); in AH, mRNA splicing is enriched in both (10 hits each against 4.3 expected; $p$ = 0.01). Neither set is enriched in the healthy livers.

Across all 8,964 prototypes, bulk and snRNA-seq scores of the same condition correlate weakly or not at all (Spearman ρ −0.03 AC, 0.18 AH, 0.24 healthy), and their top-200 lists share 12, 18 and 11 genes (p = 0.002, $5 \times 10^{-7}$, 0.005); the agreement is at the level of pathways and families rather than individual genes.

(C–H) **Figure 5** split by condition. Each panel repeats a **Figure 5** panel with AC (pink) and AH (plum) shown separately instead of pooled as ALD; rows, tests and scales are those of the main panel.

(C) As **Figure 5B**: UMAP of the 26,807 nuclei in the ExPert latent space, colored healthy (gray), AC and AH, with the 25 highest-scoring prototypes for AC and for AH hepatocytes in the condition's color. Diamonds, prototypes; dots, nuclei. Labeled prototypes belong to mRNA export or splicing (bold) or to the literature-curated pathway rows of **Figure 5E** (chromatin modification, SUMOylation, macroautophagy; italic).

(D) As **Figure 5E**: the same 15 pathways, enriched among the top 200 ExPert predictions of healthy, AC and AH hepatocytes separately (one-sided hypergeometric test; background, all perturbations in the ExPert bank). Rows and the count of further pathways are those selected on the pooled ALD column in **Figure 5E**. Among predicted losses, mRNA export is enriched mainly in AC (17 hits against 2.0 expected, $p = 8 \times 10^{-12}$; AH, 5 hits, $p$ = 0.05) and mRNA splicing in AH (10 hits against 4.3 expected, $p$ = 0.01; AC, $p$ = 0.14).

(E) As **Figure 5F**: the same 15 pathways among the top 200 downregulated and upregulated DE genes of AC and AH hepatocytes separately.

(F) As **Figure 5I**: nuclear-to-cytoplasmic ratio of poly(A) foci (log2 scale) per imaging frame (n = 84 healthy, 103 AC and 90 AH frames from three livers per condition; ALD, AC and AH pooled, open box). Frames could not be assigned to individual livers and are treated as independent. Two-sided Mann-Whitney U test, Benjamini-Hochberg adjusted over the four comparisons shown: healthy versus ALD, AC and AH, all $q < 10^{-13}$; AC versus AH, $q = 8 \times 10^{-9}$ (median ratio 2.6 healthy, 4.7 AC, 8.7 AH).

(G) As **Figure 5J**: diameter of DAPI-segmented hepatocyte nuclei (n = 44 healthy, 93 AC and 33 AH nuclei from 3 livers per condition; median 8.2, 7.4 and 8.1 µm). Open diamonds, per-liver medians, the units tested. Two-sided Mann-Whitney U test on per-liver medians: healthy versus ALD (n = 3 and 6 livers), $p = 0.10$; AC versus AH (3 and 3), $p = 0.70$.

(H) As **Figure 5L**: ExPert predicted loss-of-function scores (z) of the mRNA export machinery families in healthy, ALD (AC and AH pooled; open box), AC and AH hepatocytes. Points, family members; large outlined points, members among the top 200 predicted losses of AC or of AH. Paired two-sided Wilcoxon test over all family members versus healthy, Benjamini-Hochberg adjusted over families and the three disease groups: in ALD, NUP complex (q = 0.05); in AC, NUP complex ($q = 1 \times 10^{-4}$) and EJC / cap-binding (q = 0.04); in AH, SR proteins (q = 0.04).

# SI 13

A

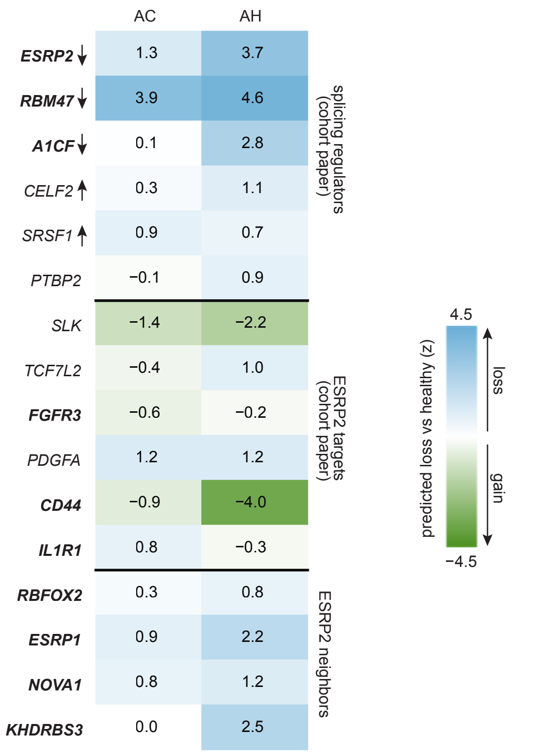
AC
AH
ESRP2 1.3 3.7
RBM47 3.9 4.6
A1CF 0.1 2.8
CELF2 0.3 1.1
SRSF1 0.9 0.7
PTBP2 −0.1 0.9
SLK −1.4 −2.2
TCF7L2 −0.4 1.0
FGFR3 −0.6 −0.2
PDGFA 1.2 1.2
CD44 −0.9 −4.0
IL1R1 0.8 −0.3
RBFOX2 0.3 0.8
ESRP1 0.9 2.2
NOVA1 0.8 1.2
KHDRBS3 0.0 2.5
splicing regulators (cohort paper)
ESRP2 targets (cohort paper)
ESRP2 neighbors
4.5
−4.5
predicted loss vs healthy (z)
loss
gain

B

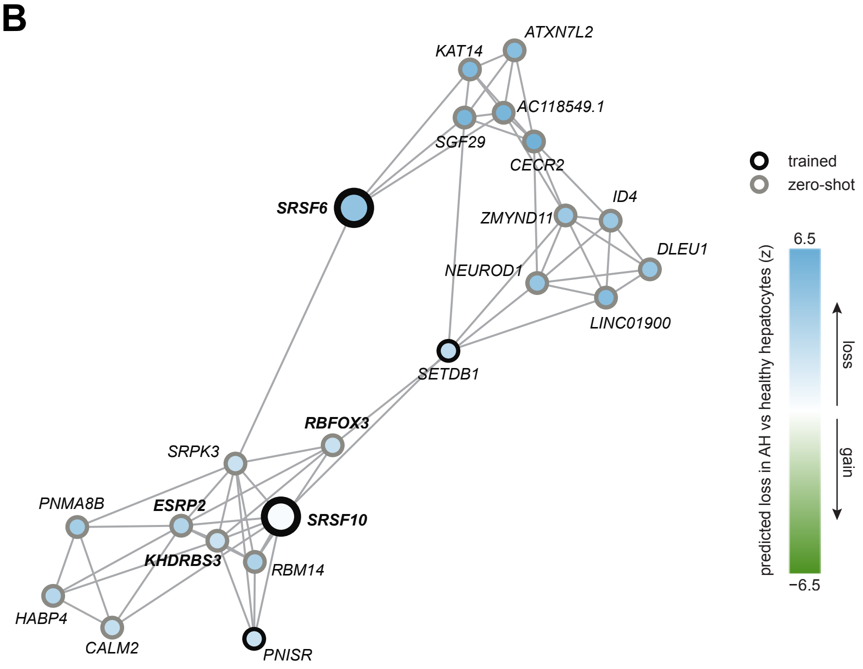
KAT14
ATXN7L2
AC118549.1
SGF29
CECR2
ID4
ZMYND11
DLEU1
NEUROD1
LINC01900
SETDB1
SRSF6
RBFOX3
SRPK3
PNMA8B
ESRP2
SRSF10
KHDRBS3
RBM14
HABP4
CALM2
PNISR
trained
zero-shot
6.5
−6.5
predicted loss in AH vs healthy hepatocytes (z)
loss
gain

**Figure S13. Zero-shot predictions in alcohol-associated liver disease for the splicing regulators and ESRP2 targets implicated in this cohort, related to Figure 5**

**Figure 5** scores the 8,964 trained prototypes. Here the same hepatocyte profiles (condition latents centered per cell type, as in **Figure 5**) are scored against the full vocabulary of 19,895 rows, which adds the 10,931 zero-shot prototypes placed from prior knowledge (**Figure S6**). Exact-gene zero-shot recovery is about 26-fold over chance in contexts seen in training and 14-fold in held-out contexts (**Figure 3B**), weaker evidence than a trained prototype. Score, the cosine similarity between the condition's centered hepatocyte latent and the prototype, as in **Figure 5** (higher, the state resembles loss of the gene), shown as the change from healthy hepatocytes in units of the standard deviation of the healthy scores over all 19,895 rows (z; healthy = 0); color scale as **Figure 5E**: blue, predicted loss relative to healthy; green, predicted gain.

(A) Change from healthy (z) in AC and AH hepatocytes of the genes implicated by the study of this cohort[73]: the splicing regulators it reports as changed in severe alcoholic hepatitis (arrow, the reported direction of their expression: ESRP2, RBM47 and A1CF down; CELF2 and SRSF1 up; PTBP2 cell-type dependent), the ESRP2 targets it finds mis-spliced (SLK, TCF7L2, FGFR3, PDGFA, CD44 and IL1R1), and the four prototypes nearest to ESRP2 (RBFOX2, ESRP1, NOVA1 and KHDRBS3). Bold, zero-shot prototypes. The three regulators reported down are all predicted losses in AH (ESRP2 3.7, RBM47 4.6, A1CF 2.8; in AC 1.3, 3.9 and 0.1), as are ESRP1 (2.2) and KHDRBS3 (2.5). The ESRP2 targets do not shift consistently: SLK and CD44 shift toward predicted gain in AH (−2.2 and −4.0).

(B) The neighbors with the strongest predicted loss around two hubs: SRSF6, the strongest predicted splicing loss in AH hepatocytes (z = 5.1 relative to healthy; rank 25 of 19,895), and SRSF10, the trained SR protein nearest to the ESRP2 zero-shot prototype. For each hub, the 10 prototypes with the strongest predicted loss in AH relative to healthy among its 50 nearest, joined into one graph with edges to each gene's four nearest within the set (width, cosine). Fill, change from healthy (z) in AH hepatocytes; border, trained (black) or zero-shot (gray); large nodes, the two hubs (SRSF10 itself is near healthy, z = 0.3); bold labels, the hubs and the regulators of (A). Around SRSF10, ESRP2 links most strongly to KHDRBS3, SRPK3 and RBM14, and to RBFOX3; the trained SETDB1 joins the two groups. The neighbors of SRSF6 are zero-shot chromatin regulators (SAGA/ATAC components SGF29, KAT14 and ATXN7L2; CECR2; ZMYND11), NEUROD1, ID4 and long noncoding RNAs.

Zero-shot prototypes are the shipped graph fill ($\cos^4$ weights, k = 25); with the earlier cosine-weighted fill the results are similar.

SI 14

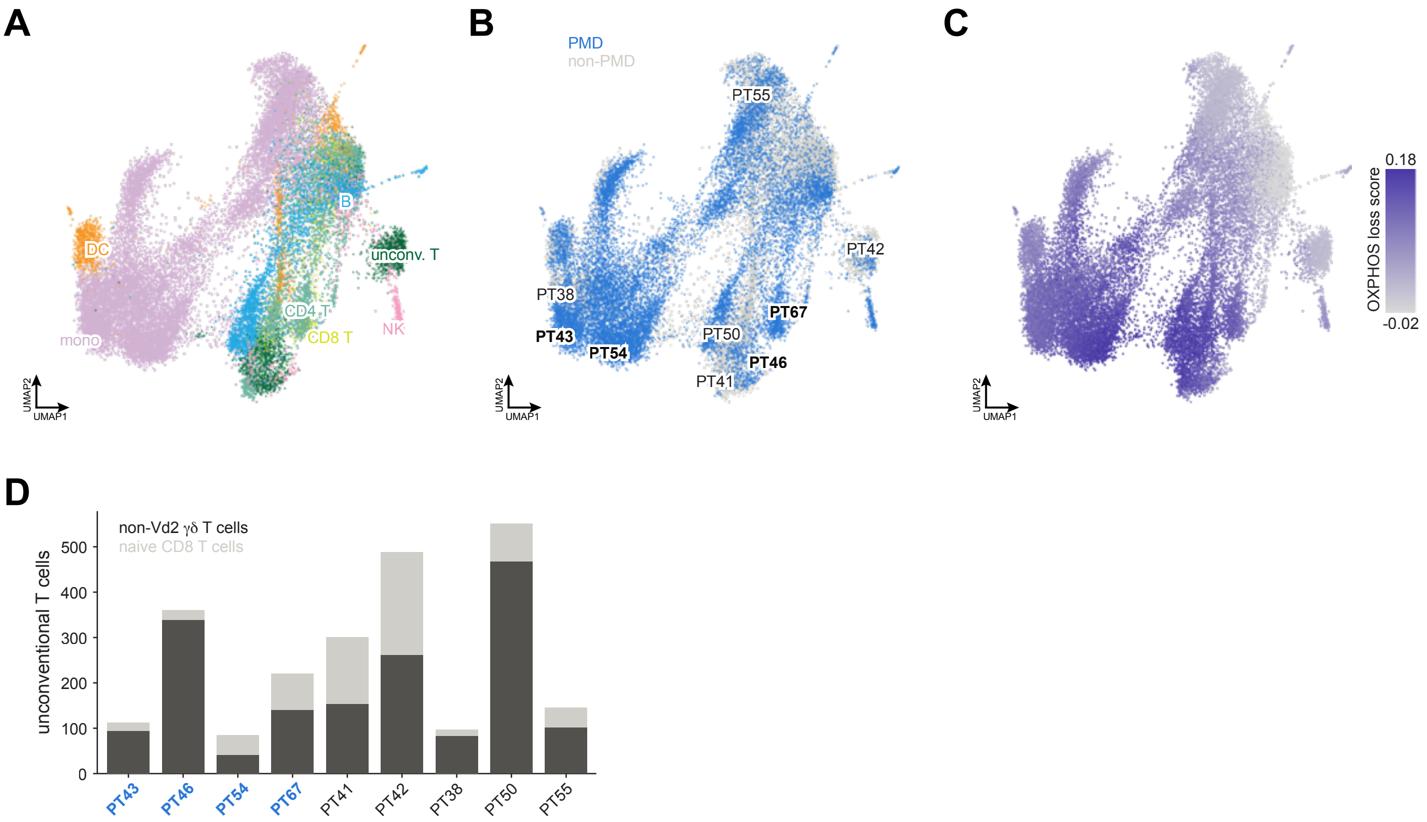
A
DC
mono
B
unconv. T
CD4 T
CD8 T
NK
UMAP2
UMAP1
B
PMD
non-PMD
PT55
PT42
PT38
PT43
PT54
PT50
PT67
PT46
PT41
UMAP2
UMAP1
C
0.18
OXPHOS loss score
-0.02
UMAP2
UMAP1
D
non-Vd2 γδ T cells
naive CD8 T cells
unconventional T cells
0
100
200
300
400
500
PT43
PT46
PT54
PT67
PT41
PT42
PT38
PT50
PT55

**Figure S14. PBMC overview: all cell types, donors and the OXPHOS-loss score, related to Figure 6**

All 28,513 ex vivo peripheral blood mononuclear cells (PBMCs) from nine donors, encoded by ExPert as in **Figure 6** (MT-masked input, latent centered per cell type); UMAP of the ExPert latent. Donors: PMD, primary mitochondrial disease (PT43, PT46, PT54, PT67); non-PMD (PT41, PT42); undiagnosed (PT38, PT50, PT55). Cell types are the SingleR main labels.

(A) Cells colored by cell type (Mono, monocytes; DC, dendritic cells; unconv. T, unconventional T cells).

(B) Cells colored by donor group (blue, PMD; light gray, non-PMD; gray, undiagnosed); each donor is named at its densest region (PMD in bold).

(C) Cells colored by the uncentered OXPHOS-loss score (cosine of the cell's latent with the OXPHOS knockout direction, smoothed over its nearest neighbors in the latent), as in **Figure 6C**; color scale clipped to the 2nd–98th percentile.

(D) Composition of the unconventional T cells (SingleR main label "Nonclassical T cells") by the SingleR fine label, per donor (stacked bars; PMD donors in bold): in total 1,683 non-Vδ2 γδ T cells and 677 naive CD8 T cells.

### SUPPLEMENTAL INFORMATION

#### Supplementary Notes

**Supplementary Note 1. Prior-graph prototype augmentation Rationale.** During training, ExPert can pull the prototypes of perturbations it has seen least toward the positions their prior-knowledge neighbors predict. The pull blends a prototype toward the similarity-weighted mean of its nearest neighbors in the prior graph. We tested whether this helps the trained perturbations themselves, and not only genes absent from training. Models were trained with the K562 screens held out and evaluated on 10,061 held-out K562 queries of trained perturbations.

**Gated pull.** Pulling every trained prototype raised top-10 hits by 2% ($p = 0.042$; **Figure S2B**). The gain concentrated where training support was weakest: perturbations in the lower half of training support gained 4% more hits, while those in the upper half did not change (**Figures S2C** and **S2D**). ExPert therefore restricts the pull to the 1,004 prototypes with the lowest support.

**Train-time adjustment.** The pull had to be applied during training. Moving these prototypes by the same amount in a finished model raised hits on their own queries by 44% but cost 10% elsewhere, a net loss of 2%. Training absorbed those losses instead, for a net gain of 3% (**Figure S2E**). Pull strength mattered as well. Light post-hoc adjustments gained up to 2%, but stronger ones lost steadily more, down to −23% when prototypes were moved all the way to their prior positions (**Figure S2E**).

**Supplementary Note 2. Selection of the input gene panel**

**Statistic.** Separability requires within-perturbation variance, since with one profile per perturbation every perturbation is trivially its own nearest centroid. Each perturbation's cells were therefore split into two pseudobulk replicates and genes scored by the ratio of between- to within-perturbation variance (**Methods**). The ratio was used unmoderated, since adding a variance prior, the usual guard against near-zero within-group variance, lowered held-out separability monotonically (0.234 to 0.124 at the strongest prior tested). Per-gene scaling and control subtraction cannot change the ranking, because scaling multiplies numerator and denominator alike and subtracting a constant profile shifts every group equally, and empirically left the selected genes unchanged.

**Criterion and construction.** At 500 genes per dataset, unioned, the F-statistic panel reached 0.419 held-out separability, against 0.333 for highly variable genes (seurat_v3 union of per-dataset top 500; 0.356 at their best size of 1,500) and 0.131 for a size-matched random panel. Genes measured in every dataset scored 0.13–0.20 at all sizes (**Figure S1F**). The gain held in both study families, Marson2025, which contributes 12 of the 22 datasets (0.642 against 0.503), and the rest (0.151 against 0.129). Giving each dataset only its own top 500 scored higher (0.495) but is not deployable, because a query's dataset of origin is unknown. Separability was flat between 250 and 500 genes per dataset (0.426 and 0.419) and declined above.

**Expression level.** The statistic correlates with expression (Spearman 0.745; selected genes sit at the 93rd expression percentile), which follows from the mean–variance relationship rather than from normalization. Removing the trend collapses the correlation to −0.040 but costs 0.038 separability on average, mostly in the cell-line datasets, so it was kept. The two criteria are not nested. A union of 3,000 variable genes per dataset (16,094 genes) still misses 10.4% of the panel, mostly highly expressed genes with low relative variance such as B2M, LDHB, PFN1 and ENO1.

**Target genes.** Recomputing the statistic with each perturbation's own target masked, so that a gene cannot be selected for being knocked down rather than for reporting responses, changed 6% of the panel (Jaccard 0.94).

**Panel composition.** The union of 500 genes per dataset gives 3,096 genes, 71.9% fewer than the sum of the per-dataset selections; 42.1% are selected by a single dataset and the median gene by two. This panel was chosen because masking has no inference-time counterpart, since the mask depends on the dataset of origin and dropping it would feed the encoder genes that were zero throughout training.

**Benchmark impact.** The panel was selected once over all 22 datasets, so a fold's held-out datasets contribute to the axis on which they are evaluated. Rescoring each of the 34 fold-and-held-out-dataset pairs under a panel rebuilt from that fold's training datasets alone, the shipped panel was higher by a mean of 0.012 top-1 (3.2% relative; maximum 0.056, 17.1%) in 29 of 34 pairs. The effect is bounded because held-out datasets keep 84.8% of their own top-500 genes on average even when excluded, most being selected independently by other datasets. It concentrates in the condition folds, where the resting CD4$^+$ T-cell fold removes four contributing datasets at once and shows the largest gain (0.556 to 0.601), while donor and cell-line folds range from −0.0005 to +0.0053 and three cell-line folds sit near the floor under any gene set (top-1 0.009–0.017). Because every method uses the same panel, relative comparisons are unaffected, though a per-fold panel is the strictly correct design where absolute condition-fold accuracy matters. Datasets outside the 22, including the drug screens and disease cohorts of **Figures 4-6**, are unaffected.

**Supplementary Note 3. Placement of zero-shot prototypes**

**Rationale.** Of the 19,895 genes in ExPert's vocabulary, 10,931 have no training cells, so they have no learned prototype. ExPert places each of them from prior knowledge, using the prototypes of trained genes that the priors consider similar. These zero-shot prototypes serve two purposes: they let ExPert name untrained genes, and during training they act as negatives, which teaches the model to keep trained perturbations away from plausible but wrong candidates. We varied how the placement map is built, how many neighbors it uses and how they are weighted, and scored each choice on the 12 frozen fold models (**Figure 3B**; **Table S5**).

**Map**. A graph fill places each untrained gene at the similarity-weighted mean of the prototypes of its 25 nearest trained neighbors in each prior, averaged over the text, DepMap[46,47] and STRING[51] priors. It recovered unseen perturbations as well as a linear map fitted from priors to prototypes (10.5- against 10.4-fold over chance) and better than a multilayer perceptron (8.0-fold), and it needs no fitting (**Figure S6D**). The best of the 62 other combinations of prior sources reached 10.8-fold with the graph map, so the choice of priors matters little once the three are combined.

**Neighbor weighting**. Weighting each neighbor by its prior similarity alone hardly distinguishes them: within a gene's 25 nearest neighbors the similarities are so close that the weighted mean acts as a near-uniform average (24.5 effective neighbors of 25), which pulls every fill toward the center of the space. Raising the similarity to the fourth power lets the closest neighbors dominate. This raised zero-shot top-10 recall by 8.8% in contexts seen in training (12 of 12 fold models improved; $p$ = 0.0005) and by 15.9% in held-out contexts (10 of 12; $p$ = 0.04), while trained perturbations lost 0.4% (**Figure S6C**). Sharper weightings gained less (eighth power, +8.1% and +13.1%; softmax at temperature 0.05, +2.7% and +7.8%), and weighting neighbors by how well supported or how well recovered they were in training lost up to 79% (tested on the fold-4 model only).

**Number of neighbors**. With fourth-power weights, 25 neighbors gave the best net change. Using 50 or 100 neighbors helped unseen perturbations in training contexts more (+12.9% and +12.0%) but helped them less in held-out contexts (+5.0% and +8.7%) and cost trained perturbations more (−0.9%

and −1.2%) (**Figure S6B**). How the neighbors were weighted therefore mattered more than how many were used.

**Geometry**. Fills sit closer to the center of the trained prototypes than trained prototypes do (median angle to their mean direction 85.8° against 89.0°). Trained genes placed by the same map, as if they were unseen, shift as far or further (83.6°), so the shift comes from averaging over neighbors rather than from inaccurate priors (**Figure S6A**). It bounds how finely two untrained genes can be told apart.

**Training with zero-shot prototypes as negatives**. Adding the zero-shot prototypes of the 9,874 untrained genes measured in a screen to the 8,964 trained ones costs held-out queries of trained perturbations top-10 hits, because the fills are plausible distractors. A model filled only after training lost 488 hits on 10,061 held-out K562 queries. Training with the fills as negatives recovered 52% of that loss for ExPert (+252 hits) and 59% for a model pulling every prototype (+290; **Figure S6E**), so the zero-shot prototypes are built into the model during training rather than added afterwards.

**Supplementary Note 4. Prompts for the gene-description text priors**

**Rationale**. The text priors are embeddings of a short description of each gene, written by a language model from records retrieved for that gene (**Methods**). Two choices shape what the embedding captures. The description should use vocabulary that many genes share, such as named pathways, protein families and compartments, so that functionally related genes embed near each other. It should also withhold the gene's identity, its symbol, aliases and full name, so that similarity between two descriptions reflects their biology rather than their names. The instructions below encode both.

**Source records**. For each gene, the records were assembled into one block of text, one section per source in a fixed order, each under a header: NCBI[128] Gene Summary, UniProt Annotations[129], Reactome Pathways[111], STRING[51] Top Interactors, DepMap[46,47] / Open Targets, and Recent PubMed Literature. Sources that returned nothing were left out.

**Request**. Each request consisted of the system prompt below, identical for every gene, and a user message of the form "GENE: ", a blank line, and "SOURCE DATA:" followed by the source records. Descriptions were limited to 256 words, and responses to 1,500 tokens. Claude Sonnet was sampled at temperature 0.3; Claude Opus, which does not accept a temperature setting, at its default. When a model declined to answer or returned no text, the same request was repeated with progressively fewer sources: without PubMed; without PubMed and DepMap; with UniProt, Reactome and STRING only; with UniProt and Reactome; with NCBI and UniProt; with UniProt alone; and with NCBI alone. The descriptions written by Claude Opus 4.8, used by the production model, were generated by collaborators at NVIDIA with the same instructions and request format (**Table S12)**.

**Embedding**. Each description was embedded as plain text. Encoders run locally were applied to at most 512 tokens, with the last hidden states mean-pooled and L2-normalized; API encoders returned one vector per description, which was L2-normalized. The production model uses the Claude Opus 4.8 descriptions embedded with Qwen3-Embedding-8B (4,096 dimensions).

**Supplementary Note 5. Justifications for recovered drug misses**

**Rationale**. A drug counts as a miss when ExPert does not rank its annotated target in the top ten, but the ranking may still reflect the drug's biology. A drug may activate rather than inhibit its target, act through a secondary target the annotation omits, or be labeled by an activity that is not its main one. Counting these drugs as misses would understate how much drug biology ExPert recovers. We therefore examined the ranked lists of the drugs ExPert missed and compared them with the literature and known drug properties (**Methods; Table S8**). Glucocorticoids act through activation, which ExPert captured. Glucocorticoids activate their receptor, NR3C1, rather than inhibiting it, so their effect is missed when only the loss-of-function ranking is considered. Consistent with this, NR3C1 sat near the bottom of the loss-of-function ranking for every glucocorticoid (ranks 7,799–8,845 of 8,964), as expected for an activator. Instead, their rankings were led by FKBP5, a canonical target induced by the activated receptor, which ranked #1-#3 in the gain-of-function direction for triamcinolone, betamethasone dipropionate, dexamethasone and budesonide. These four drugs account for 4 of the 13 explained misses. Annotations that name a secondary activity or omit known properties. ExPert captures the secondary activity and misannotations of the drugs within the dataset. Tahoe-100M annotates clobetasol propionate by its CYP3A4/5 inhibition, but the drug is a potent glucocorticoid. ExPert placed it with the annotated glucocorticoids (glucocorticoid-class similarity z = 5.32) and ranked FKBP5 first in the gain-of-function direction. FKBP5 is a direct transcriptional target of the activated glucocorticoid receptor and one of the most consistent markers of glucocorticoid signaling. Independently of ExPert, clobetasol propionate induced FKBP5 expression more strongly than any other drug in the screen. Belinostat is annotated as HDAC6-selective but is a pan-HDAC inhibitor. ExPert ranked the SIN3–HDAC1/2 corepressor complex at the top of its list. Abemaciclib is annotated as a CDK4/6 inhibitor, but it also inhibits GSK3α and GSK3β. ExPert ranked GSK3A #2 and GSK3B #13, while palbociclib, a CDK4/6 inhibitor without this activity, ranked them #1,575 and #8,119. Methylprednisolone succinate is annotated as a glucocorticoid receptor agonist, but it is a prodrug that must be activated before it binds the receptor. ExPert detected no glucocorticoid response at any dose (class similarity z = 0.63; FKBP5 rank 4,477), and neither did plain expression, as expected for the inactive ester. Responses that match other members of the target pathway. BI-3406 inhibits SOS1, which activates RAS upstream of MEK, so it reduces the same MAPK signaling that MEK inhibitors block. Consistent with this, ExPert placed it with the MEK inhibitors (MEK-class similarity z = 3.09; 3.50 in expression) in every cell-line group except those with BRAF V600 mutations, where mutant BRAF activates MEK independently of RAS. Ponatinib, a multi-kinase inhibitor, likewise resembled the MEK inhibitors (z = 3.07; 4.48 in expression), increasingly with dose, consistent with its reported direct inhibition of BRAF, which also acts on the same pathway. Zero-shot recovery of an untrained target. The annotated targets of tofacitinib and tofacitinib citrate, JAK1 and JAK3, have no trained prototype, so they could only be recovered among the zero-shot prototypes placed from gene priors. Ranked against all 19,895 prototypes, the zero-shot JAK1 prototype placed #31-#51 across doses. Among the trained prototypes, ExPert ranked the type I interferon receptor arm that JAK1 belongs to near the top, with TYK2, IRF9 and IFNAR1 in the top five, in agreement with the known suppression of interferon signaling by tofacitinib. ExPert recovers possible sample mixup. The wells labeled elimusertib, an ATR inhibitor, behaved like a MEK inhibitor in both ExPert and expression (MEK-class similarity z = 4.83 and 6.30), at all three doses, with no ATR signal. The MEK inhibitor TAK-733 sat in the neighboring well for all drug doses, so we attribute this to a likely sample mix-up and did not count it among the explained misses.

#### Supplementary Note 6. Prototype bridges between drug classes

**Rationale**. If perturbing genes in one pathway produces related responses, their prototypes should lie close together and the space should contain the routes between drug targets. This analysis uses no drug response data. It uses only the trained prototype table and Tahoe-100M's[64] target annotations, and so asks what training on genetic perturbations has already encoded.

**Graph and paths**. Each of the 8,964 trained prototypes was joined to its 8 nearest neighbors by cosine similarity, the graph symmetrized, and each edge weighted by one minus the cosine. For each of the 12,090 pairs among the 156 Tahoe-100M drugs with at least one annotated target, we traced the shortest path between the two target sets; 11,744 pairs are connected. A gene's traversal count is the number of drug pairs whose path passes through it, excluding path endpoints and either drug's own targets.

**Null**. A frequently traversed gene need not be specific, since a hub lies on many shortest paths whatever the endpoints. Each drug's target set was therefore replaced by a random set of the same size, each gene drawn from the same popularity bin as the target it replaced (20 bins, 200 permutations). A bridge must exceed expectation both absolutely (≥50 drug pairs) and as a ratio (≥2-fold); hubs meet the first criterion and fail the second. 102 genes qualify on both, 89 of them also at $p \leq 0.05$. **Figure S9A** shows the 24 with the largest excess, all at $p = 0.005$, the floor set by 200 permutations.

**Comparison with prior knowledge**. The same graph and paths were rebuilt from seven gene-level priors (GO[109] Biological Process, language-model gene descriptions, STRING[51], DepMap co-essentiality[46,47], L1000[40], ProteinTalks[127] and cell morphology[114]; **Methods**) and from genomic position as a negative control. Most priors connect 11,159–11,664 of the 12,090 drug pairs; ProteinTalks connects 7,321 and genomic position 6,651. Nine of the 24 bridges reach twofold in at least one prior's graph: *MTHFD1, GPBP1, MLST8, PTPN11, PDPK1, RNF4, THAP1, FGFR1* and *MLLT3*. DepMap co-essentiality recovers six, more than any other prior, and no prior recovers all nine. The remaining 15, including the strongest (*FOXQ1*, 382-fold), appear only in the prototypes.

*PDPK1*, between the *mTOR* and *PI3K* inhibitors (**Figure S9B**). *PDPK1* is the only gene on the shortest path between *MTOR*, the target of five mTOR inhibitors, and *PIK3CA*, the target of alpelisib. 581 drug pairs run through it, 26-fold over matched expectation, and 573 of those join drugs of different annotated mechanism. The position matches the biochemistry. *PDPK1* is recruited by the *PIP3* that *PI3K* produces, phosphorylates *AKT* at Thr308 and activates the mTORC1 substrate S6K, placing it between the two along one axis, and its neighborhood on the mTOR side contains *RPTOR, RICTOR, MLST8, MAPKAP1* and *RHEB*. DepMap routes the same pairs through *PDPK1* at 34-fold, with text and STRING weaker (8.6- and 6.6-fold). This bridge is a known connection the prototypes recover independently.

*RALGDS*, between the *MEK* and *FGFR* inhibitors (**Figure S9C**). *RALGDS* is the only gene on the shortest path between *MAP2K1*, the target of five MEK inhibitors, and *FGFR1*. 288 drug pairs run through it, 40-fold over expectation. *RALGDS* binds GTP-bound RAS and activates the Ral GTPases on a branch parallel to RAF–MEK–ERK[161], while *FGFR* signals to *RAS* through *FRS2*, *GRB2* and *SOS* [162], so it sits between a receptor upstream of RAS and a kinase on a different *RAS* effector arm. None of the seven biological priors routes a single drug pair through *RALGDS*, and only the genomic-position control reaches twofold, which we read as chance.

**Interpretation and limits**. A bridge means the response to knocking down that gene lies between the responses to knocking down the two drugs' targets in the learned space. It is a claim about response similarity, not physical interaction or direction of signaling, and it inherits whatever errors the drug-target annotation carries. Prior folds were computed only for the 24 bridges shown, and pair counts include drugs whose mechanism Tahoe leaves unclassified. Source data in **Figure S9**.

**Supplementary Note 7. SOX2 ranks on the gain side in the degron screen**

**Observation**. *SOX2* is one of three degron targets never perturbed in training, so it could be recovered only through its zero-shot prototype, ranked against all 19,895 prototypes. In the *SOX2* clone (521 auxin-treated nuclei), ExPert ranked *SOX2* 236th, against a median of 4,212 for the genes depleted in the other clones scored in the same response, but in the gain direction; in the loss direction it ranked 8,495th (**Figure 4J**). The *SOX2* clone also had the most reproducible response of the screen (split-half correlation 0.50, against 0.17 for the next clone), so the gain-side placement is not noise.

**Transcript level**. Auxin-induced degradation removes the protein but leaves the gene's transcription intact. In the *SOX2* clone, *SOX2* mRNA itself rose after auxin (log2 fold change +0.44 against DMSO; *SOX2* is the best-sampled clone of the screen, with several times more nuclei than any other). A transcript that increases while its protein is degraded resembles a gain of *SOX2* in expression, and a readout of the cell state can place the response on the gain side even though the protein is lost.

**Interpretation**. Two explanations, not mutually exclusive, fit this. The first is a property of the degron system. Tagged proteins can be partly degraded before auxin is added, and cells may compensate by raising the gene's transcription, so auxin removes a protein whose transcript is already elevated and rises further. The second is *SOX2*'s documented autoregulation [71,72]: *SOX2* binds the regulatory elements of its own gene, so removing the protein changes *SOX2* transcription and the transcription of its targets in the same direction as a change in *SOX2* dose. The data here cannot separate the two. A clone-matched comparison of *SOX2* mRNA before auxin, or a degron of SOX2 lacking its autoregulatory binding, would. Either way, the *SOX2* clone shows that ExPert detected a specific, reproducible *SOX2* response, while the sign of the call depends on how the depletion is achieved.

**Table S1. Training corpus**

(A) Store: one row per dataset, with study, DOI, perturbation type, cell line or cell type, donor, condition and context; quality-control-passed perturbed cells; perturbations with cells and with ≥10 cells; median cells per perturbation; percentage of perturbed cells with a measured effect size and their median effect size (Mixscale[120] score).

(B) Training database: one row per dataset, with context; whether the dataset counts toward the selection rule; perturbed and control cells; selected perturbations; median cells per perturbation; training and validation cells of the production split; perturbations pulled toward priors.

(C) Functional classes of the selected perturbations: number and percentage of selected, all measured and not selected perturbations in each class, with odds ratio and Fisher's exact test (selected vs not selected). Classes overlap. Transcription factor, Lambert et al. human transcription factor census[164]; chromatin regulator, signaling component, metabolic enzyme and transcription, membership of the corresponding Reactome top-level pathway; complex subunit, member of a CORUM[112] complex; common essential, gene effect < −0.5 in ≥90% of DepMap[46,47] cell lines.

**Table S2. Ablations of the prototype source, model capacity, invariance and prior pull**

Each part is one stage of model development, scored on a single held-out (out-of-distribution, OOD) set, so values are comparable within a part but not across parts. Each row is one training run, with its baseline, the change relative to the baseline (from the run configurations), the readout that was scored (cosine similarity to the learned prototypes, or the classifier head), top-k recall, and the change in top-10 recall relative to the baseline. Per-cell top-10 recall on training and validation cells, and their ratio, indicate over- or underfitting.

(A) Prototype source (earlier training database, full scale): prototypes from a frozen text prior with a learned adapter, compared with a randomly initialized learnable table and a free classifier.

(B–D) Model capacity (B), invariance to dataset (C), and prior pull (D) (fold 4): scored on the K562 essential and genome-wide screens held out together (10,061 pseudobulk queries of up to 25 cells) against all 8,964 prototypes. w_top10 and w_rankmed weight each query by its knockdown effect size; rank_med, median rank. beyond_resolution marks changes in top-10 recall larger than the paired resolution (±0.0040, 2 SD); carried_forward marks runs that became the baseline of a later stage (in D, the selection path to the final model). In D, every run is compared with the common reference run without the pull (BASE); parent gives the run it was derived from.

**Table S3. Ablations of the training objective**

Columns as in **Table S2**.

(A) Training objective (fold 0; held-out HepG2 cell line and $CD4^+$ T cell donor 2): in-training OOD top-10 recall, each run scored on the readout it trains (classifier head for cross-entropy runs, cosine similarity to the learned prototypes for Proxy Anchor runs).

(B) Prototype initialization (200-perturbation database): prototypes initialized from the text-prior embeddings and fine-tuned with Proxy Anchor, compared with random initialization.

(C) Prototype readout compared with a classifier head (small-scale runs, earlier training database): frozen text prior with a linear adapter and cosine readout, compared with a classifier head trained with cross-entropy; replicated runs (mean of n = 3), runs without the conditional domain-adversarial term, and a donor and cell-line holdout.

(D) Proxy Anchor compared with a classifier head (small-scale runs, current training database, fold 4): randomly initialized prototypes trained with Proxy Anchor, compared with linear and multilayer-perceptron classifier heads at two loss weights.

**Table S4. Hyperparameters of ExPert v0.1.0**

Settings of the released model, trained on all 22 datasets with no held-out context. Only settings that act in this configuration are listed: loss terms at weight 0, modules that are built but never trained, and options of encoder, adapter or prior-map types that are not used are omitted. Every value was read from the run's resolved configuration and checkpoint, and checked against the training log: each listed loss weight held its value at every step, and every other loss weight was 0 throughout. The objective is the sum of the terms under Objective. Proxy Anchor, the prior pull and the zero-shot negatives act on the perturbation prototypes. The prior pull moves only the prototypes with the least training evidence toward a prediction from their prior neighbors, and it never reaches the encoder directly. The efficiency head reads a detached latent, so it does not shape the latent. Zero-shot fill, the prototypes written into the released model for the 10,931 untrained vocabulary genes after training. Learned parameters count the encoder, decoder, efficiency head and the trained prototype rows. Columns: section, parameter, value, configuration key (encoder., decoder. and cls_emb. denote the encoder, decoder and prototype-table keyword arguments; anneal., the loss-weight schedules) and notes.

**Table S5. Benchmark training databases and their held-out-context folds, related to Figure 2**

(A) The two databases. The 200-perturbation database is a subset of the full training database: each dataset contributes at least 30% of the 200 perturbations, chosen greedily by the number of datasets measuring them, with random tie-breaks and fill (seed 42). The 1,157-perturbation database (ctx7) keeps every perturbation measured in at least 7 of the 10 cell contexts. 128 perturbations are in both. The databases hold 671,242 and 2,741,678 cells over the same 22 datasets and 10 contexts (the 12 Marson2025 donor × condition datasets are one context, primary T cells). Each database has 12 folds (seed 42). In-distribution validation and test sets each take at most 10 cells per perturbation and dataset from the training contexts. The out-of-distribution test set is capped the same way in ctx7 and uncapped in the 200-perturbation database. The ctx7 sets are the full database's sets restricted to its perturbations, so no ctx7 evaluation cell was used to train a full-database model.

(B) Cells and perturbations per dataset in each database, with the dataset's context.

(C) Folds. Each fold holds out one donor together with one cell line, one cell line, or one T cell condition (Rest, Stim8hr, Stim48hr); the out-of-distribution test set is every cell of the held-out datasets (subject to the cap in A), and no cell of these datasets is used for training. Columns: held-out context(s) and datasets; cells and perturbations in the training, in-distribution validation, in-distribution test and out-of-distribution test sets. Fold numbers do not name the same held-out contexts in the two databases (for example, fold 3 holds out D1 + HCT116 in the 200-perturbation database and D1 + TeloHAEC in ctx7).

(D) Every perturbation in either database: membership in each, and the number of contexts and datasets measuring it in ctx7.

**Table S6. Zero-shot recovery per screen**

Values behind **Figure 3B**. One row per fold model (n = 12), context and split: ID, the context's cell line or donors were in training; OOD, the fold model held the context out. Each query is the pseudobulk of up to 25 cells of one perturbation in one screen, ranked against all 19,895 prototypes. "All unseen perturbations" pools the contexts; "all seen perturbations" gives the trained perturbations of the same split for reference. Columns: unique perturbations and queries; top-10 recall weighted by knockdown effect size and unweighted, each as a fraction and relative to chance (× chance; chance = 10/19,895); top-10 hits; and the score with the zero-shot prototypes shuffled across perturbations (shuffled null; mean and SD over the shuffles). Over all unseen perturbations, the median over fold models is 26.2× (ID) and 13.5× (OOD).

**Table S7. Prior screen**

Zero-shot score: top-10 recall of the unseen perturbations, each query weighted by its knockdown effect size, relative to chance (× chance; 9,874 unseen perturbations measured in a screen, ranked against all 19,895 prototypes). Each prior set places the unseen perturbations by the shipped graph fill ($\cos^4$-weighted mean of the k = 25 nearest trained prior neighbors).

(A) All 48 single priors: family (text, gene descriptions written by a language model and embedded by a text encoder; STRING protein–protein interactions (PPI)[51,52]; DepMap[46,47] co-dependency; proteome; morphology; L1000; genomic position), KEGG neighbor lift (share of a gene's 10 prior neighbors that share a KEGG pathway with it, relative to random pairs, size-matched), zero-shot score, top-10 hits, and the score with the prior shuffled across genes (shuffled null).

(B) All 1,023 subsets of the ten best single priors (at most three text priors; the shipped text prior ranks fifth among single priors and is not in this pool): members and their families, size, zero-shot score over all unseen perturbations and on each half of a random split of the unseen genes, the score of the same subset with each prior shuffled across genes (shuffled-gene twin), number of text priors, whether a source family appears twice, rank, and whether the subset scores above the production prior set (text, DepMap and STRING; 25.53×, scored separately on the same screen). 425 subsets score above it, 407 of which (96%) contain two versions of one source (two text priors, 222; three, 102); the best subset with one prior per source scores 28.37× (Opus 4.7 text, DepMap and STRING experimental interactions), and the production set would rank 19th of these 143 subsets.

(C) Production text prior: gene descriptions written by Claude Opus 4.8 (shipped; single prior 21.79×, rank 5 of 48) or Opus 4.7 (22.87×, rank 1), both embedded with qwen3-embed-8b, as single priors (zero-shot score, rank and KEGG neighbor lift) and combined with DepMap and STRING (zero-shot score and top-10 hits: 25.53×, 84 hits vs 25.34×, 83 hits).

**Table S8**. Drug-target references for sci-Plex3: the HDAC machinery gene set, the class I corepressor subunits it was calibrated on, and vendor-datasheet targets, related to **Figure 4**

(A) The 48 genes against which **Figure 4D** scores each HDAC inhibitor condition: the members of the curated SIN3–HDAC1/2, CoREST, NuRD and NCoR–HDAC3 complexes and the catalytic subunits HDAC1–3 (38 genes), together with the CORUM-core partners of the HDAC family (27 genes, 17 of them also curated members). A CORUM-core partner shares CORUM complexes of at most 20 members with an HDAC target gene, scored $S_g = n_g/\sqrt{(N_T \cdot D_g)}$ ($n_g$, complexes containing the gene; $N_T$ = 67, all such complexes; $D_g$, all CORUM complexes containing the gene) and kept at $S_g \geq 0.15$ among the sci-Plex3 candidates. Columns: complexes, curated membership, partner status and $S_g$, and whether the gene is among the 3,802 candidates shared by all methods (34 of 48; the remaining 14 cannot be ranked).

(B) The 35 subunits of the four class I HDAC corepressor complexes (the curated complexes without the catalytic HDAC1–3), used to set the partner size limit and threshold: complex, and whether the partner rule recovers the subunit (with its $S_g$). The rule recovers 17 of the 35 subunits and 17 of the 21 subunits among the candidates; 17 of its 27 partners are subunits.

(C) Targets of the 30 sci-Plex3 drugs without a usable ChEMBL target, from vendor datasheets (Selleck Chemicals; Cayman Chemical for pirarubicin). Every protein in the vendor's targets, IC50 or Ki table was counted, excluding entries reported inactive and non-protein readouts; class-level entries were expanded to all members of the class. Columns: drug, ChEMBL ID and target, vendor source, vendor target entries, entries used and excluded, class expansion, target genes used, and the references cited on the datasheet (with PMID and DOI).

**Table S9. Tahoe-100M drug misses whose ExPert ranking matches the drug's known pharmacology, related to Figure 4**

One row per drug whose annotated target ExPert did not rank in the top ten (a miss) but whose ranking is explained by the drug's biology or by an annotation or sample problem (**Supplementary Note 5**). Columns: drug; Tahoe-100M mechanism of action and annotated targets; ExPert evidence (ranks among the 8,964 trained prototypes in the loss or activation direction; drug-class similarity as z against 2,000 random drug sets of the same size from other plates); the same evidence from plain gene expression, without the model; interpretation; literature source; and whether the drug is counted among the 13 explained misses. Elimusertib is listed but not counted: its wells behave like the MEK inhibitor TAK-733 in the neighboring well, a likely sample mix-up.

**Table S10. Pathway enrichment in alcohol-associated liver disease hepatocytes, and the curated gene sets**

Values behind **Figure 5E** and **5F**. Each part lists every non-redundant pathway with nominal $p < 0.05$ in alcohol-associated liver disease (ALD; alcohol-associated cirrhosis and alcohol-associated hepatitis combined) hepatocytes, ranked by $p$; shown_in_figure marks the 15 rows drawn in the figure. Gene sets: Reactome 2024 pathways with 10–500 members, plus curated mRNA export and mRNA splicing sets (A only). One-sided hypergeometric test in the direction with the smaller $p$; $q_{BH}$, Benjamini–Hochberg adjusted over all gene sets and both directions tested. Viral and disease-named pathways are excluded; a pathway whose hit genes overlap a higher-ranked pathway by ≥50% of the smaller hit set is folded into it (folded_sets). Columns: gene set and source; direction; set size; hits, expected hits and fold enrichment (hits/expected); $\log_2$ fold enrichment as plotted ((hits + 0.5)/(expected + 0.5)); $p$; $q_{BH}$; hit genes; and folded sets.

(A) ExPert: the union of the top 200 predicted perturbations of AC and AH hepatocytes, in each direction (predicted loss or gain of function); background, the 8,964 perturbations in the ExPert bank. $healthy_{hits}$ and $healthy_p$ give the same gene set and direction in healthy hepatocytes. 28 pathways.

(B) Differential expression: the union of the top 200 downregulated and upregulated genes (pool-level pseudobulk, ranked by Wald statistic) of AC and AH hepatocytes versus healthy; background, all genes tested. 41 pathways.

(C) The curated gene sets behind (A) and (B): every set tested that is not a plain Reactome 2024 pathway, and the literature rows of **Figure 5E**. The grouped mRNA-export set and its four parts (nuclear pore, TREX / TREX-2 and export adaptors, export receptors and the Ran cycle, Reactome mature-mRNA transport); the mRNA-splicing set (Reactome major and minor spliceosome pathways plus every SR protein); the readout modules, defined by gene-name patterns before any scoring; and each literature row with the pathway or module it maps to and its reference. Columns: gene set, label in **Figure 5E**, source, definition, number of member genes among the 8,964 trained prototypes, reference, and the member genes. The other 1,227 gene sets tested are Reactome 2024 pathways with 10–500 member genes among the trained prototypes (Enrichr library Reactome_Pathways_2024), listed by name in (A) and (B) when enriched.

**Table S11. Donors of the PBMC samples, related to Figure 6**

One row per donor of the single-cell RNA-seq PBMC samples (n = 9): donor number (PT in **Figure 6**), group (primary mitochondrial disease or control) and detailed group, likely affected gene and detailed genetics, sex, age at sample collection (years) and clinical phenotype. **Figure 6** treats the four primary mitochondrial disease donors (PT43, PT46, PT54, PT67) as PMD, the two TBCK donors (PT41, PT42) as non-PMD, and the remaining controls (PT38, PT50, PT55) as undiagnosed.
